# Needler: An Algorithm to Develop a Comprehensive Targeted MS Method Capable of Monitoring the Human Proteome

**DOI:** 10.1101/2022.01.25.477578

**Authors:** Damien B. Ready, Gregory K. Potts

## Abstract

Targeted proteomics for addressing biological hypotheses through quantitative mass spectrometry (MS) is increasingly popular because of its ability to deliver sensitive and reproducible results to inform protein concentration and dynamics. While the technique is powerful, the associated tooling to develop unique targeted data acquisition methods per experiment is limited, often focused on producing a minimally viable analytical approach for a limited set of analytes, rather than one which can maximally quantitate all targets of interest. We developed the Needler algorithm to produce comprehensive targeted MS methods capable of quantitating the maximum number of proteins given available instrument constraints. Using this algorithm, we demonstrate that the minimum instrument parameters required to produced targeted MS quantitation of all tryptic peptides from the human proteome are well in excess of current capabilities. Given these constraints, we provide a visualized scope of the boundaries for practical MS monitoring to maximize proteome and subproteome coverage dependent upon instrument duty cycle capabilities.

## Introduction

Quantitative liquid chromatography tandem mass spectrometry (LC-MS/MS) proteomics is a standard analytical approach to robustly monitor peptides, metabolites, lipids, and other biomolecules from a complex sample matrix.^1^ The peptides produced from proteolytic digestion of a source protein are first identified in a data-dependent acquisition (DDA) discovery MS analysis and used as the basis for development of a targeted MS acquisition by monitoring specific peptide precursor mass to charge ratios (m/z) and instrument-generated fragment ions over specific retention time windows.^2^ The gold standard technique for MS quantitation is selected reaction monitoring (SRM) or multiple reaction monitoring (MRM) performed on a triple quadrupole instrument with scheduled targeting windows. These methods can analyze a limited set of peptide precursors and are often used for high accuracy quantification measurements for a subset of the proteome dictated by the experimental aims. Parallel reaction monitoring (PRM) is a more recent development that utilizes a high-resolution instrument such as an Orbitrap mass spectrometer to simultaneously monitor all peptide fragment ions.^3^ For each of these approaches, the quantifiable peptides must be carefully chosen to maximize utilization of instrument time without compromising the quantitative integrity of the assay. Data acquisition methods that attempt to monitor too many peptides may capture fewer data points per peptide precursor and risk limiting instrument sensitivity. Consequentially, the analytes of interest may become undetectable or quantitative accuracy may be sacrificed. Scheduled targeted MS methods that monitor only where the peptide is known to chromatographically elute risk missing the detection of the peptide if the elution gradient experiences an unexpected shift in retention time.

Despite the breadth of publicly available MS resources, such as protein databases, *in silico* protein processing tools, and peptide retention time libraries, building a targeted method from scratch to monitor a known list of proteins’ proteotypic peptides can be very time consuming. Constructing a scheduled MS acquisition instrument method is still a manual task requiring human judgment for what peptides to monitor when faced with multiple available candidates. Despite available resources which can aid the analyst,^4–6^ these existing tools are typically algorithms which are focused on producing a minimally sufficient method rather than maximally utilizing the available duty cycle of the instrument. Current mass spectrometry instruments and standard methods typically monitor in the 10s to 100s of peptides per run,^7^ greatly limiting the number of proteins that could be quantified for addressing broader proteomics questions in biology.

Given the limited time frame in which peptides chromatographically elute and the available instrument duty cycle, a frequent question arises: what is the best subset of peptides to monitor to quantitate the greatest number of proteins per experiment? To address this need, we developed the Needler algorithm, which is a system capable of constructing a targeted MS method to maximally utilize the available duty cycle of an instrument over the targetable LC gradient. Using this algorithm, we further endeavored to ask: what are the required performance characteristics of a MS instrument and LC system required to produce a comprehensive MS method capable of monitoring the entire proteome and tens of thousands of peptides? Towards this end, we applied Needler to the human proteome and demonstrate its capability to build large (> 10,000 analytes) targeted MS methods that can maximally utilize available instrument capacity. The results of the analysis indicate that existing instrumentation and techniques are incapable of monitoring the full proteome without significant advances in instrumental design and construction. Additional use cases highlight Needler calculations for the MS targeting capacity for proteins from the liver proteome or human kinase panels which demonstrate that subproteomes are more readily achievable for comprehensive MS targeting, depending on instrument duty cycle and the number of targeted analytes. Finally, the visualized scope of MS targeting boundaries provides a resource for determining the feasibility of various target peptide list sizes.

## Methods and Materials

### Peptide Database

The Uniprot SwissProt 2020_03 release^8^ was downloaded and reduced to the 20,368 canonical Human entries. Using Pyteomics,^9^ the proteins were digested *in silico* with trypsin allowing for no missed cleavages, size-restricted to 5-30 amino acids in length, and filtered to only consider peptides with the 20 canonical amino acids. Peptides that could not be distinctly assigned to a single protein were discarded. Methionine containing sequences were eliminated to avoid oxidation variability. All proteins were required to be identified by at least two peptides. The final collection of 431,811 peptides sequences represented 19,339 proteins (94.9% of initial proteome). The retained proteome peptide distribution is shown in Figure 1. The liver tissue peptide subset was created by the same procedure, limited to the proteins reported in^10^ with greater than zero protein copies per cell for liver tissues, covering 9,731 proteins with 260,700 peptides. The kinase peptide subset was created by the same procedure, limited to the proteins reported in the Uniprot list of human protein kinases, retaining 511 proteins with 18,194 peptide sequences.

**Figure 1:**
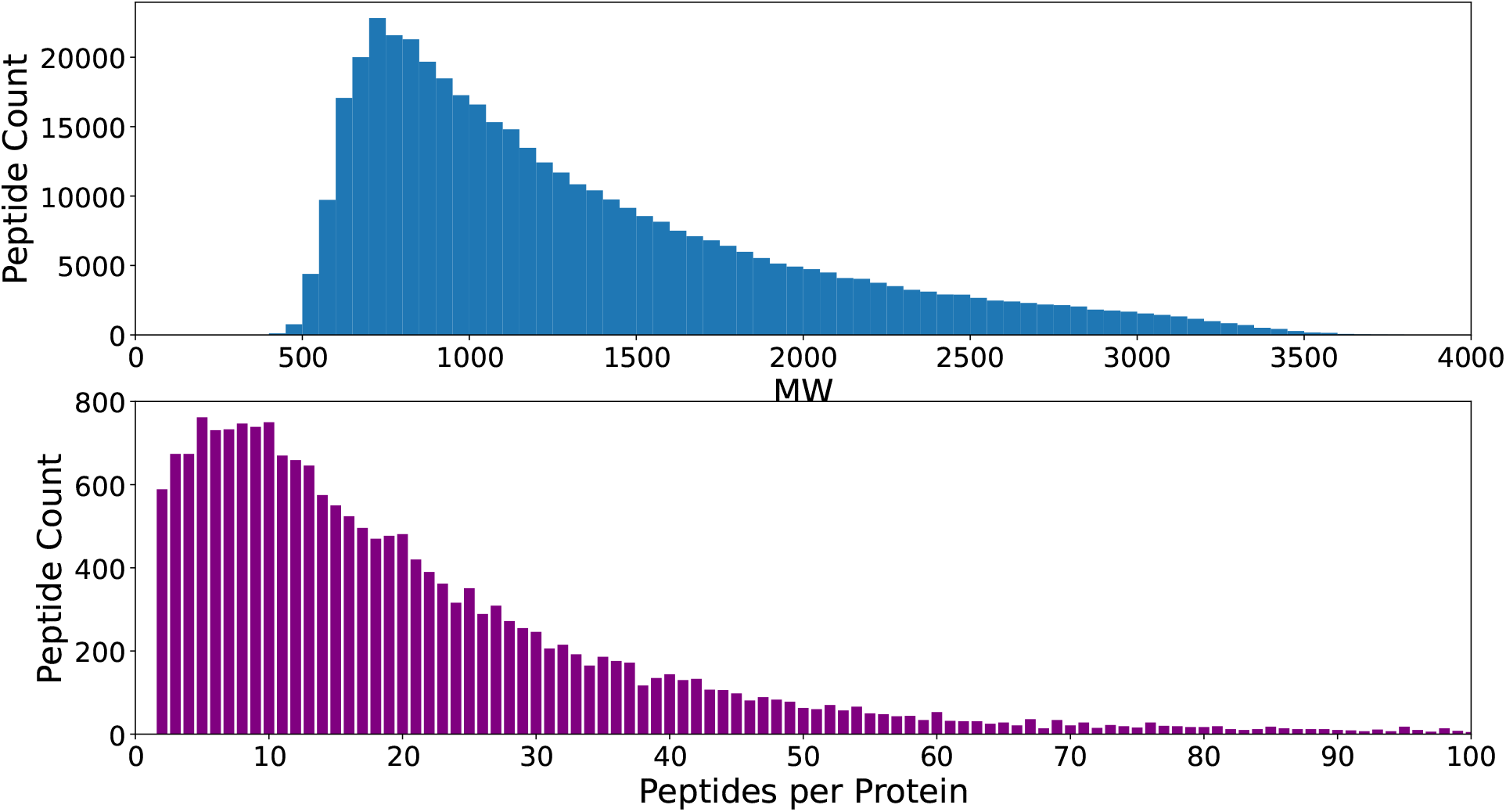
Top: Molecular weight distribution of the retained 431,811 tryptic peptides after filtering. Bottom: Distribution of peptides per protein that were retained after filtering.

### Peptide Retention Time Assignment

The PROSIT^11^ deep learning model was downloaded locally (commit `ce93f3aaed49aab4 628223b9025dcaf0512ea767`) and used to predict the indexed retention time (iRT) for all peptides retained after filtering. To increase throughput, the PROSIT Flask application was modified to predict only the iRT value and not infer the fragment ion intensities. The PROSIT predicted iRT values were converted to a 60 minute gradient equivalent by linear regression against the empirically observed retention times of the 40 PROCAL^12^ peptides using SciPy,^13^ Pandas,^14^ and Matplotlib^15^ (Supplementary Table 1). Predicted peptide apex times were rounded to whole second integer values. The linear fit and the distribution of peptide retention times are shown in Figure 2.

**Figure 2:**
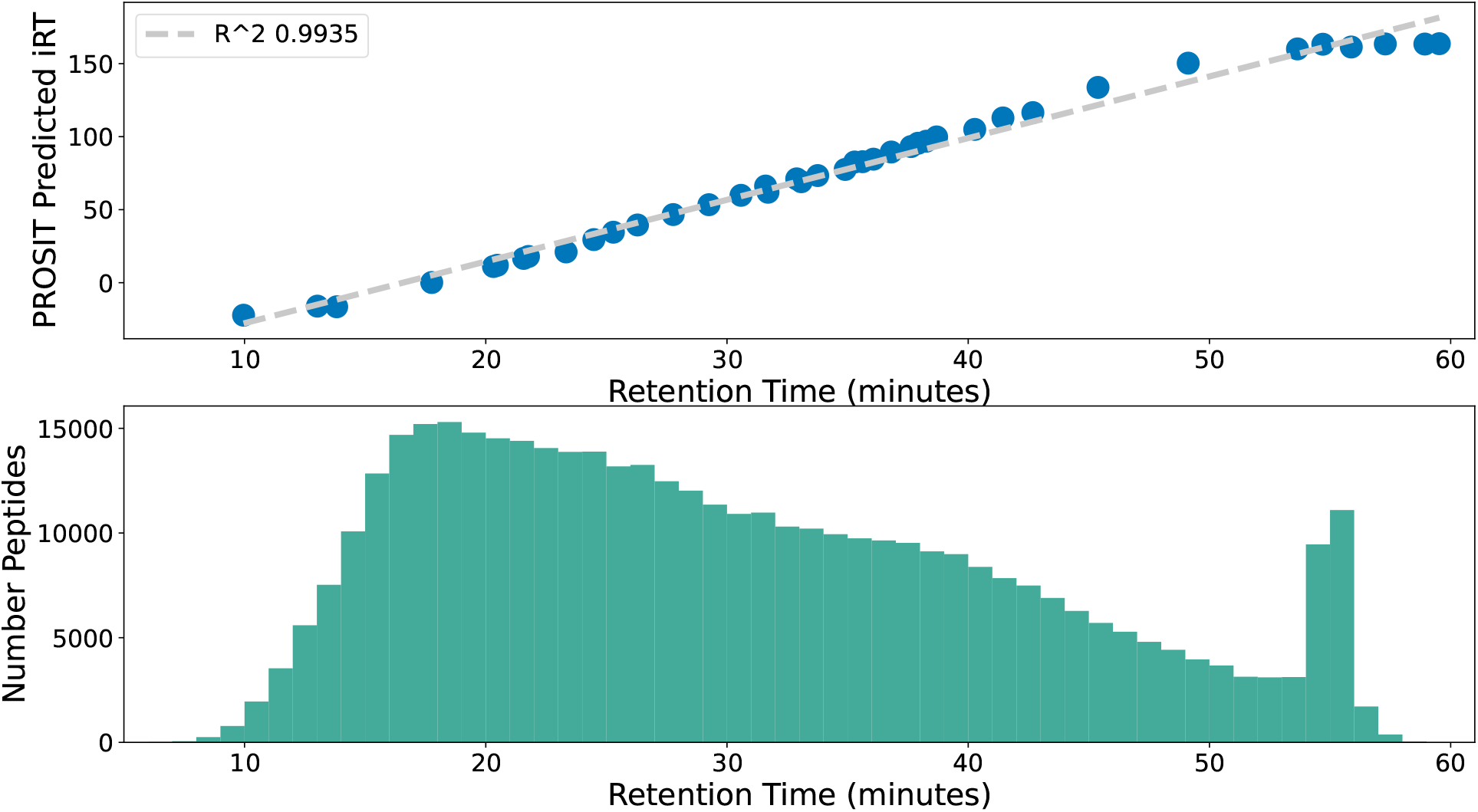
Top: Empirical PROCAL retention time fit against the predicted Prosit iRT with a R^2 of 0.9935. Bottom: distribution of predicted retention times of the proteome peptides used for modeling along a 60 minute gradient.

### Algorithm Overview

The Z3 Theorem Prover^16^ was used to host the peptide scheduling solver. A custom Z3 model was constructed to represent a scheduled targeted MS experiment with constraints on maximum targetable peptides per cycle where each peptide was considered from its predicted peak retention time plus the configured retention time width for the experimental configuration. The solver goal for each model was to identify the subset of peptides which would monitor the maximum number of proteins, requiring two peptides for a protein to be considered quantified. A matrix of instrument parameters was created to evaluate the effect of targets per cycle and the scheduled retention time targeting tolerance. The considered targets per cycle were: 1, 2, 3, 4, 5, 10, 20, 50, 100, 200, 500, 1000, 2000, 5000, and 10000. The retention time tolerances constructed by taking the peptide peak apex and monitoring the number of seconds before and after this point: 1, 3, 7, 15, 30, 45, 60, 90, 120, 180, 240, and 300. Peptides and proteins were randomly ordered at model construction to allow each replicate an opportunity to sample a different area of the solution space. Each model was constructed and then allowed to optimize for 24 hours on a single Cascade Lake processor core before being halted. Optimization jobs were distributed onto an 82 node cluster using Slurm. ^17^ Each model configuration was run three times, retaining the solution which targeted the maximum number of proteins. Full algorithm code is available on GitHub at https://github.com/dbready/needler.

## Results

The Needler algorithm solves a job scheduling problem given challenging constraints. Peptide elution times are not evenly distributed across the gradient and proteins have unequal numbers of candidate peptides available for targeting. A simplified example of the target selection problem is shown in Figure 3. The hypothetical instrument is capable of simultaneously monitoring two targets per cycle. There are three proteins (A, B, C), each with their own peptides (protein A has 3 available peptides, protein B has 4 available peptides, and protein C has 2 available peptides) which have distinct retention time targeting windows (darker shading indicates overlapping retention times). The task is to pick the subset of peptides which will yield the minimum required peptides per protein (two) without exceeding the capabilities of the MS instrument. Needler produces a solution to the scheduling problem by targeting peptides A1, A3, B1, B2, C1, and C2, allowing the instrument to monitor all three proteins of interest.

**Figure 3:**
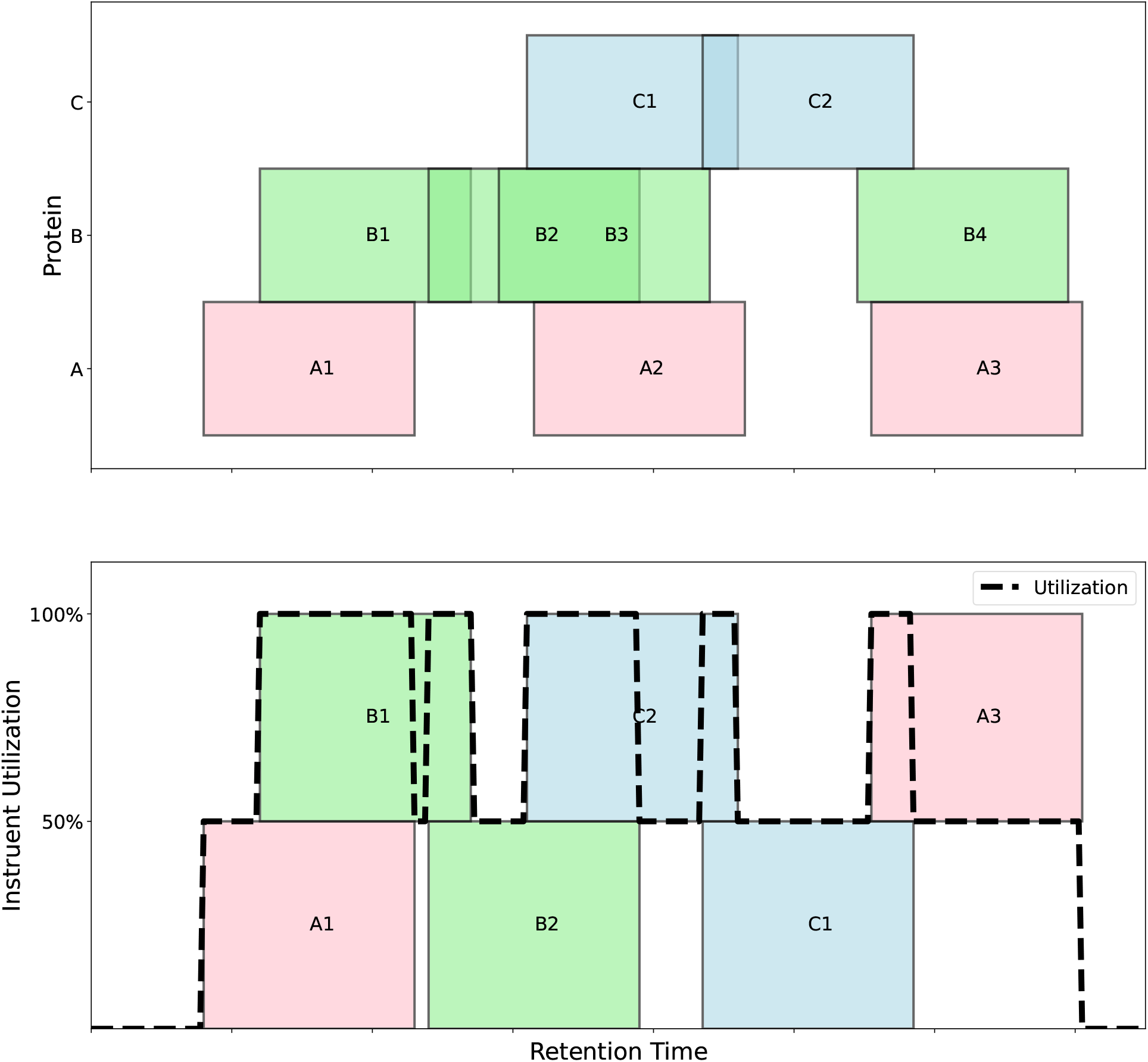
Schematic depicting the challenge in picking the subset of peptides for a targeted MS acquisition to monitor the maximum number of proteins. Top: candidate peptide retention time windows for each protein of interest Bottom: 6 Needler selected peptides resulting in quantiating all 3 proteins of interest.

An example acquisition method produced by Needler to develop a method for the human kinase panel, capable of monitoring 5 peptides per cycle, with each peptide monitored ± 180 seconds from the peak apex is shown in Figure 4. Each colored rectangle represents a different targeting window for a unique peptide chosen by Needler. In this acquisition method, the instrument would monitor 40 peptides, yielding quantitative data on 20 proteins. Despite the constraint of developing the method around the available peptide library, the method maintains high instrument utilization (gaps in utilization are indicated by non-colored areas of the graph). Additional selected utilization diagrams are provided in Supplemental Figures 3-74.

**Figure 4:**
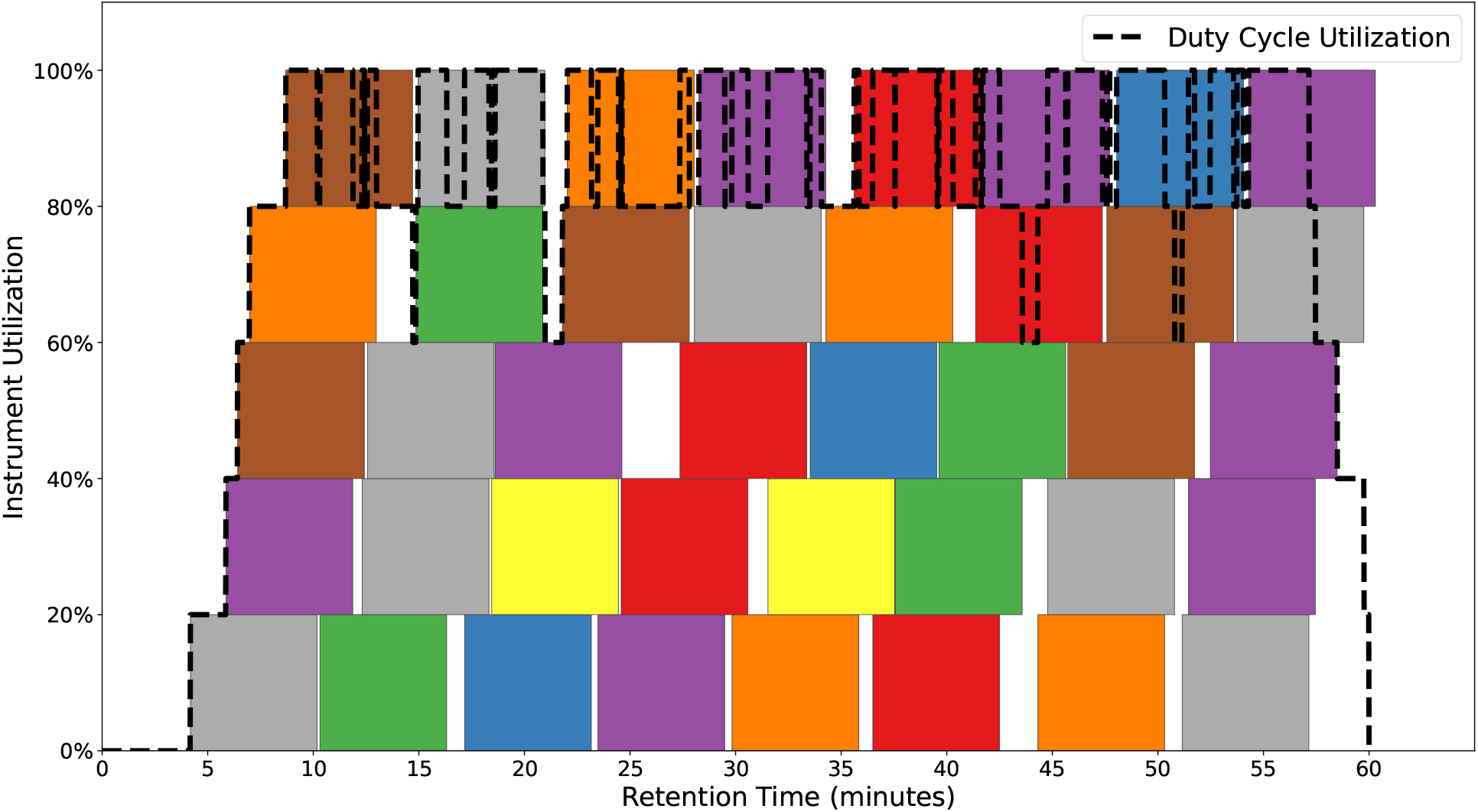
Visualization of a solved Needler MS acquisition method designed to target the human kinome proteins. Each colored block represents a scheduled targeting window per peptide. The dashed line indicates the total instrument utilization. MS acquisition method monitors 40 peptides, quantitating 20 proteins.

The best of the three models for each proteome, retention time tolerance, and targets per second combination were processed and the fraction of covered proteome was saved. The retained human proteome subset is shown in Figure 5. Monitoring more targets per second allows Needler additional flexibility in producing a comprehensive model, so solutions on the right part of the heatmap are better able to cover a high percentage of the proteome. Smaller retention time targeting windows allow the method to dedicate less time per peptide and are easier to solve. Given the human proteome with an instrument capable of monitoring 40 targets per second with scheduled retention time windows of ± 7 seconds from peak apex, Needler is able to target 19% of the available proteome: 7338 peptides representing 3669 proteins. The Needler algorithm was able to comprehensively target the human proteome in 39 of the tested models. To highlight additional uses case for Needler, we utilized the algorithm to examine the targeting capacity for protein subsets of the proteome, namely those proteins reported as being expressed in either human liver and a human kinase panel. Human liver proteome results are shown in Figure 6. Given the human liver proteome with an instrument capable of monitoring 40 targets per second with scheduled retention time windows of ± 7 seconds from peak apex, Needler is able to target 37.5% of the available proteome: 7290 peptides representing 3645 proteins. The Needler algorithm was able to comprehensively target the human liver proteome in 52 of the tested models. Finally, the optimized methods for targeting the human kinases are shown in Figure 7. Given the human kinase subproteome with an instrument capable of monitoring 40 targets per second with scheduled retention time windows of ± 7 seconds from peak apex, Needler is able to target 100% of the available protein panel: 1022 peptides representing 511 proteins. The Needler algorithm was able to comprehensively target the human kinase subproteome in 107 of the tested models.

**Figure 5:**
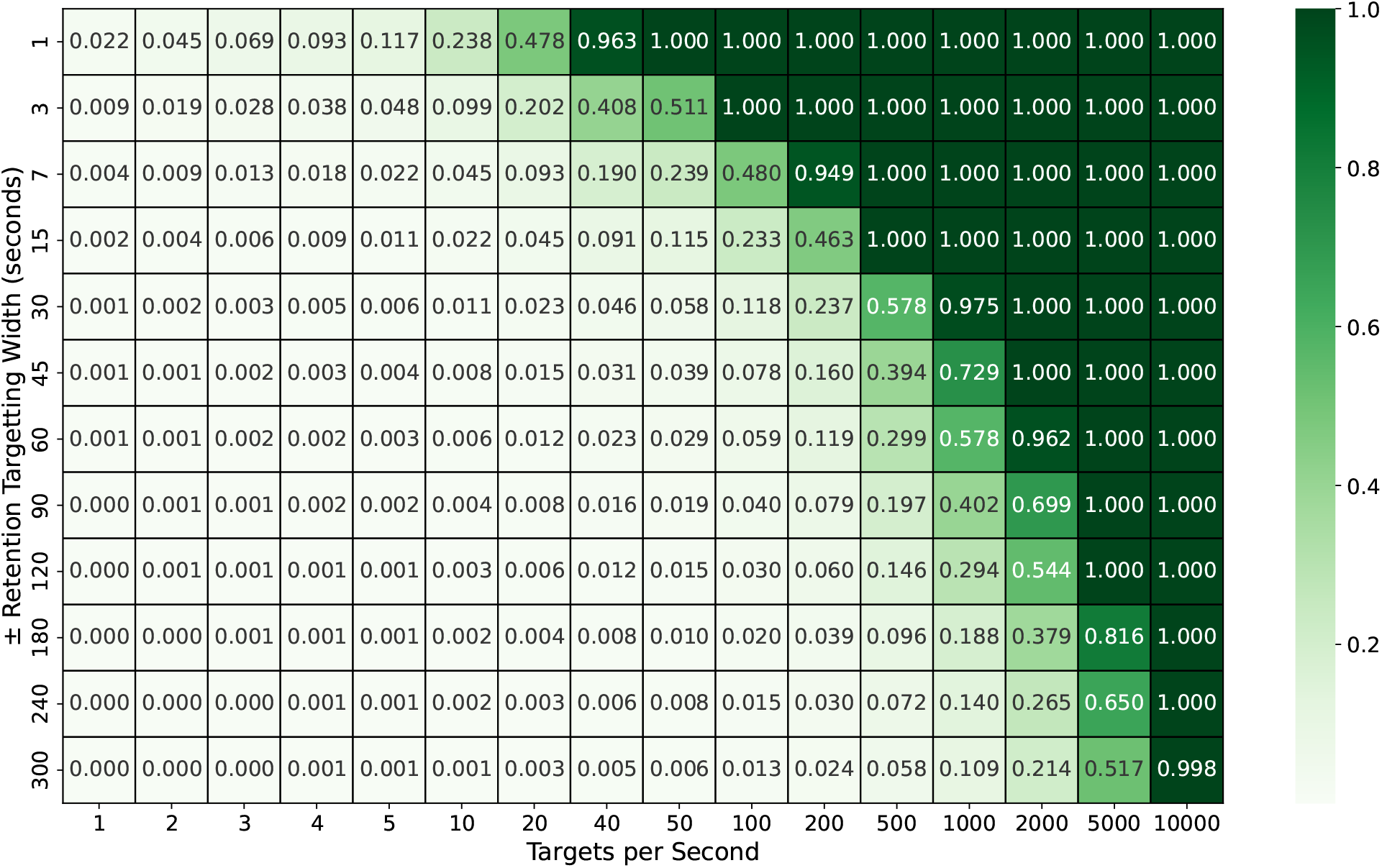
Resulting data matrix from building optimized MS methods attempting to target the 19,339 available human proteins. Every square in the matrix represents the fraction of targetable proteome identified by the algorithm.

**Figure 6:**
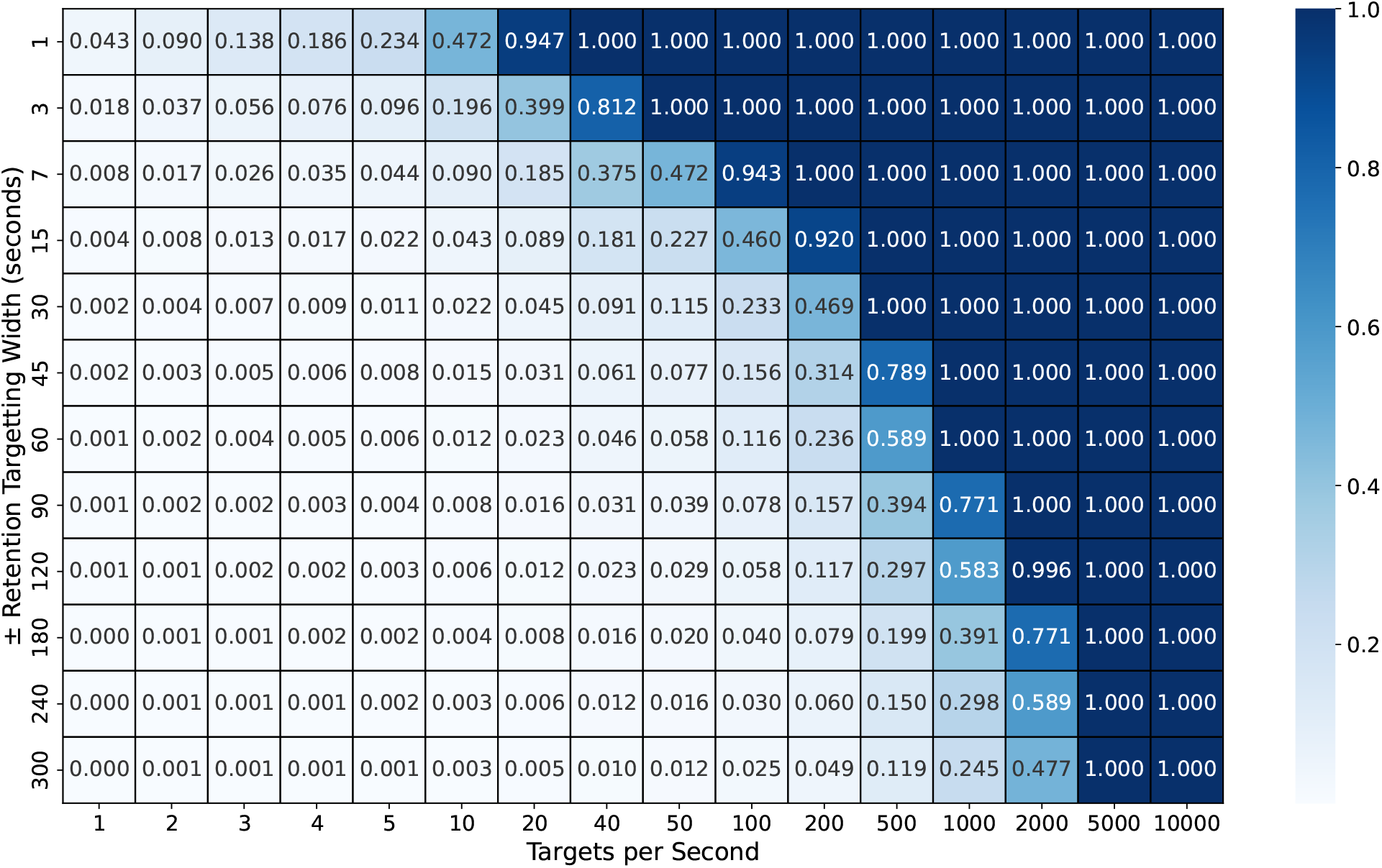
Resulting data matrix from building optimized MS methods attempting to target the 9,731 available proteins in the liver subset. Every square in the matrix represents the fraction of targetable proteome identified by the algorithm.

**Figure 7:**
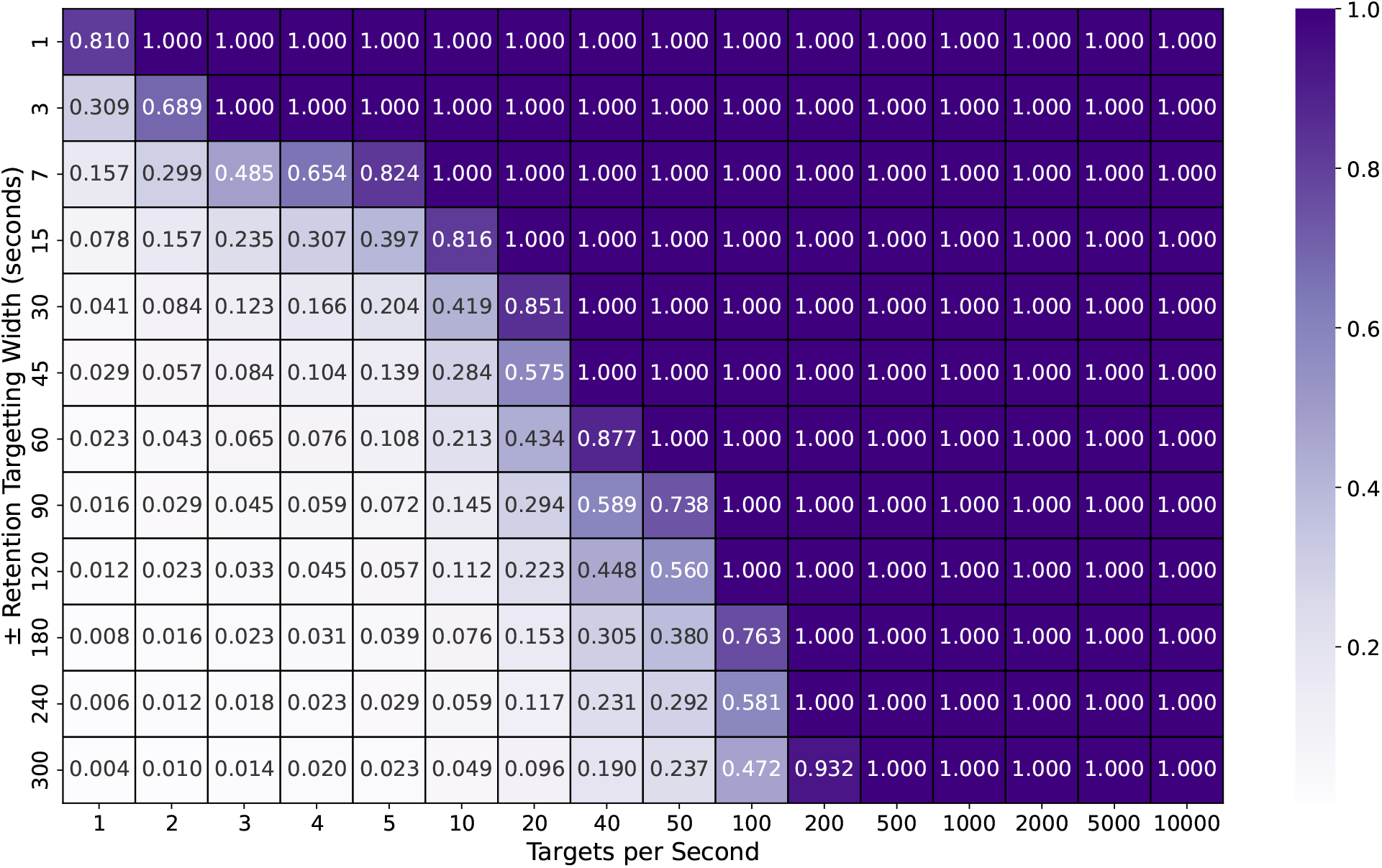
Resulting data matrix from building optimized MS methods attempting to target the 511 available proteins kinase proteins subset. Every square in the matrix represents the fraction of targetable proteome identified by the algorithm.

## Discussion

### Peptide Database

The criteria used for filtering the initial peptide database were conservative, so a real-world analysis might exclude some of the peptides considered here. For instance, peptides are often discarded when they are highly charged, exhibit poor precursor ionization efficiency, contain multiple modification sites, or generate weak fragment ionization transitions. While these factors must be considered when designing an actual experiment, we believe the results generated with the Needler algorithm are valid given expected retention times and increasing sensitivities of mass spectrometers to detect peptides with a range of characteristics. Ultimately the final curated list of peptides submitted to Needler for scheduling optimization could easily be fit to the analyst’s desired level of conservatism for peptide quantitation, but the final duty cycle utilization optimization would proceed as is reported here.

### Algorithm

The Z3 constraint solver has previously been used in operations research to tackle similar job scheduling problems,^18^ for which we thought peptide detection/MS duty cycle optimization contained parallels for consideration. Picking the subset of available proteotypic peptides from a protein can be considered a variation of the nurse scheduling problem, which is known to be NP complete.^19^ Computational complexities of modeling the full proteome model is an intractable problem, so we cannot guarantee an optimal model. It is possible that a superior modeling approach may be available that can better utilize available instrument duty cycle. Nevertheless, we consider the Needler algorithm and results to be an important step for raising awareness of the current capabilities and limitations for full proteome quantitation using existing mass spectrometry methods and instrumentation.

### Proteome Coverage

Defining the minimum acceptable quantitation criteria as 15 detection points across the chromatographic peak (± 7 seconds from peak apex), without additional retention time monitoring, the Needler algorithm calculated that the human and liver proteome would be fully monitorable with an instrument capable of operating at 500 hertz. On the other hand, the 511 identified human kinases would be fully quantifiable with an instrument operating at 40 hertz, which is within range of current MS instrumentation. For instance, the Thermo Exploris™ 480 Mass Spectrometer is considered state of the art, and when operating in PRM mode, it is capable of achieving 40 hertz. This indicates that subproteomes of ~500 proteins are within reach of modern equipment. However, the commonly observed irreproducibility of liquid chromatography means that the minimum acceptable scheduled targeting windows are much larger than a 15 data point calculation would allow. There are techniques and configurations thought to relieve this issue, but none fully resolve the concern to have wider retention time targeting windows.^20–22^ The generated methods with limited retention time coverage are unlikely to produce satisfactory results as peptides often shift during elution, escaping sufficient monitoring over their full profile.

MaxQuant Live^23^ which allows for dynamic target scheduling and reactive target selection could address this chromatographic limitation. MaxQuant Live was demonstrated to monitor 25,000 peptides through more specific instrument control that could adjust for changes in chromatography. However, the peptides were chosen with a greedy algorithm that did not maximize the number of monitored proteins, leaving an opportunity for this parameter to be further optimized.

While Needler offers a path to maximize the number of peptide analytes per analysis, considerable instrument performance improvements are required to comprehensively target and quantitatively measure the human proteome. Nevertheless, focused biological networks and protein families such as the human kinome are now feasibly targetable given existing practices and instrumentation. Needler offers an opportunity to pursue the greatest extent of protein quantification given available instrumentation. Lacking a revolution in LC-MS technology, more intelligent data acquisition modes seem the most likely means to capture a larger percentage of human biology.

## Conclusion

The Needler algorithm has demonstrated the ability to design a comprehensive scheduled targeted MS experiment within one’s available instrument constraints. We further demonstrate that existing instrumentation and techniques are not yet suited to the task of monitoring the complete human proteome. Despite the MS duty cycle limitations that exist, subprotomes from tissues or biological pathways of interest (hundreds of proteins per LC-MS run) are more readily achievable for comprehensive targeted MS with on-market instruments. Superior instrumentation or novel acquisition strategies that can dynamically react to the conditions of a chromatographic gradient will be required to quantitate a greater percentage of analytes.

Source code to the Needler algorithm used to support this publication is hosted on GitHub at https://github.com/dbready/needler under an Apache 2 license. Results for the best of three optimizations for each of the experiments is available in the provided SQLite database.

## Supporting information

SQLite Results Database

## Acknowledgement

We thank Melanie Patterson for helpful discussions and providing extensive manuscript improvements.

This work made use of the high performance computing resource that is operated by the National Center for Supercomputing Applications (NCSA) at the University of Illinois at Urbana-Champaign. We would like to thank the expertise of the staff at NCSA.

