## Supplementary Figures for "Needler: An Algorithm to Develop a Comprehensive Targeted MS Method Capable of Monitoring the Human Proteome"

### 1 Peptide Length Distribution

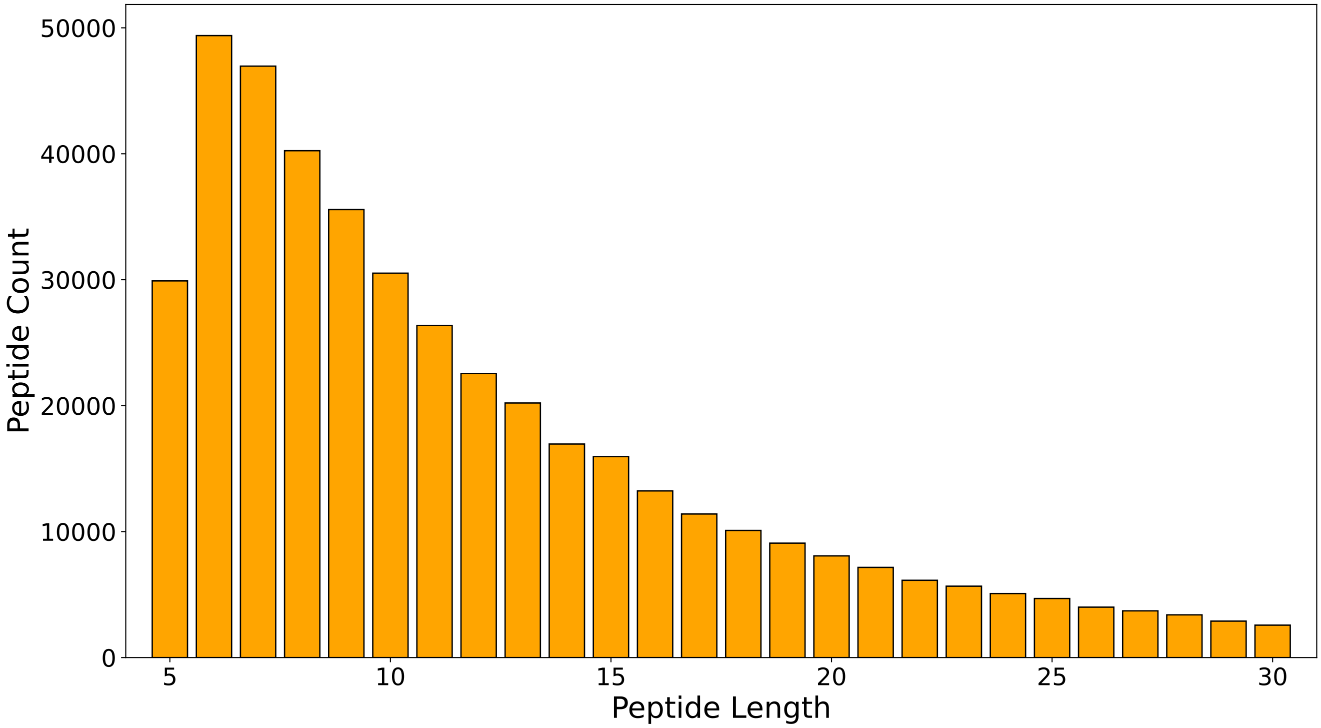

Supplementary Figure 1: Length distribution of all peptides retained in the full proteome dataset (431,811 sequences). Maximum limit of 30 residues was chosen to fit within the limitations of the PROSIT iRT predictor.

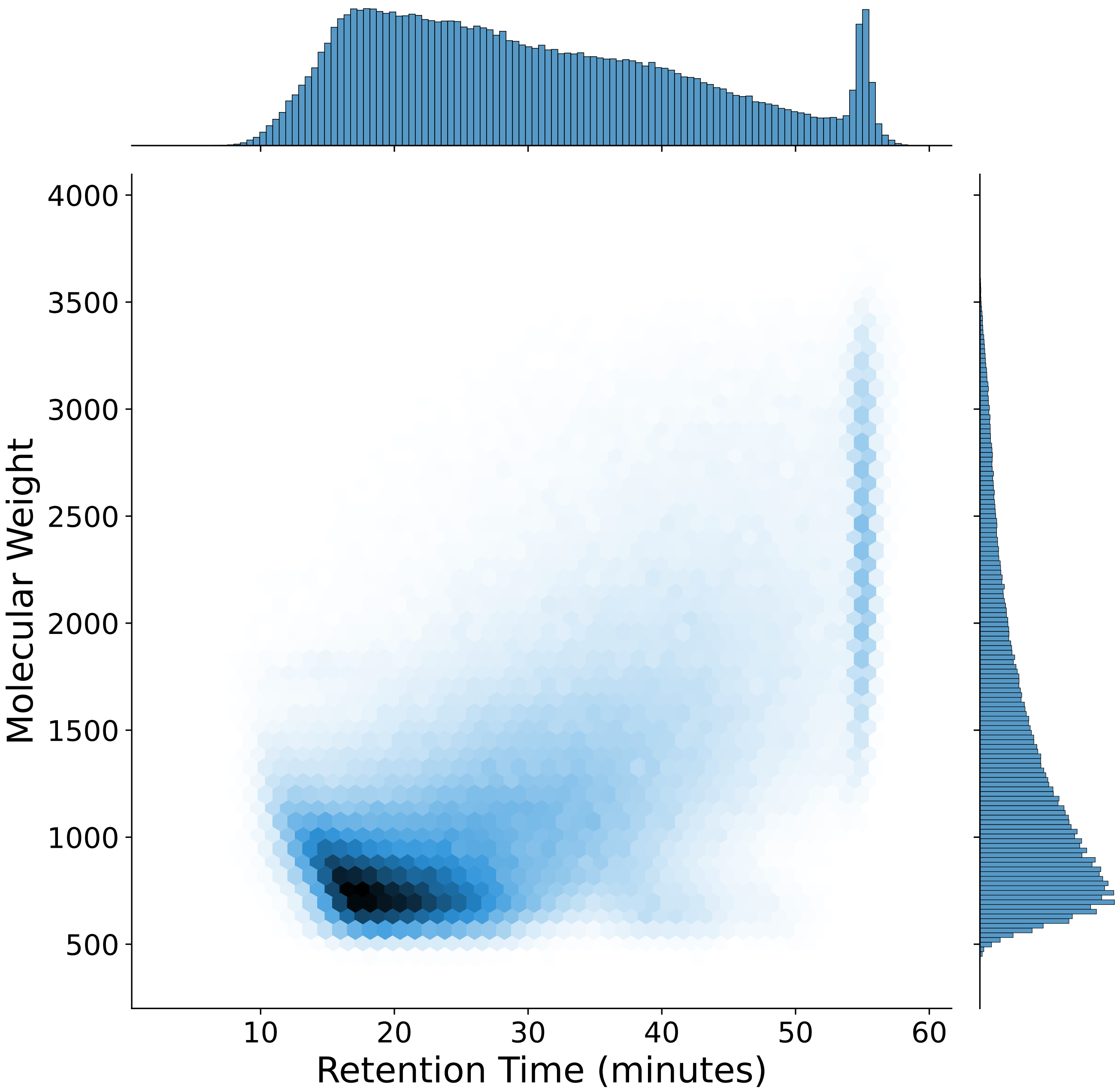

Supplementary Figure 2: Molecular weight vs 60 minute predicted retention time of the peptides contained in the full proteome dataset (431,811 sequences).

### 2 Selected Utilization Diagrams

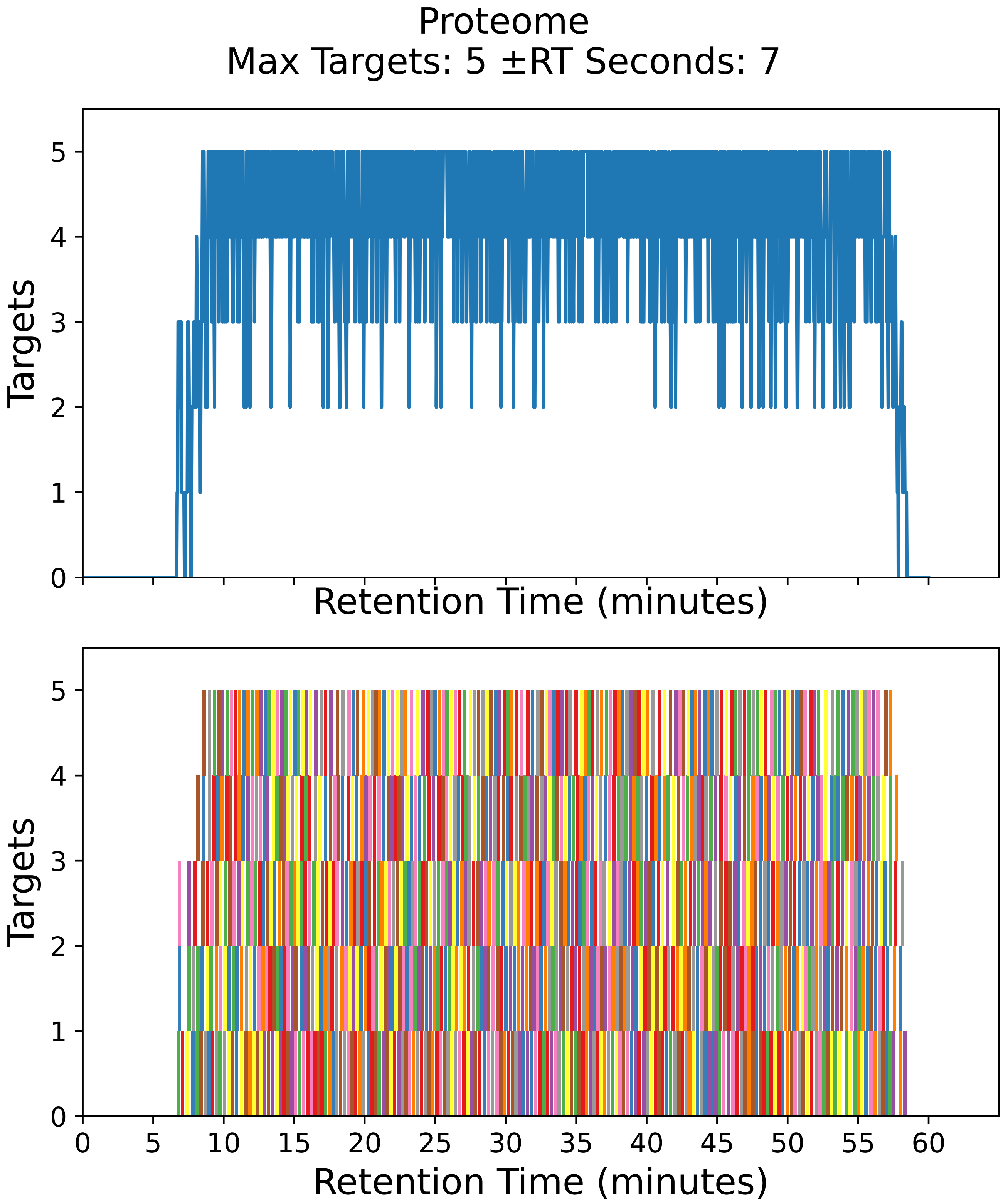

Supplementary Figure 3: Top: utilization diagram for the best fit method with instrument targeting the proteome filtered subset with a maximum of 5 targets per cycle with retention times ± 7 seconds from the peptide peak apex. Bottom: individual targeting windows for the 852 peptides, monitoring 426 proteins.

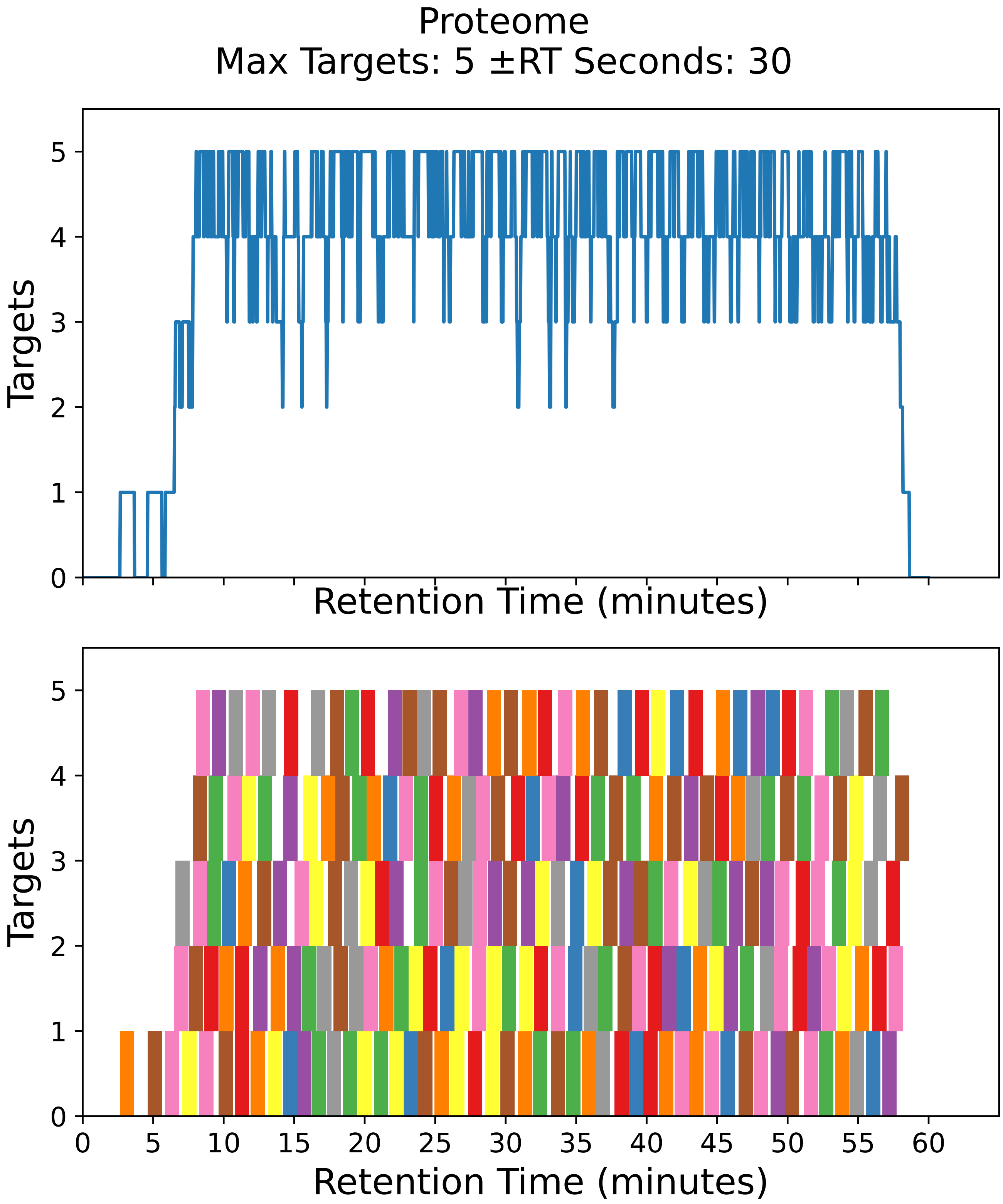

Supplementary Figure 4: Top: utilization diagram for the best fit method with instrument targeting the proteome filtered subset with a maximum of 5 targets per cycle with retention times ± 30 seconds from the peptide peak apex. Bottom: individual targeting windows for the 218 peptides, monitoring 109 proteins.

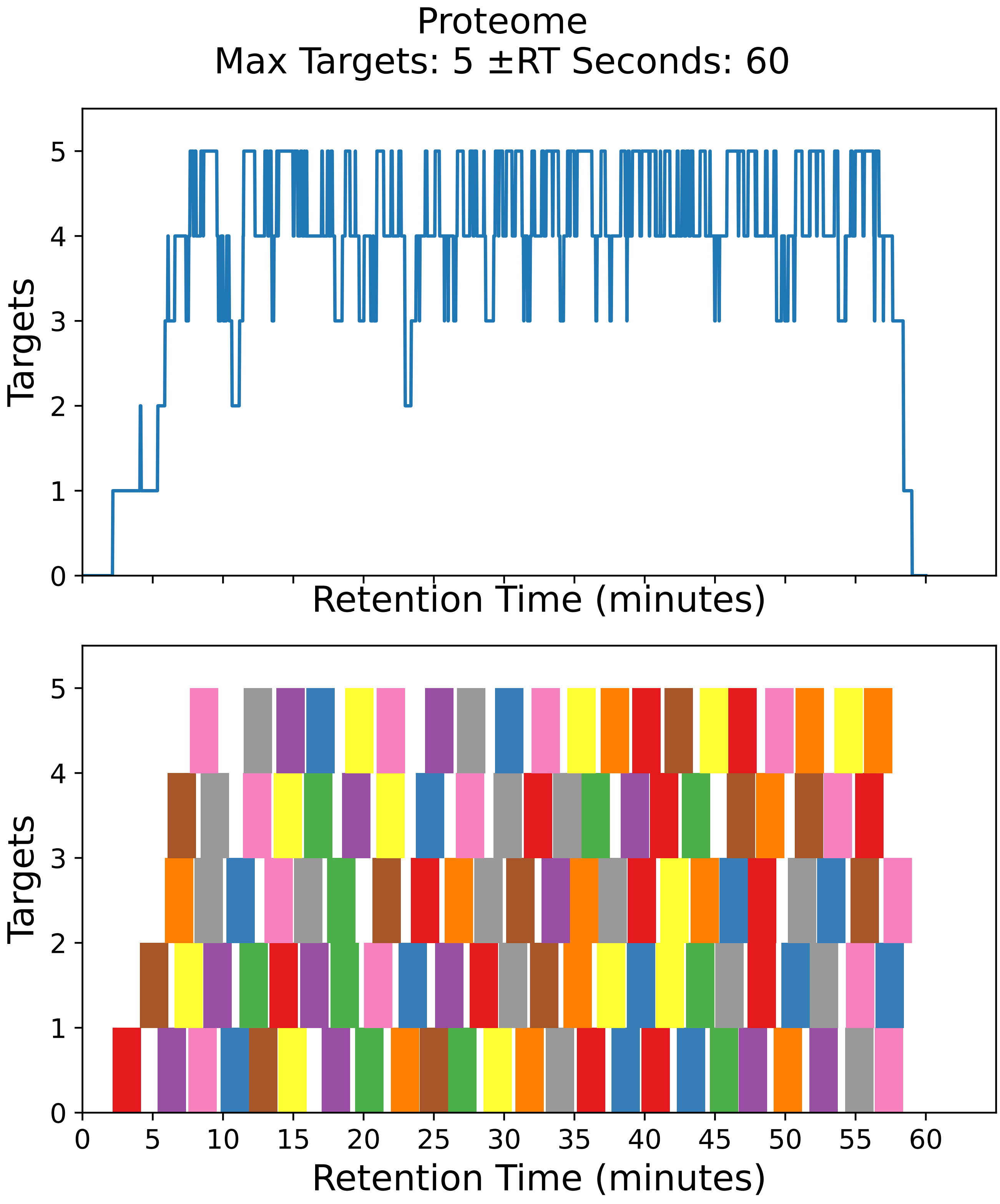

Supplementary Figure 5: Top: utilization diagram for the best fit method with instrument targeting the proteome filtered subset with a maximum of 5 targets per cycle with retention times ± 60 seconds from the peptide peak apex. Bottom: individual targeting windows for the 112 peptides, monitoring 56 proteins.

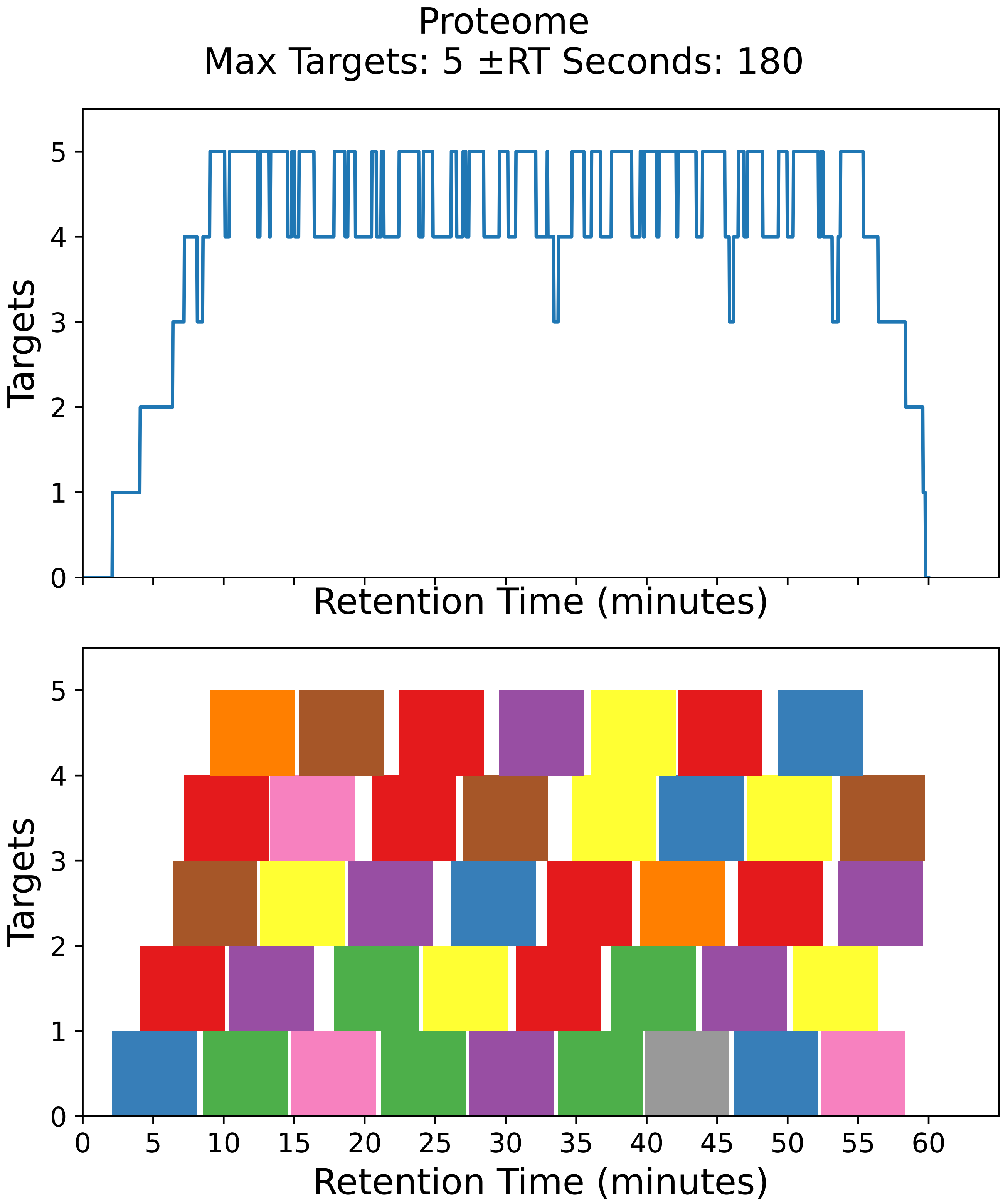

Supplementary Figure 6: Top: utilization diagram for the best fit method with instrument targeting the proteome filtered subset with a maximum of 5 targets per cycle with retention times ± 180 seconds from the peptide peak apex. Bottom: individual targeting windows for the 40 peptides, monitoring 20 proteins.

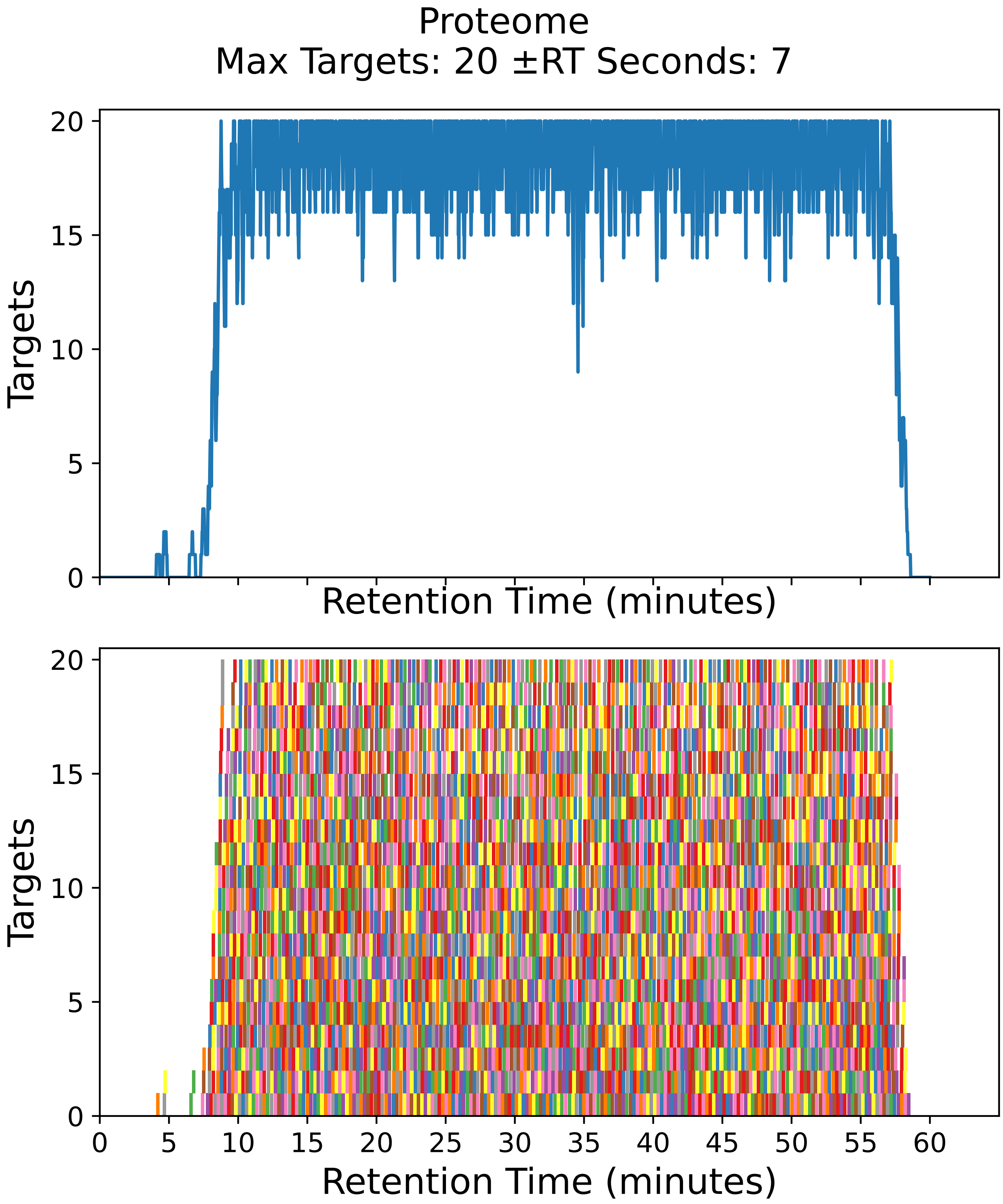

Supplementary Figure 7: Top: utilization diagram for the best fit method with instrument targeting the proteome filtered subset with a maximum of 20 targets per cycle with retention times ± 7 seconds from the peptide peak apex. Bottom: individual targeting windows for the 3592 peptides, monitoring 1796 proteins.

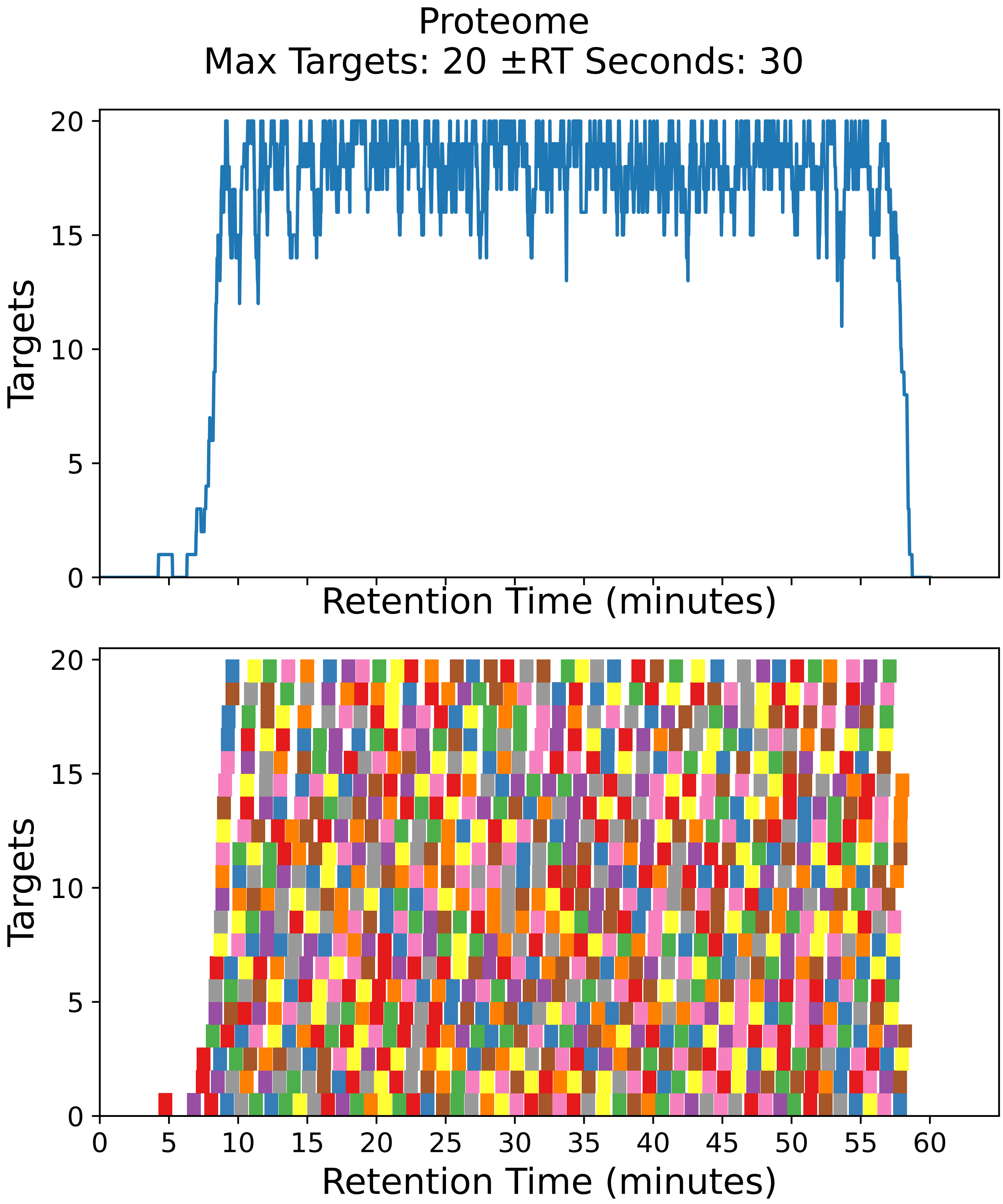

Supplementary Figure 8: Top: utilization diagram for the best fit method with instrument targeting the proteome filtered subset with a maximum of 20 targets per cycle with retention times ± 30 seconds from the peptide peak apex. Bottom: individual targeting windows for the 882 peptides, monitoring 441 proteins.

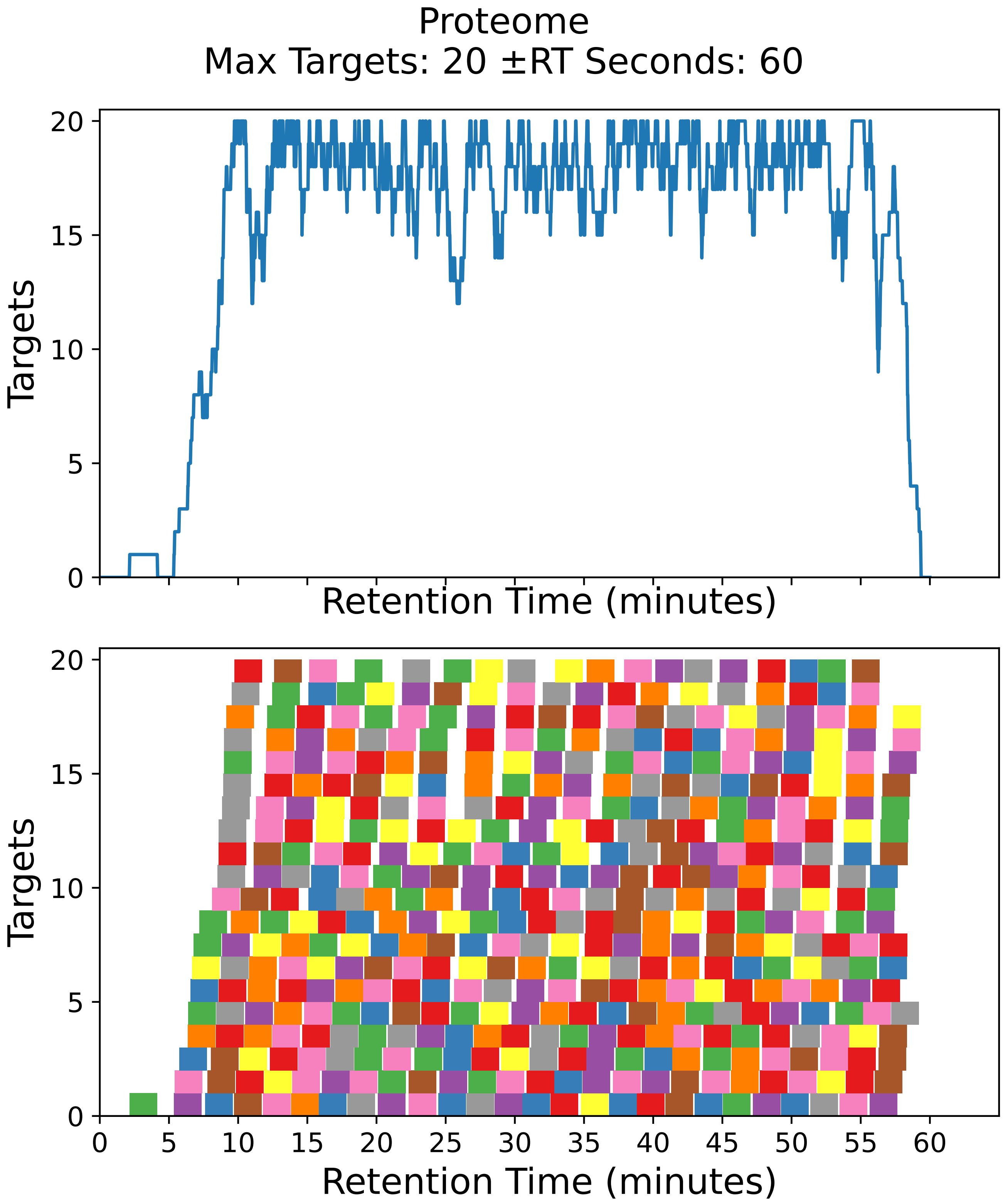

Supplementary Figure 9: Top: utilization diagram for the best fit method with instrument targeting the proteome filtered subset with a maximum of 20 targets per cycle with retention times ± 60 seconds from the peptide peak apex. Bottom: individual targeting windows for the 450 peptides, monitoring 225 proteins.

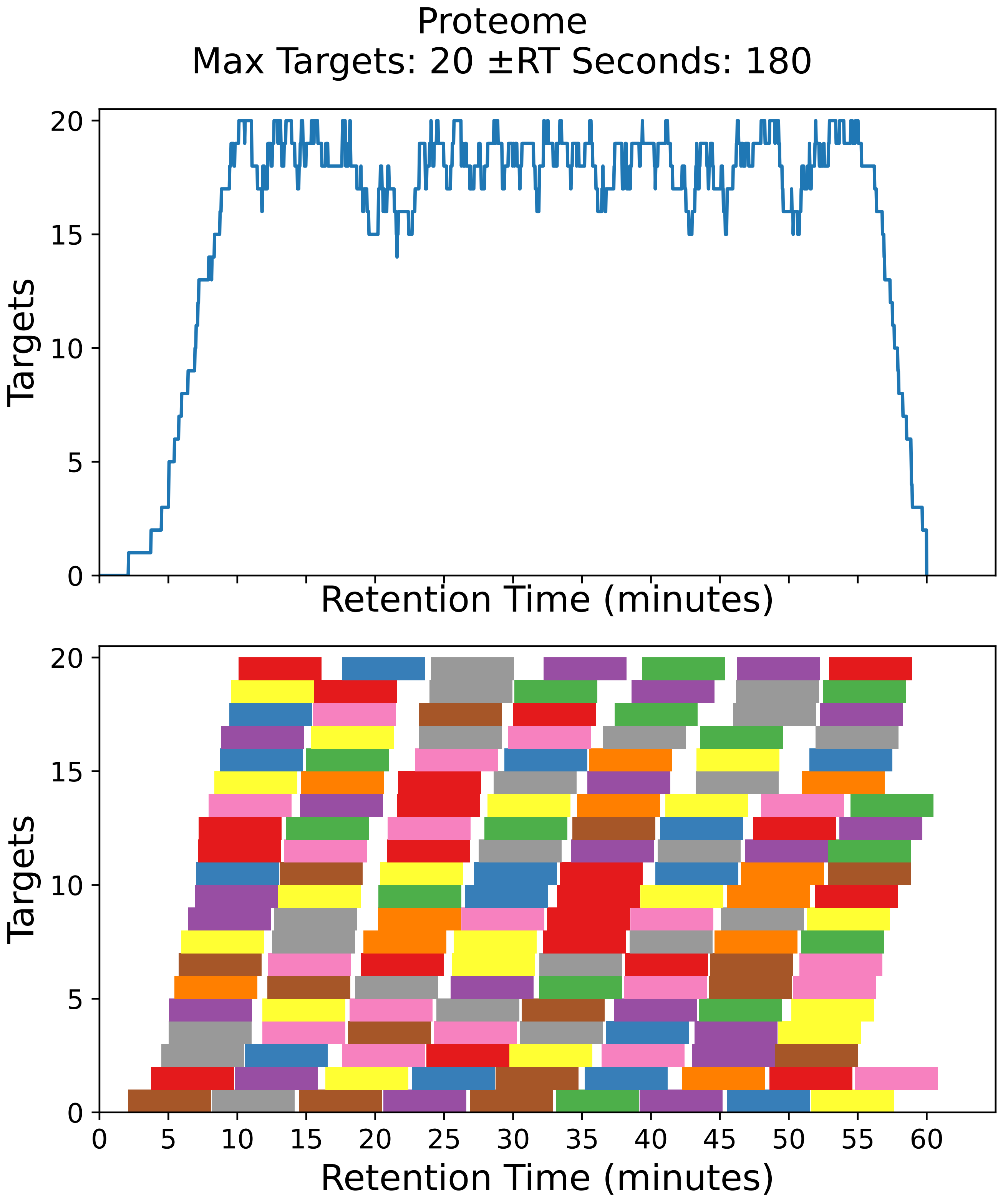

Supplementary Figure 10: Top: utilization diagram for the best fit method with instrument targeting the proteome filtered subset with a maximum of 20 targets per cycle with retention times ± 180 seconds from the peptide peak apex. Bottom: individual targeting windows for the 156 peptides, monitoring 78 proteins.

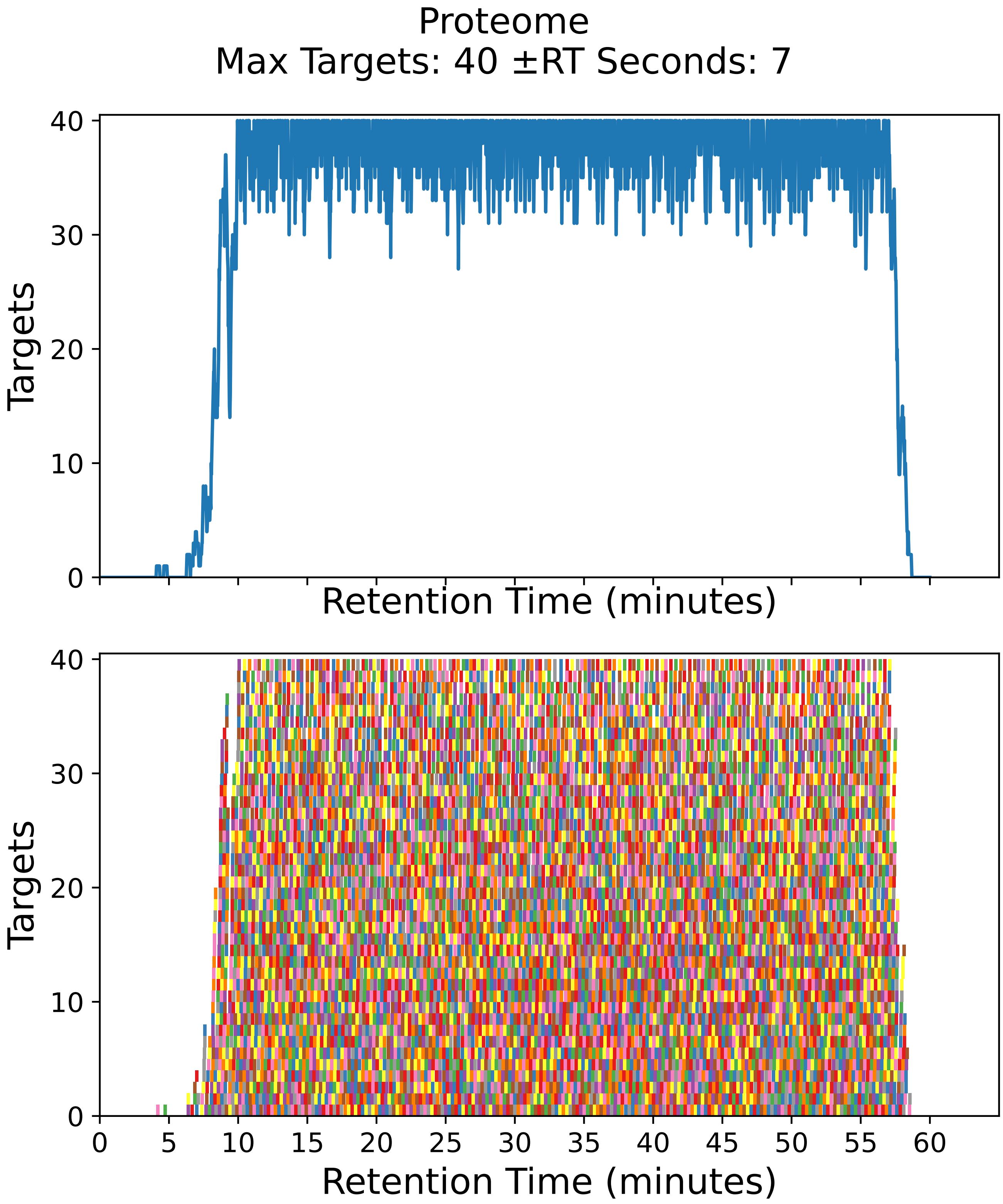

Supplementary Figure 11: Top: utilization diagram for the best fit method with instrument targeting the proteome filtered subset with a maximum of 40 targets per cycle with retention times ± 7 seconds from the peptide peak apex. Bottom: individual targeting windows for the 7338 peptides, monitoring 3669 proteins.

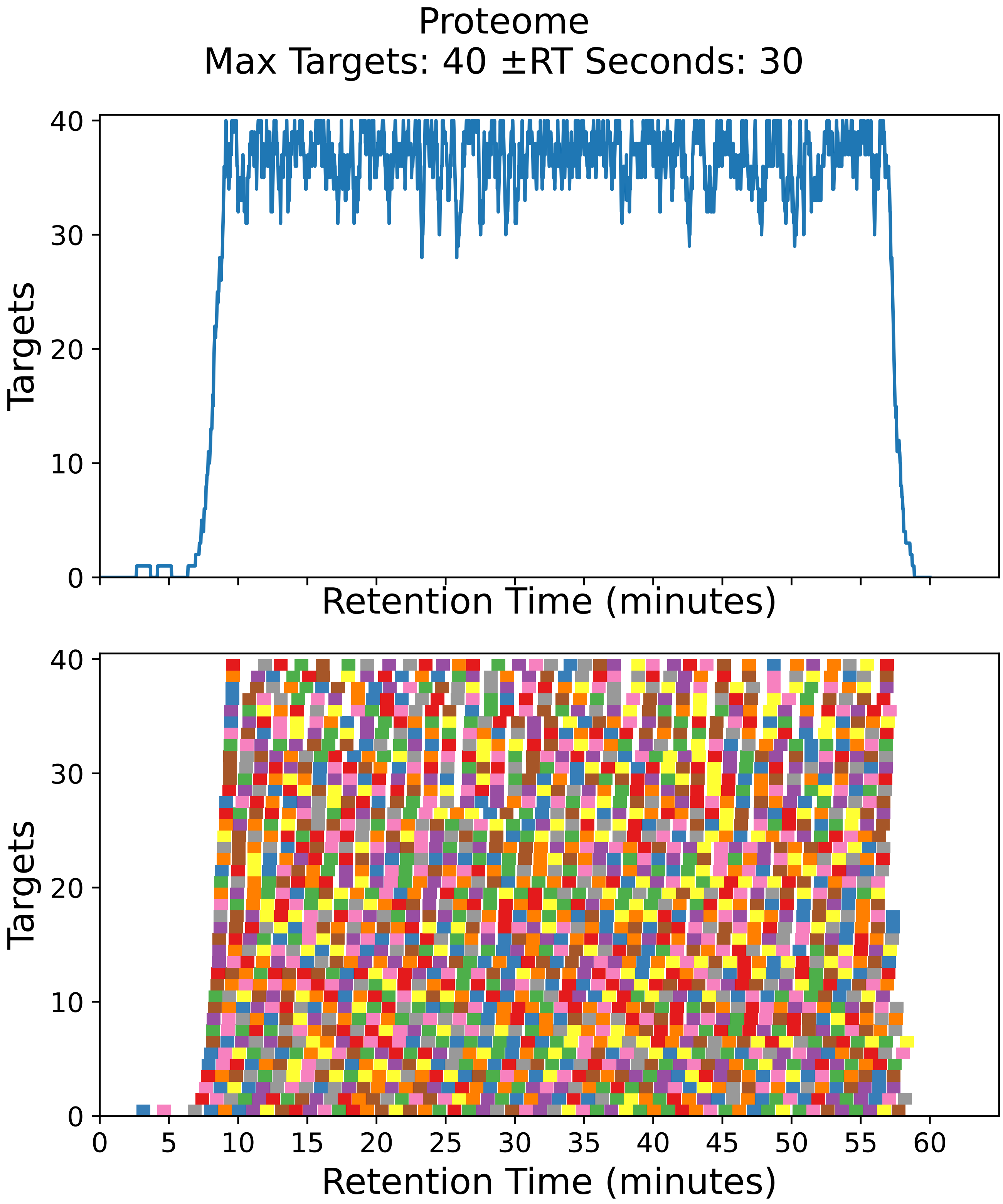

Supplementary Figure 12: Top: utilization diagram for the best fit method with instrument targeting the proteome filtered subset with a maximum of 40 targets per cycle with retention times ± 30 seconds from the peptide peak apex. Bottom: individual targeting windows for the 1786 peptides, monitoring 893 proteins.

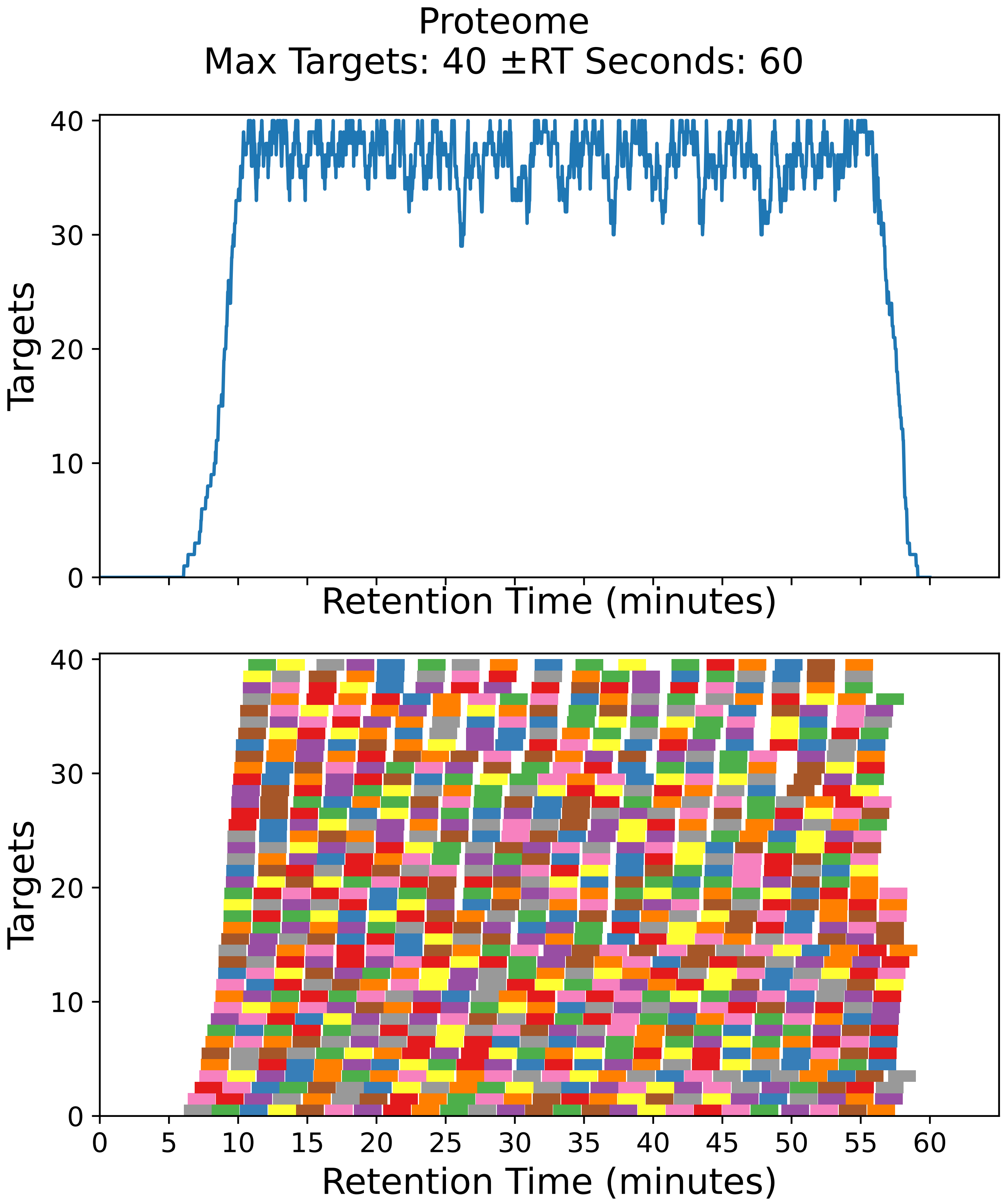

Supplementary Figure 13: Top: utilization diagram for the best fit method with instrument targeting the proteome filtered subset with a maximum of 40 targets per cycle with retention times ± 60 seconds from the peptide peak apex. Bottom: individual targeting windows for the 890 peptides, monitoring 445 proteins.

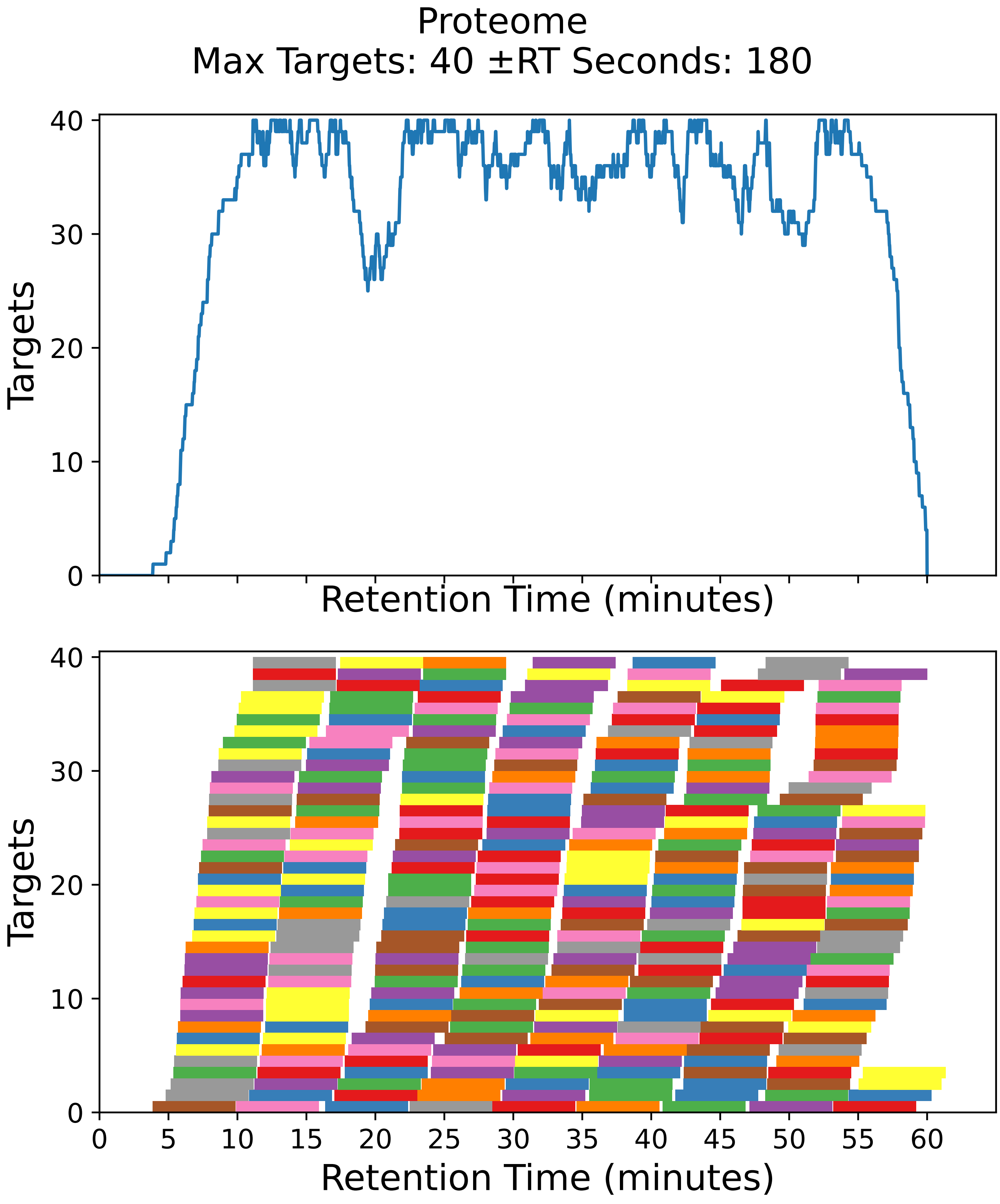

Supplementary Figure 14: Top: utilization diagram for the best fit method with instrument targeting the proteome filtered subset with a maximum of 40 targets per cycle with retention times ± 180 seconds from the peptide peak apex. Bottom: individual targeting windows for the 310 peptides, monitoring 155 proteins.

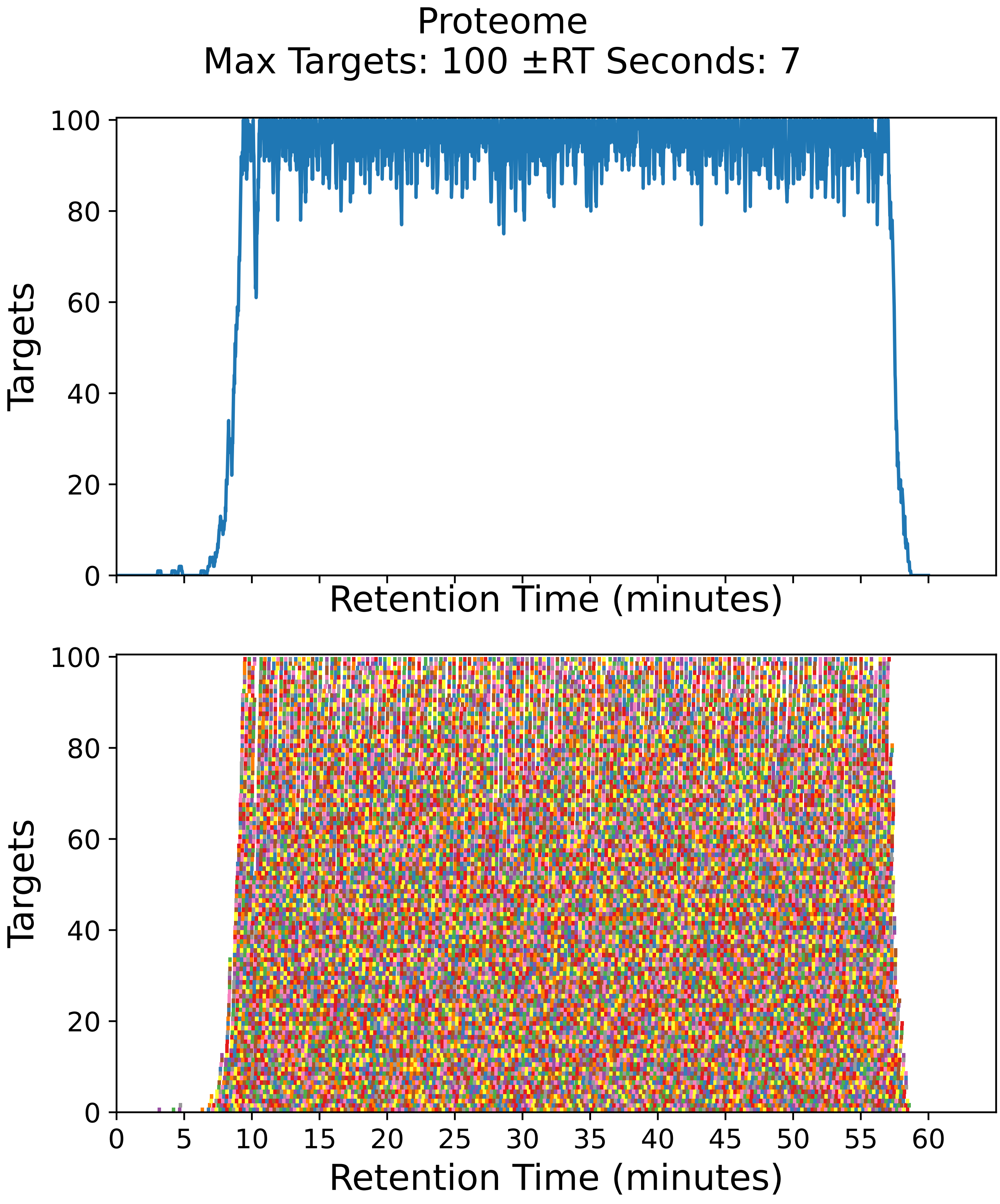

Supplementary Figure 15: Top: utilization diagram for the best fit method with instrument targeting the proteome filtered subset with a maximum of 100 targets per cycle with retention times ± 7 seconds from the peptide peak apex. Bottom: individual targeting windows for the 18572 peptides, monitoring 9286 proteins.

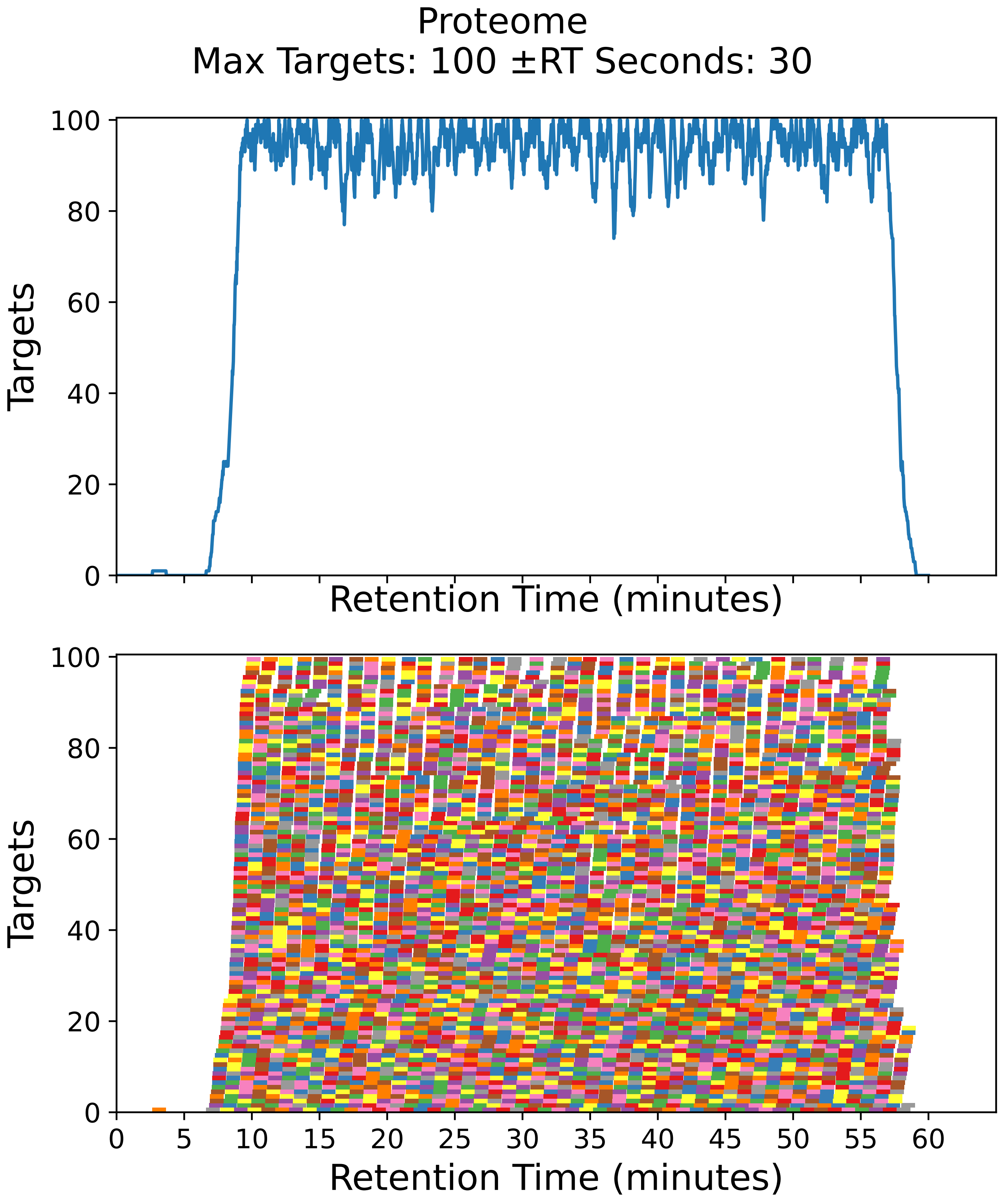

Supplementary Figure 16: Top: utilization diagram for the best fit method with instrument targeting the proteome filtered subset with a maximum of 100 targets per cycle with retention times ± 30 seconds from the peptide peak apex. Bottom: individual targeting windows for the 4566 peptides, monitoring 2283 proteins.

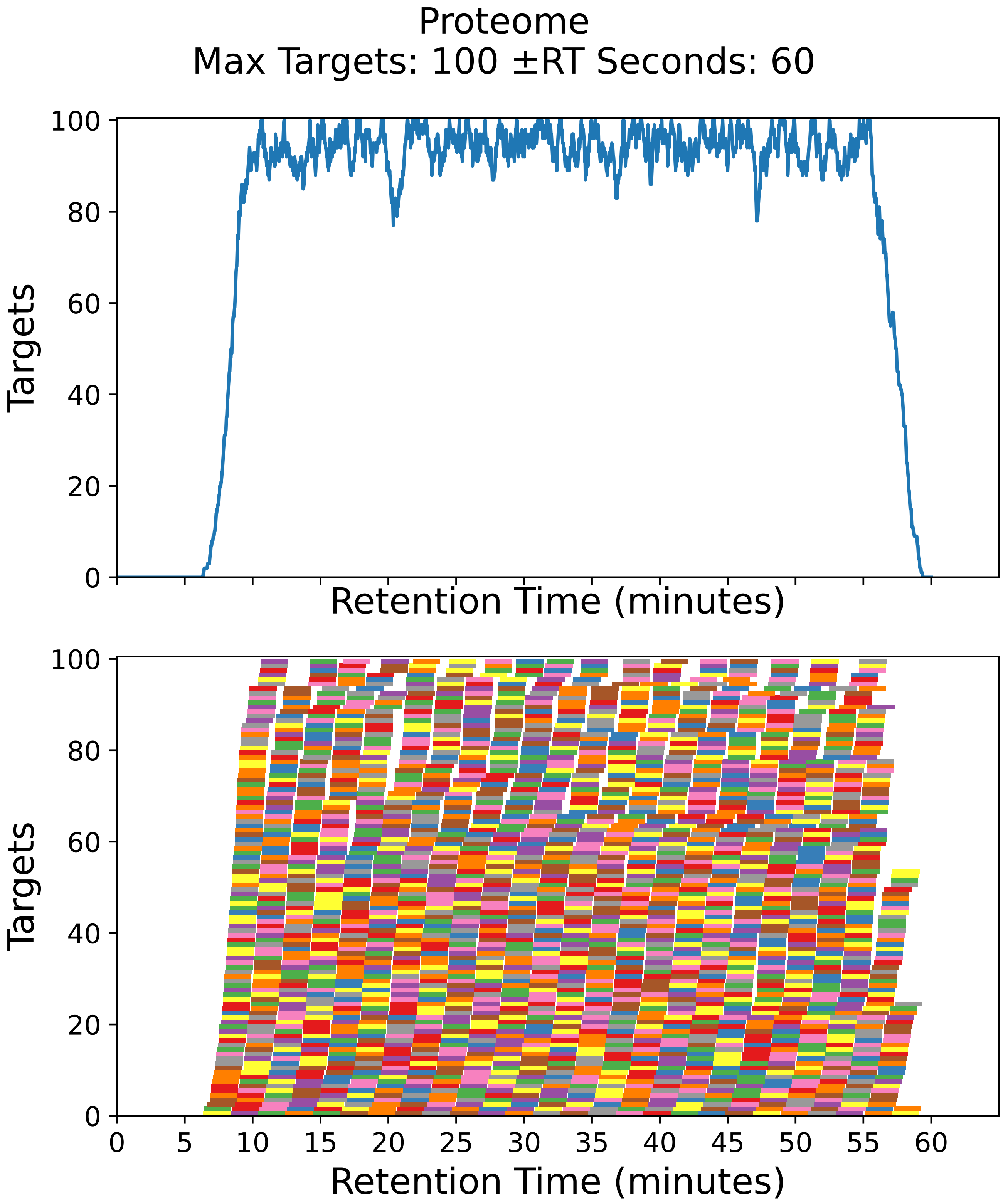

Supplementary Figure 17: Top: utilization diagram for the best fit method with instrument targeting the proteome filtered subset with a maximum of 100 targets per cycle with retention times ± 60 seconds from the peptide peak apex. Bottom: individual targeting windows for the 2294 peptides, monitoring 1147 proteins.

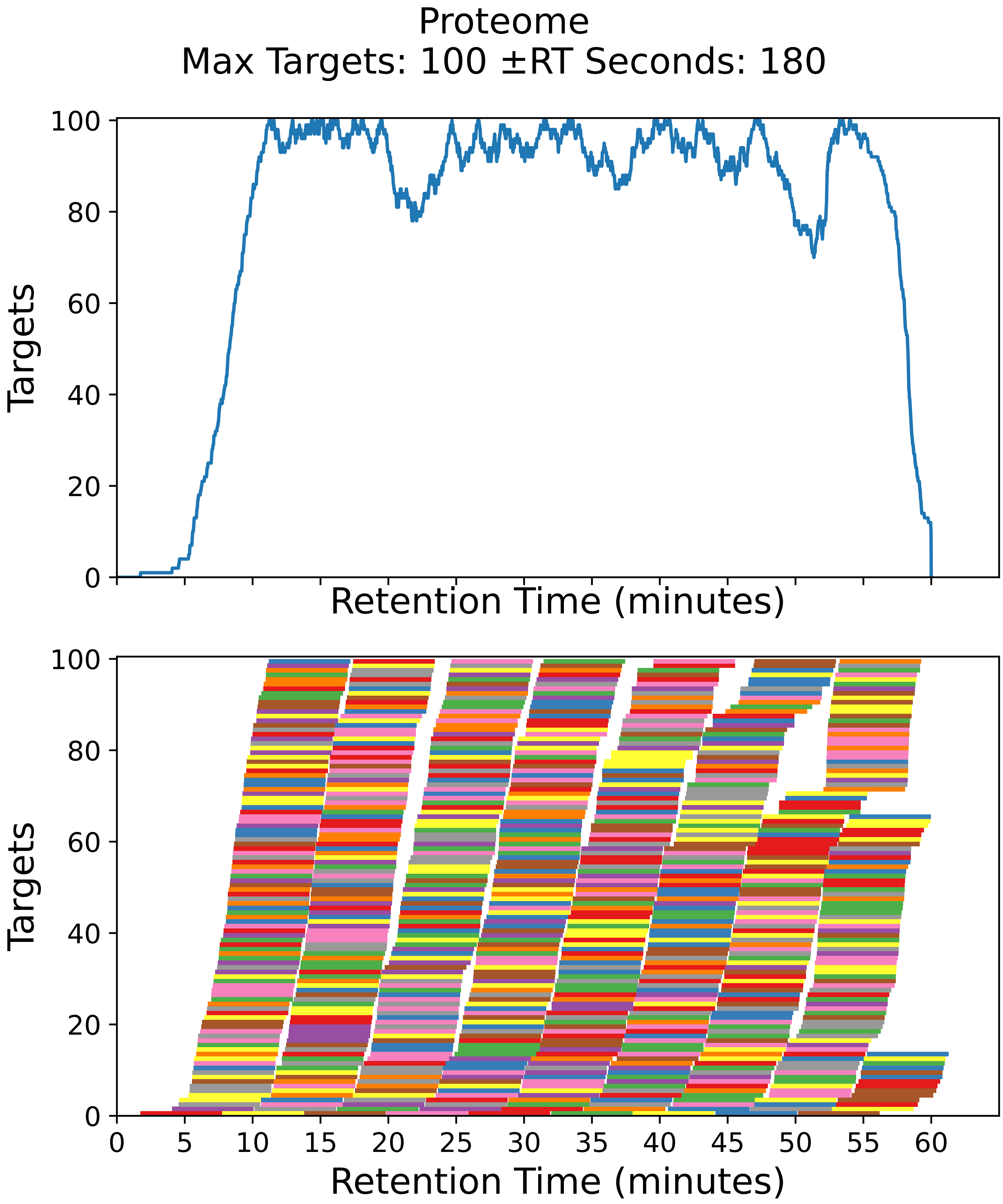

Supplementary Figure 18: Top: utilization diagram for the best fit method with instrument targeting the proteome filtered subset with a maximum of 100 targets per cycle with retention times ± 180 seconds from the peptide peak apex. Bottom: individual targeting windows for the 780 peptides, monitoring 390 proteins.

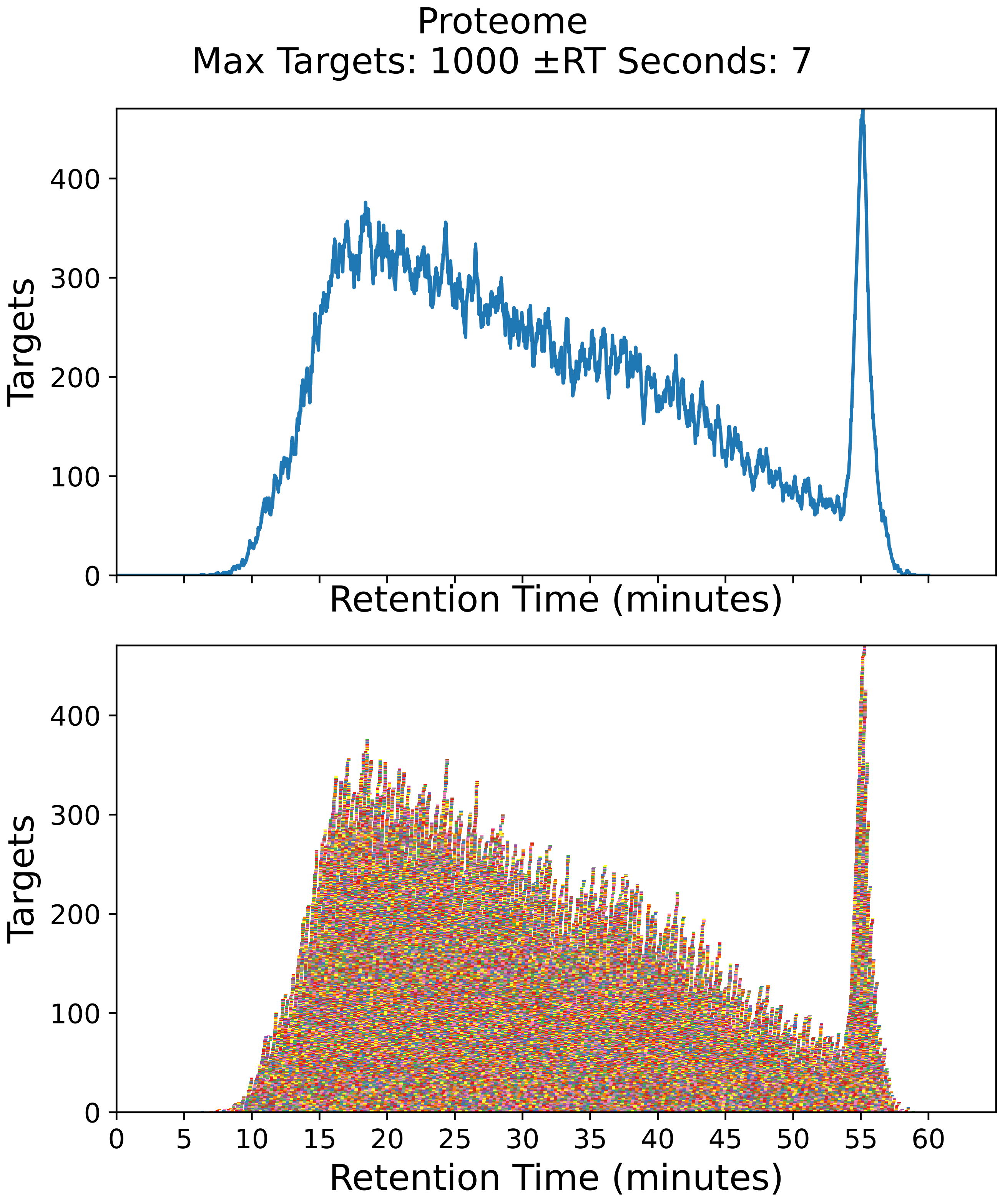

Supplementary Figure 19: Top: utilization diagram for the best fit method with instrument targeting the proteome filtered subset with a maximum of 1000 targets per cycle with retention times ± 7 seconds from the peptide peak apex. Bottom: individual targeting windows for the 38678 peptides, monitoring 19339 proteins.

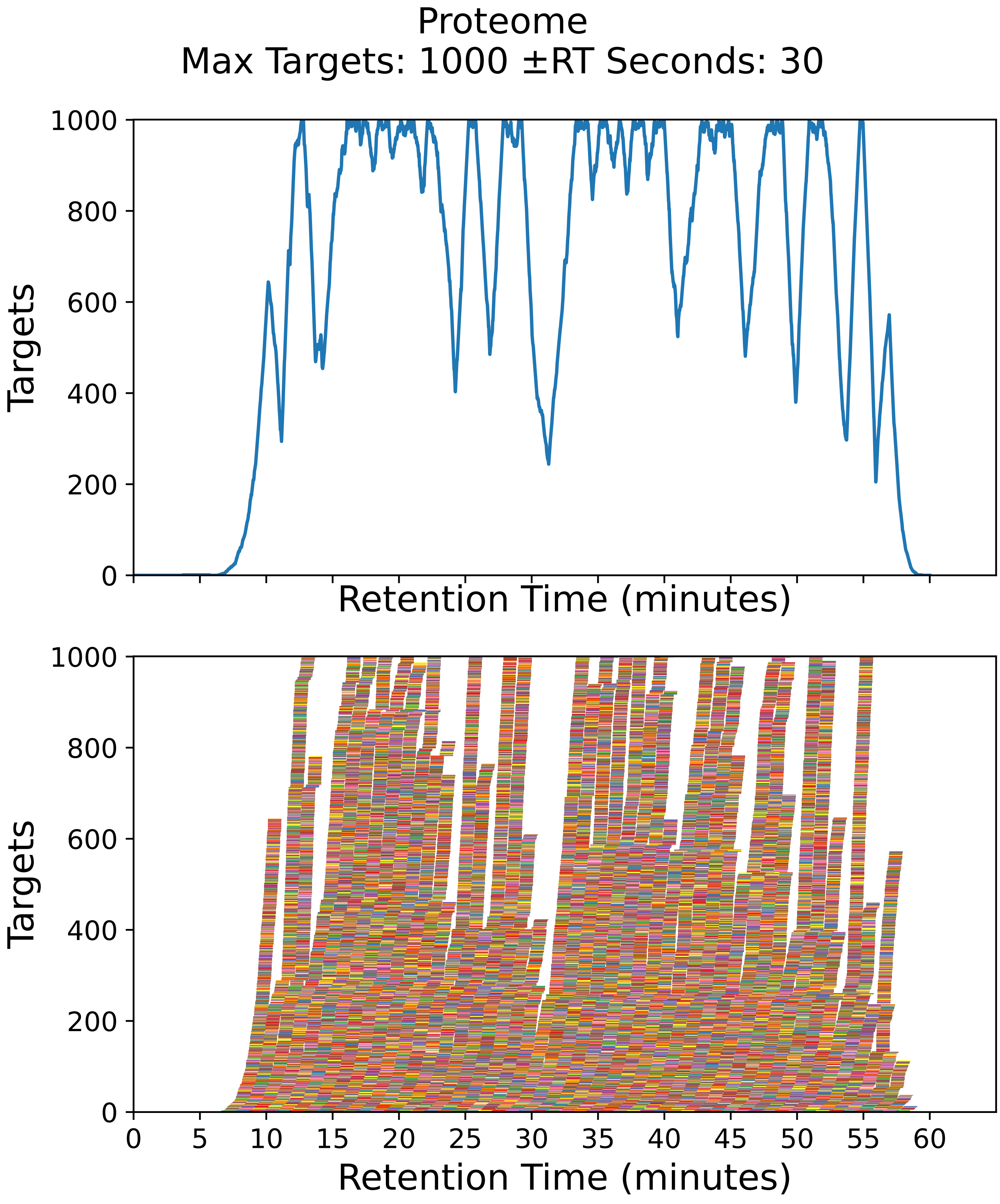

Supplementary Figure 20: Top: utilization diagram for the best fit method with instrument targeting the proteome filtered subset with a maximum of 1000 targets per cycle with retention times ± 30 seconds from the peptide peak apex. Bottom: individual targeting windows for the 37694 peptides, monitoring 18847 proteins.

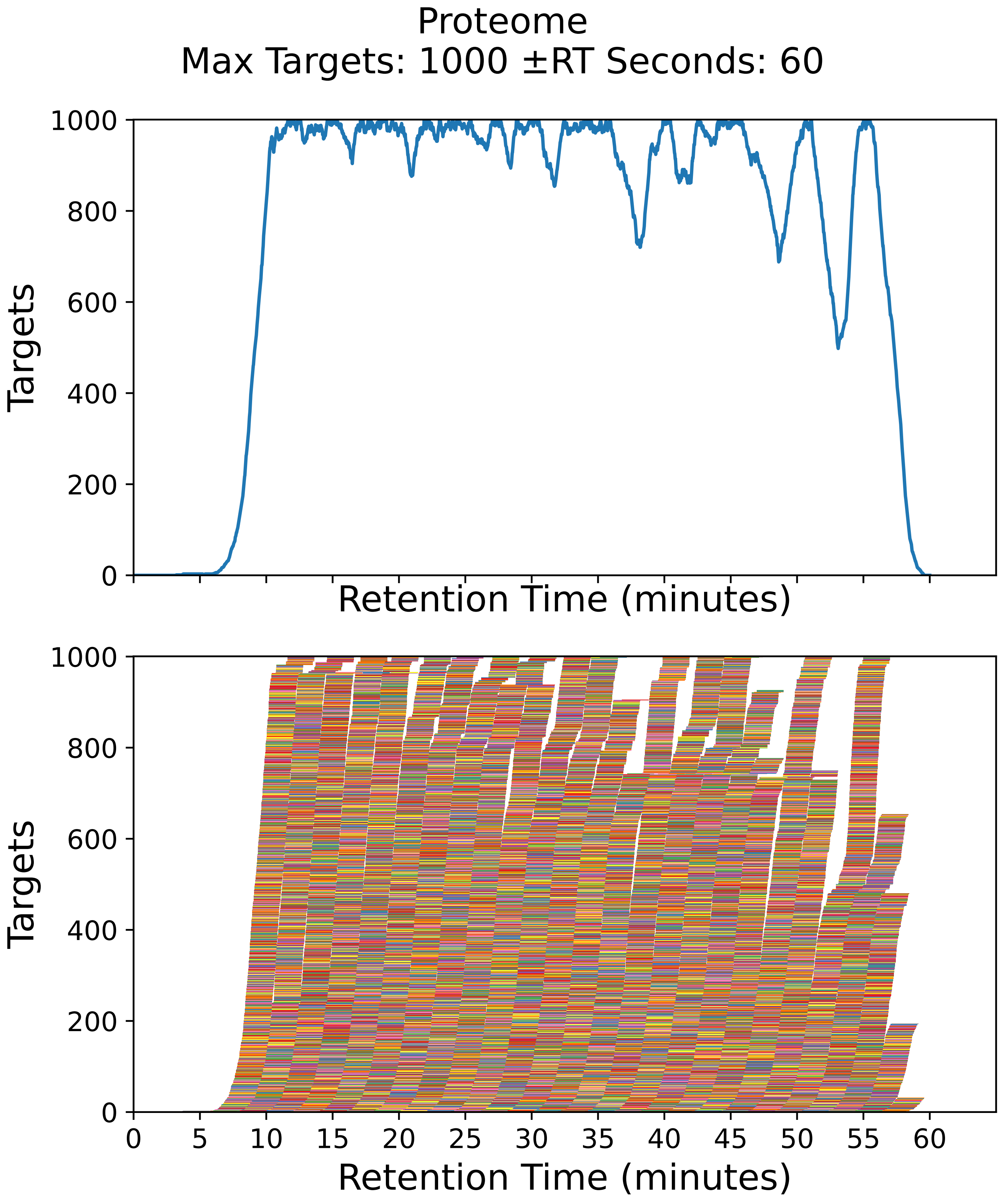

Supplementary Figure 21: Top: utilization diagram for the best fit method with instrument targeting the proteome filtered subset with a maximum of 1000 targets per cycle with retention times ± 60 seconds from the peptide peak apex. Bottom: individual targeting windows for the 22354 peptides, monitoring 11177 proteins.

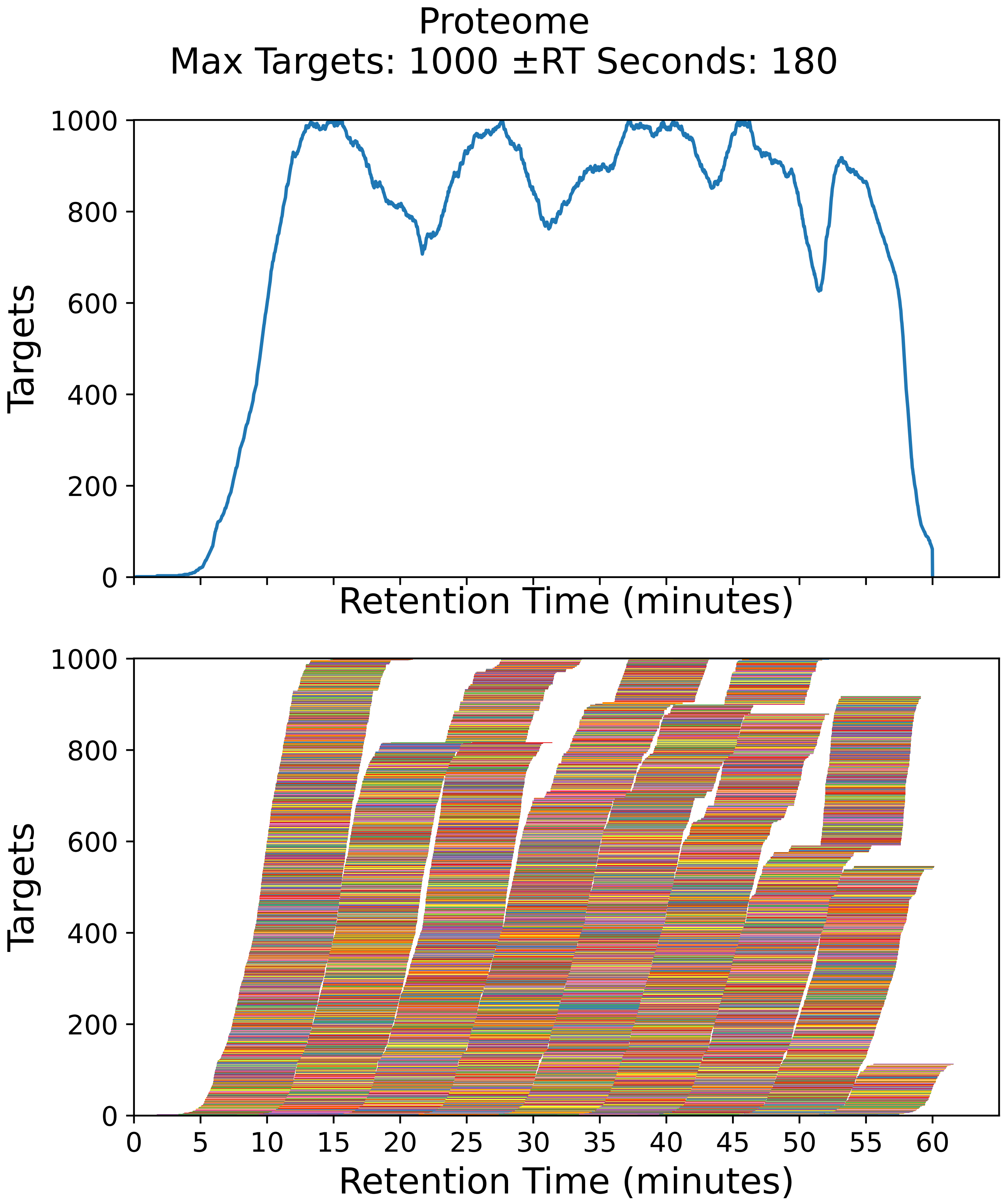

Supplementary Figure 22: Top: utilization diagram for the best fit method with instrument targeting the proteome filtered subset with a maximum of 1000 targets per cycle with retention times ± 180 seconds from the peptide peak apex. Bottom: individual targeting windows for the 7274 peptides, monitoring 3637 proteins.

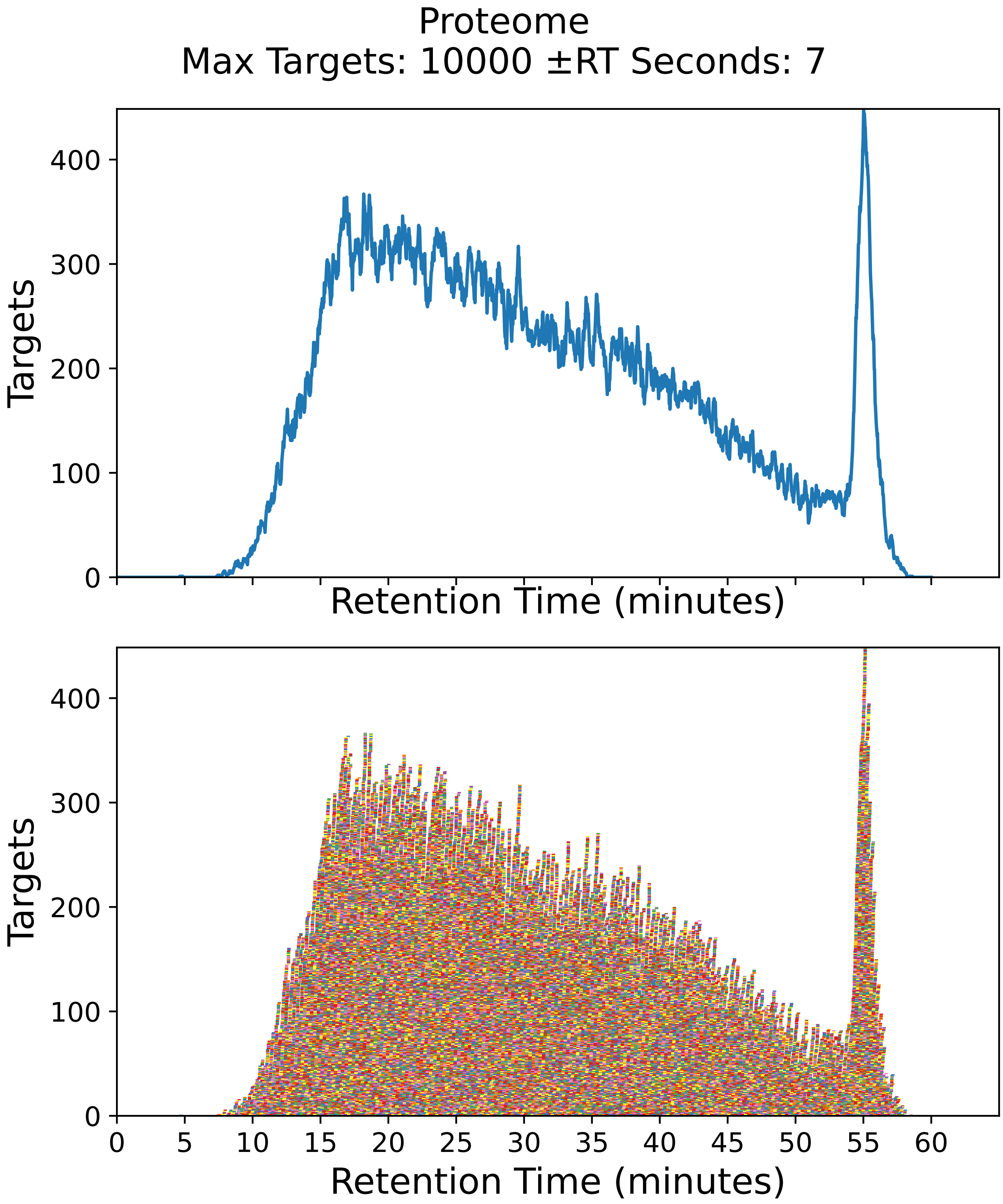

Supplementary Figure 23: Top: utilization diagram for the best fit method with instrument targeting the proteome filtered subset with a maximum of 10000 targets per cycle with retention times ± 7 seconds from the peptide peak apex. Bottom: individual targeting windows for the 38678 peptides, monitoring 19339 proteins.

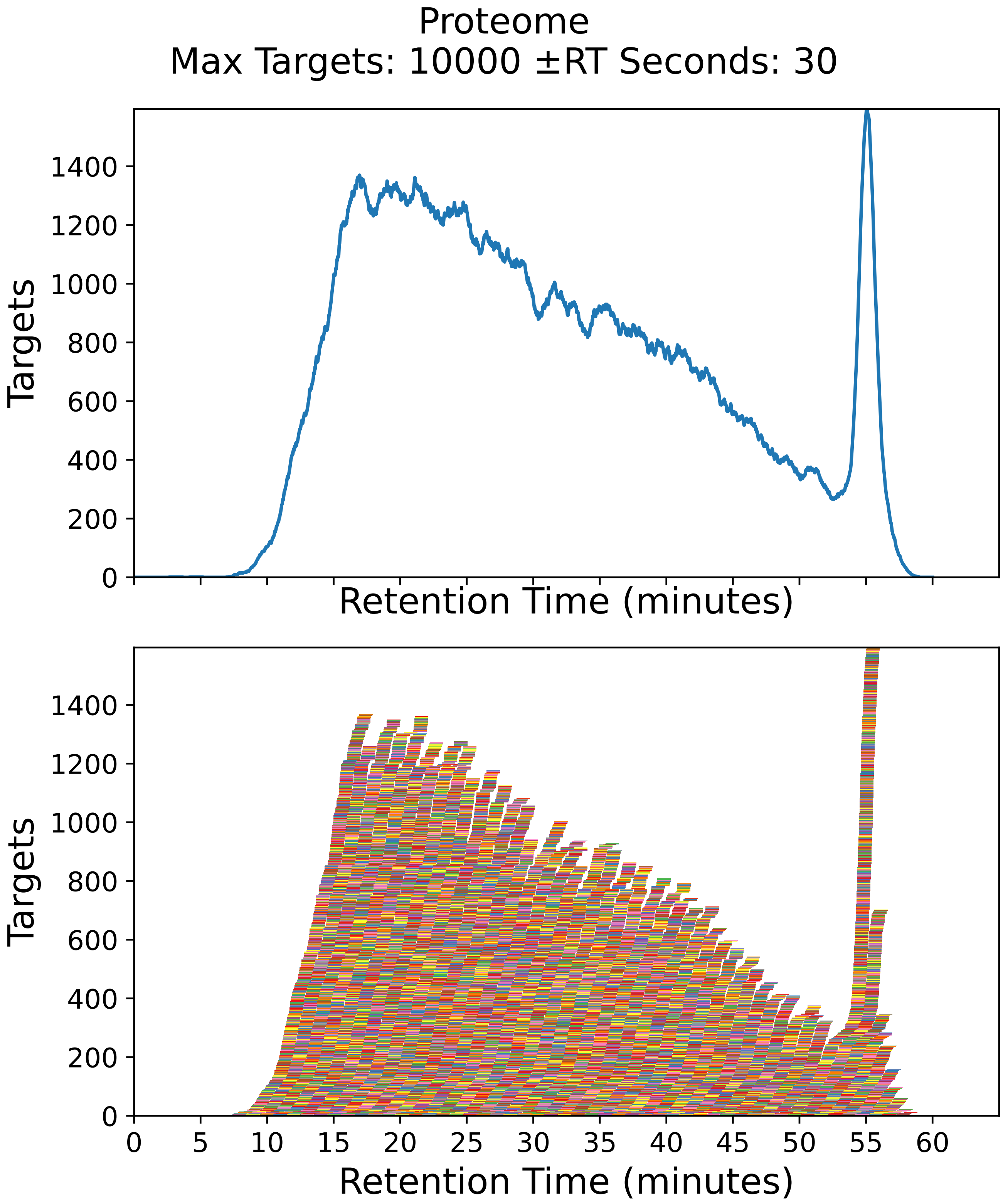

Supplementary Figure 24: Top: utilization diagram for the best fit method with instrument targeting the proteome filtered subset with a maximum of 10000 targets per cycle with retention times ± 30 seconds from the peptide peak apex. Bottom: individual targeting windows for the 38678 peptides, monitoring 19339 proteins.

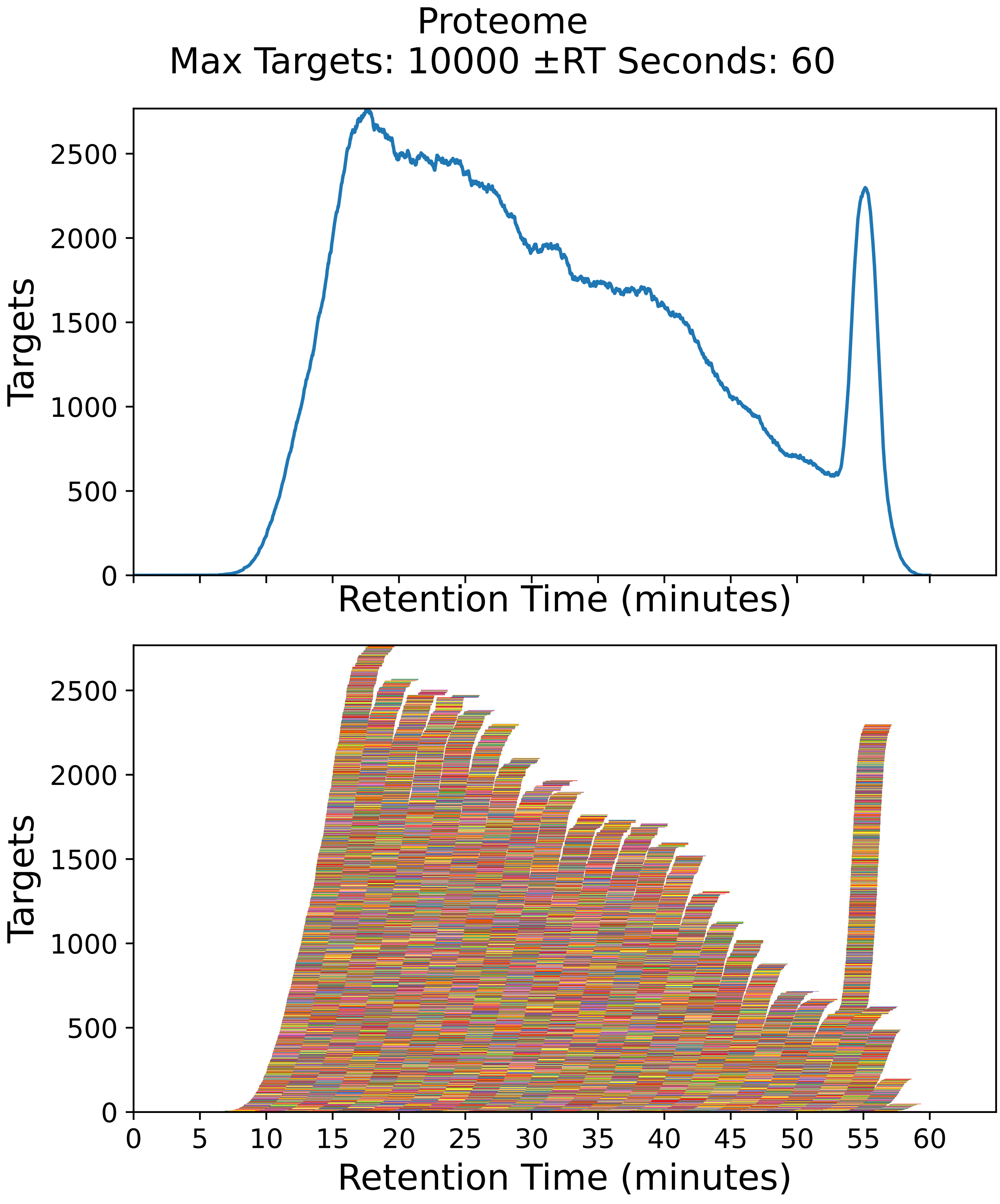

Supplementary Figure 25: Top: utilization diagram for the best fit method with instrument targeting the proteome filtered subset with a maximum of 10000 targets per cycle with retention times ± 60 seconds from the peptide peak apex. Bottom: individual targeting windows for the 38678 peptides, monitoring 19339 proteins.

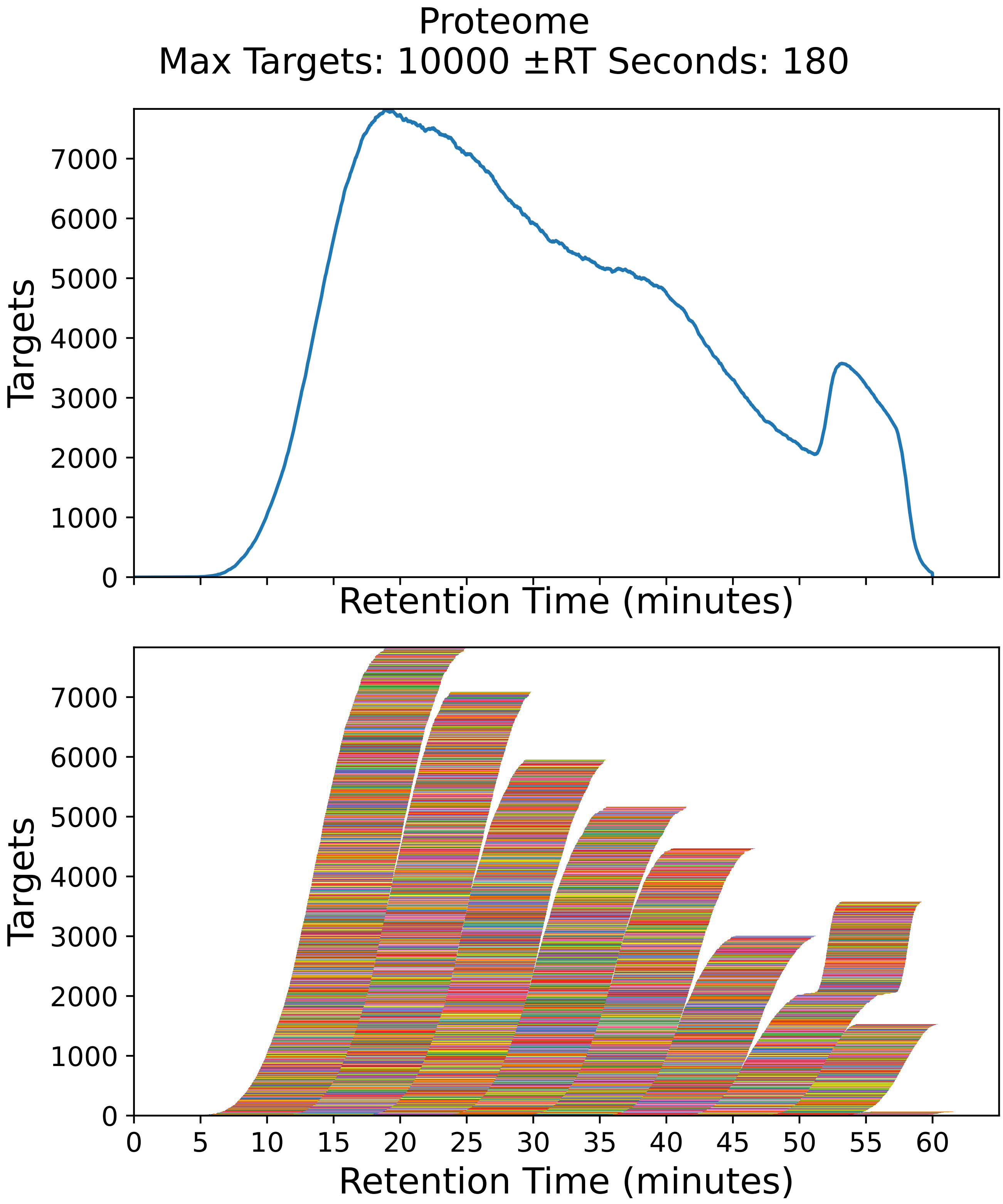

Supplementary Figure 26: Top: utilization diagram for the best fit method with instrument targeting the proteome filtered subset with a maximum of 10000 targets per cycle with retention times ± 180 seconds from the peptide peak apex. Bottom: individual targeting windows for the 38678 peptides, monitoring 19339 proteins.

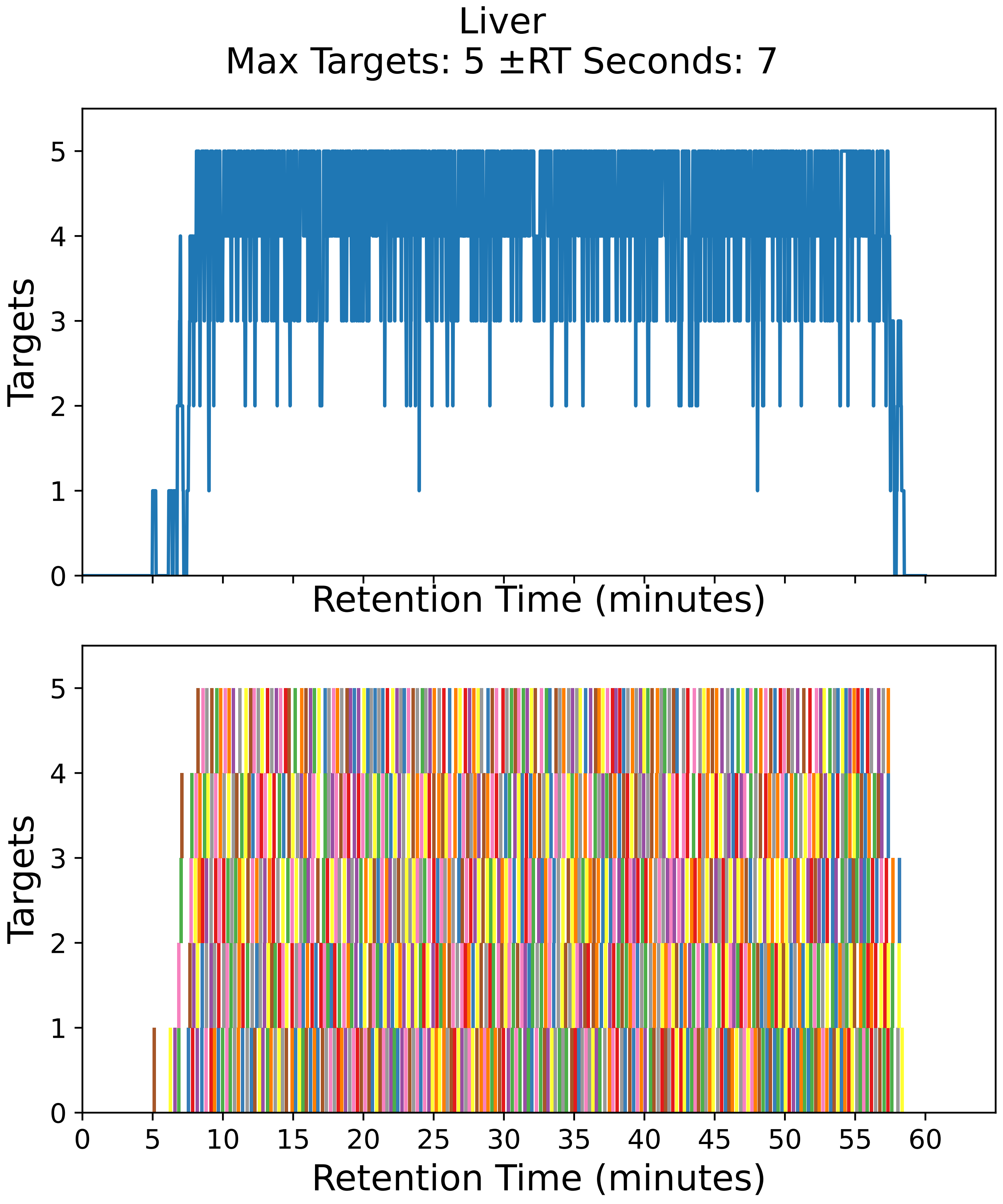

Supplementary Figure 27: Top: utilization diagram for the best fit method with instrument targeting the liver filtered subset with a maximum of 5 targets per cycle with retention times ± 7 seconds from the peptide peak apex. Bottom: individual targeting windows for the 858 peptides, monitoring 429 proteins.

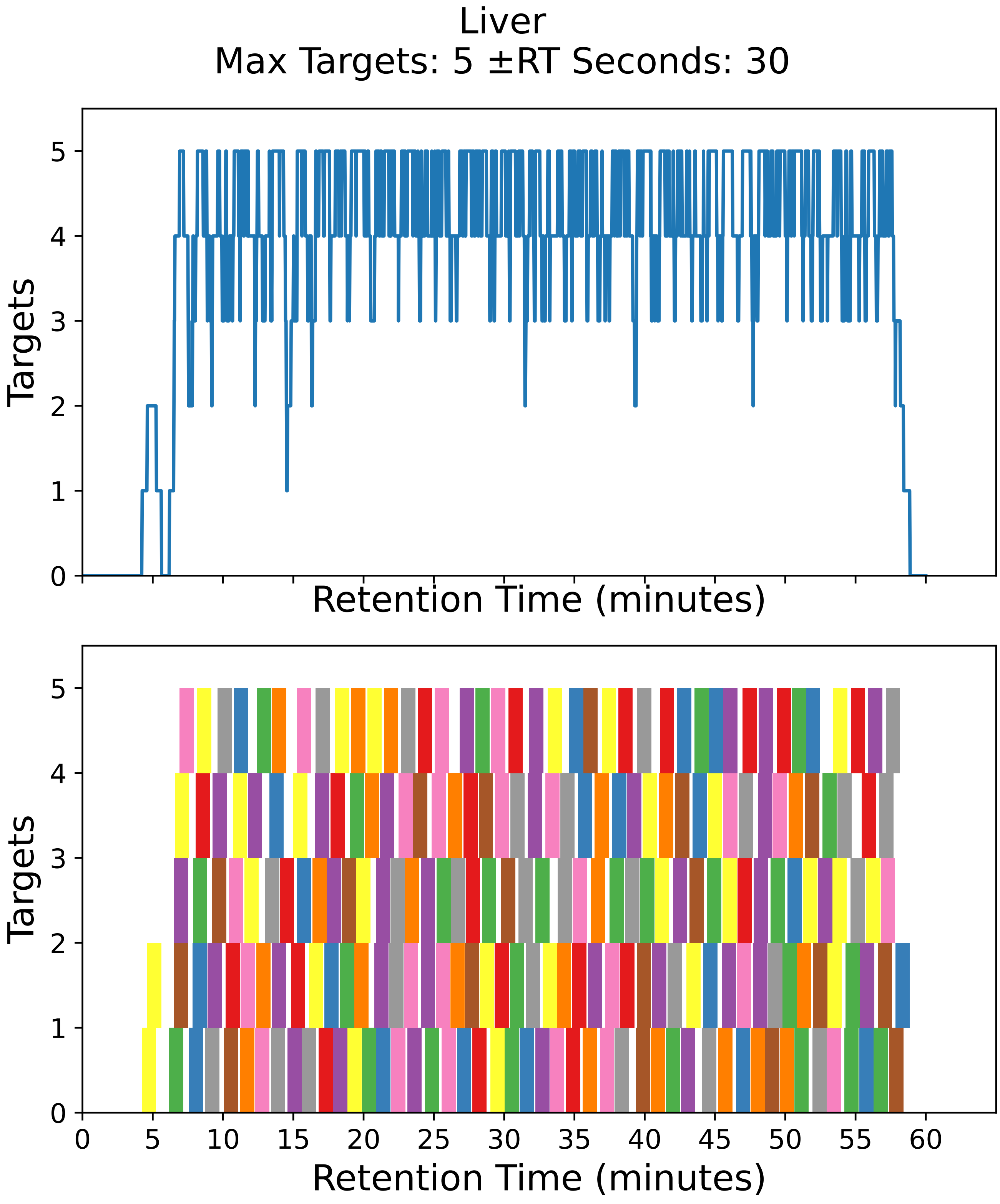

Supplementary Figure 28: Top: utilization diagram for the best fit method with instrument targeting the liver filtered subset with a maximum of 5 targets per cycle with retention times ± 30 seconds from the peptide peak apex. Bottom: individual targeting windows for the 220 peptides, monitoring 110 proteins.

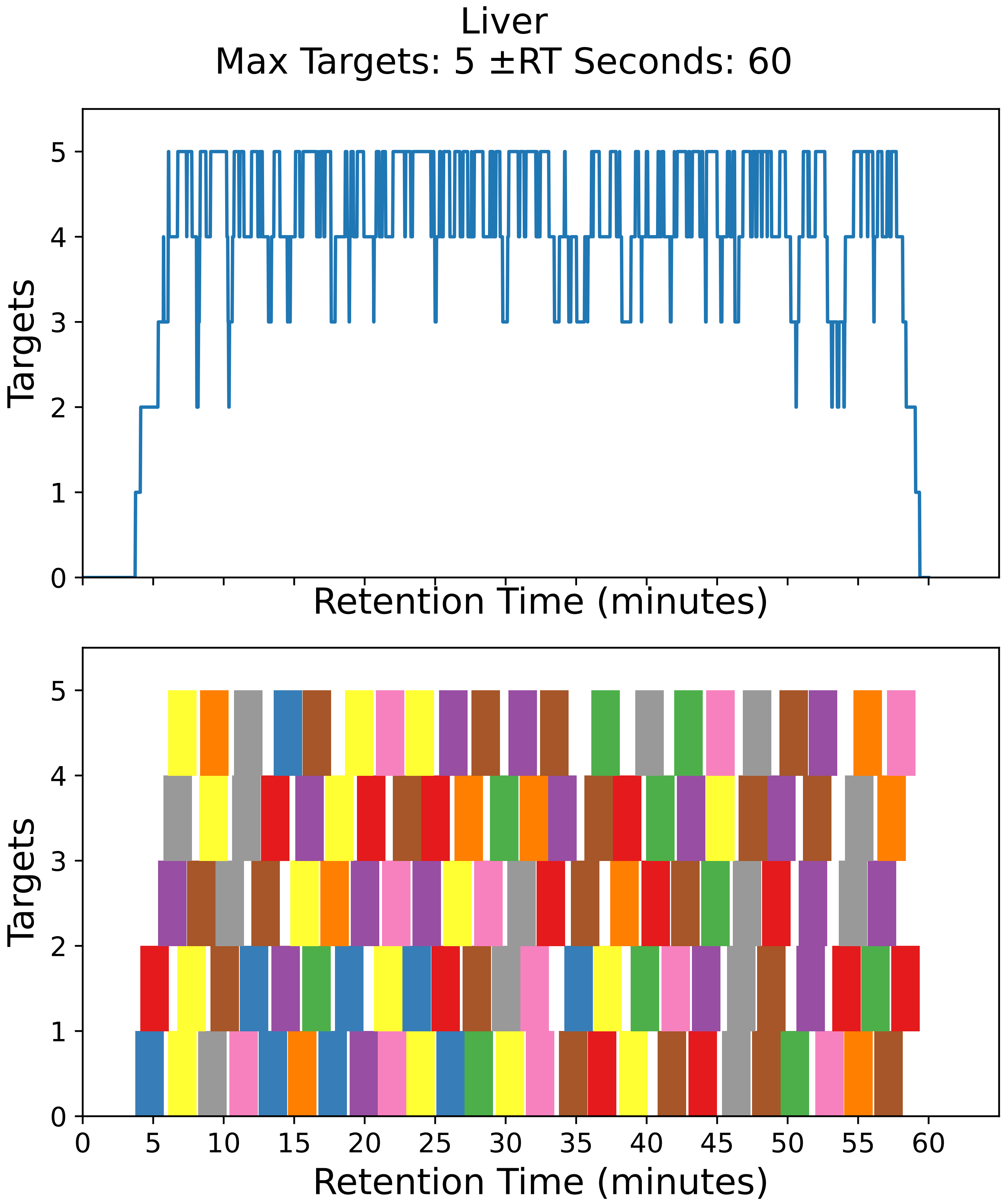

Supplementary Figure 29: Top: utilization diagram for the best fit method with instrument targeting the liver filtered subset with a maximum of 5 targets per cycle with retention times ± 60 seconds from the peptide peak apex. Bottom: individual targeting windows for the 116 peptides, monitoring 58 proteins.

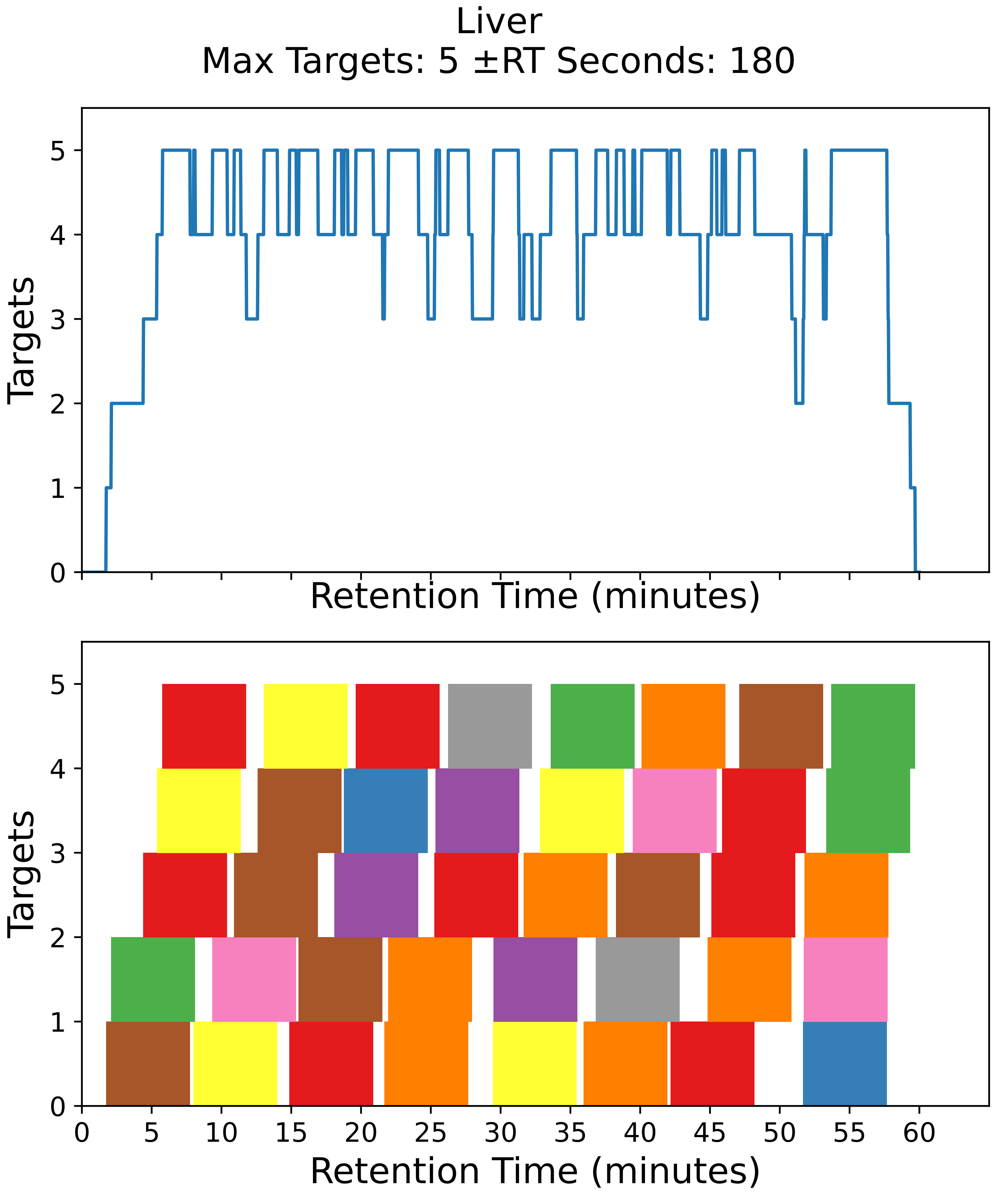

Supplementary Figure 30: Top: utilization diagram for the best fit method with instrument targeting the liver filtered subset with a maximum of 5 targets per cycle with retention times ± 180 seconds from the peptide peak apex. Bottom: individual targeting windows for the 40 peptides, monitoring 20 proteins.

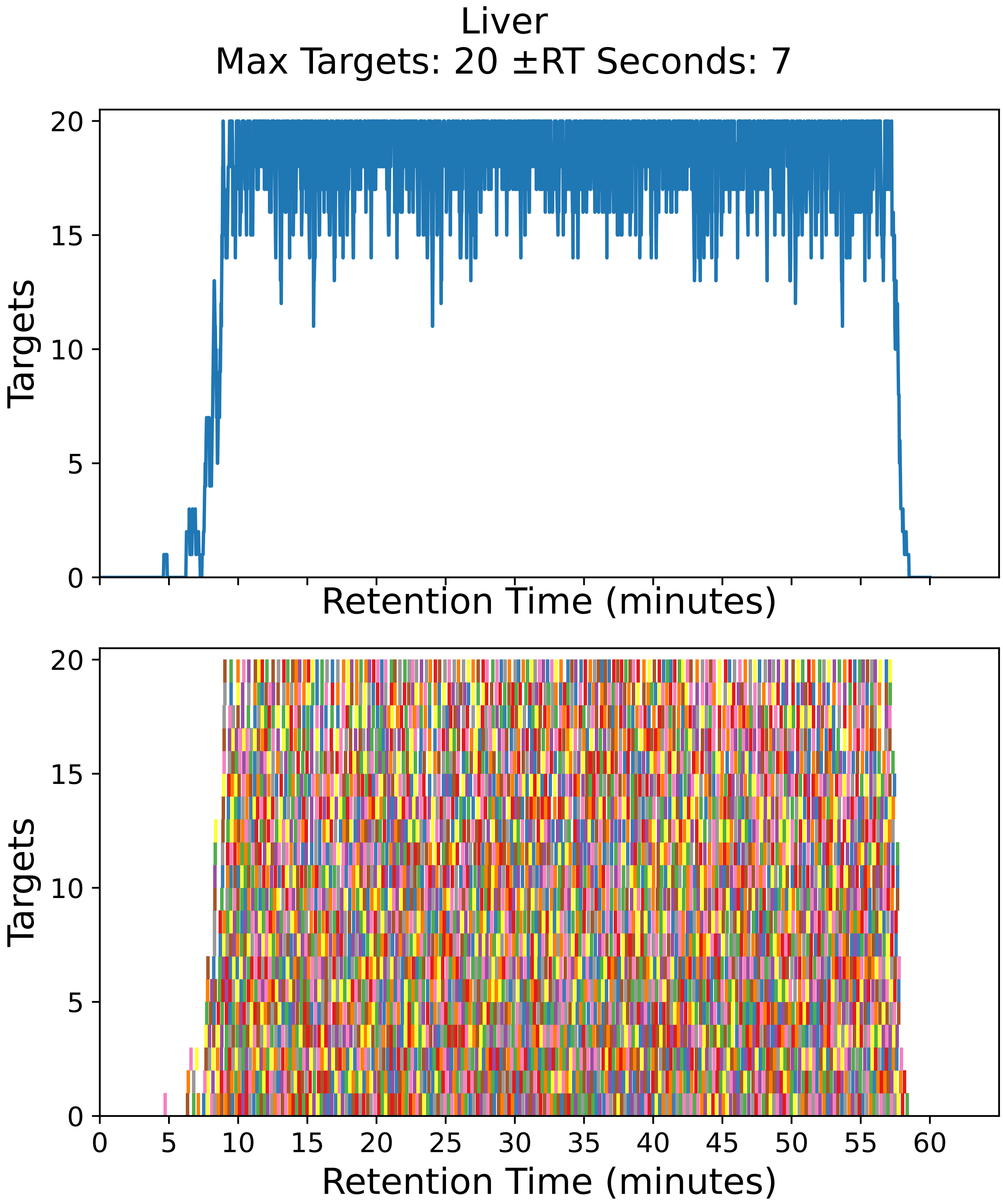

Supplementary Figure 31: Top: utilization diagram for the best fit method with instrument targeting the liver filtered subset with a maximum of 20 targets per cycle with retention times ± 7 seconds from the peptide peak apex. Bottom: individual targeting windows for the 3594 peptides, monitoring 1797 proteins.

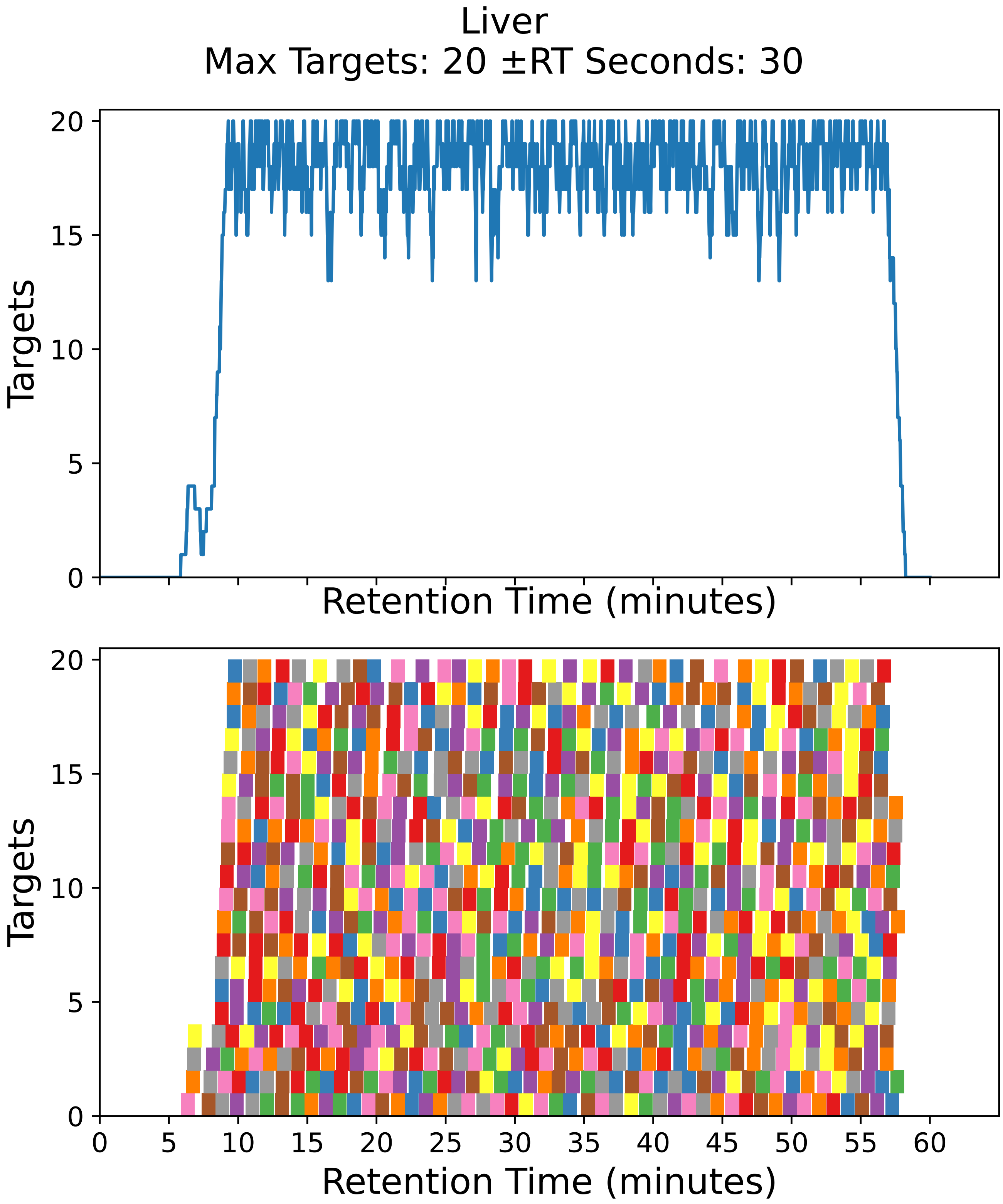

Supplementary Figure 32: Top: utilization diagram for the best fit method with instrument targeting the liver filtered subset with a maximum of 20 targets per cycle with retention times ± 30 seconds from the peptide peak apex. Bottom: individual targeting windows for the 880 peptides, monitoring 440 proteins.

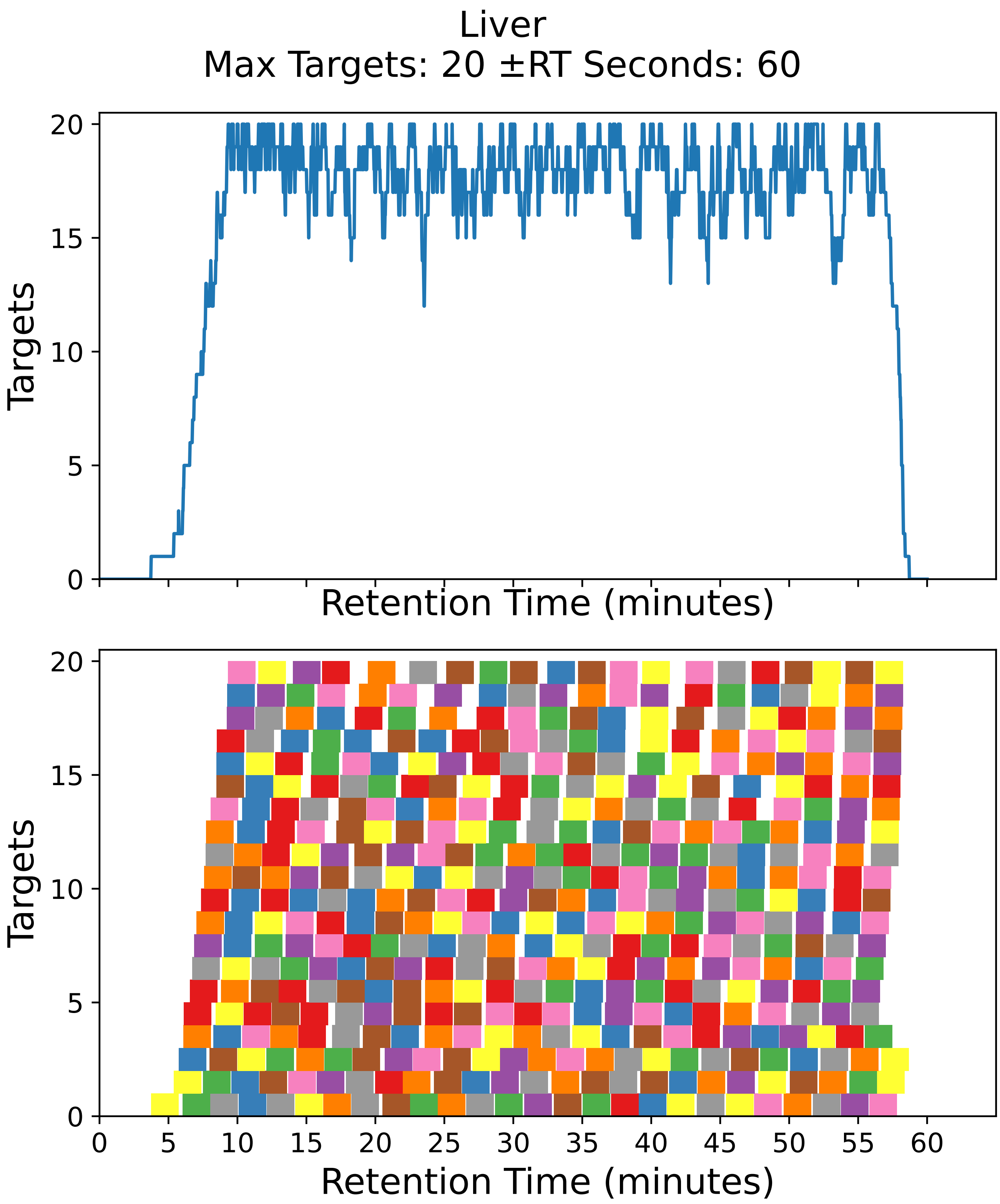

Supplementary Figure 33: Top: utilization diagram for the best fit method with instrument targeting the liver filtered subset with a maximum of 20 targets per cycle with retention times ± 60 seconds from the peptide peak apex. Bottom: individual targeting windows for the 450 peptides, monitoring 225 proteins.

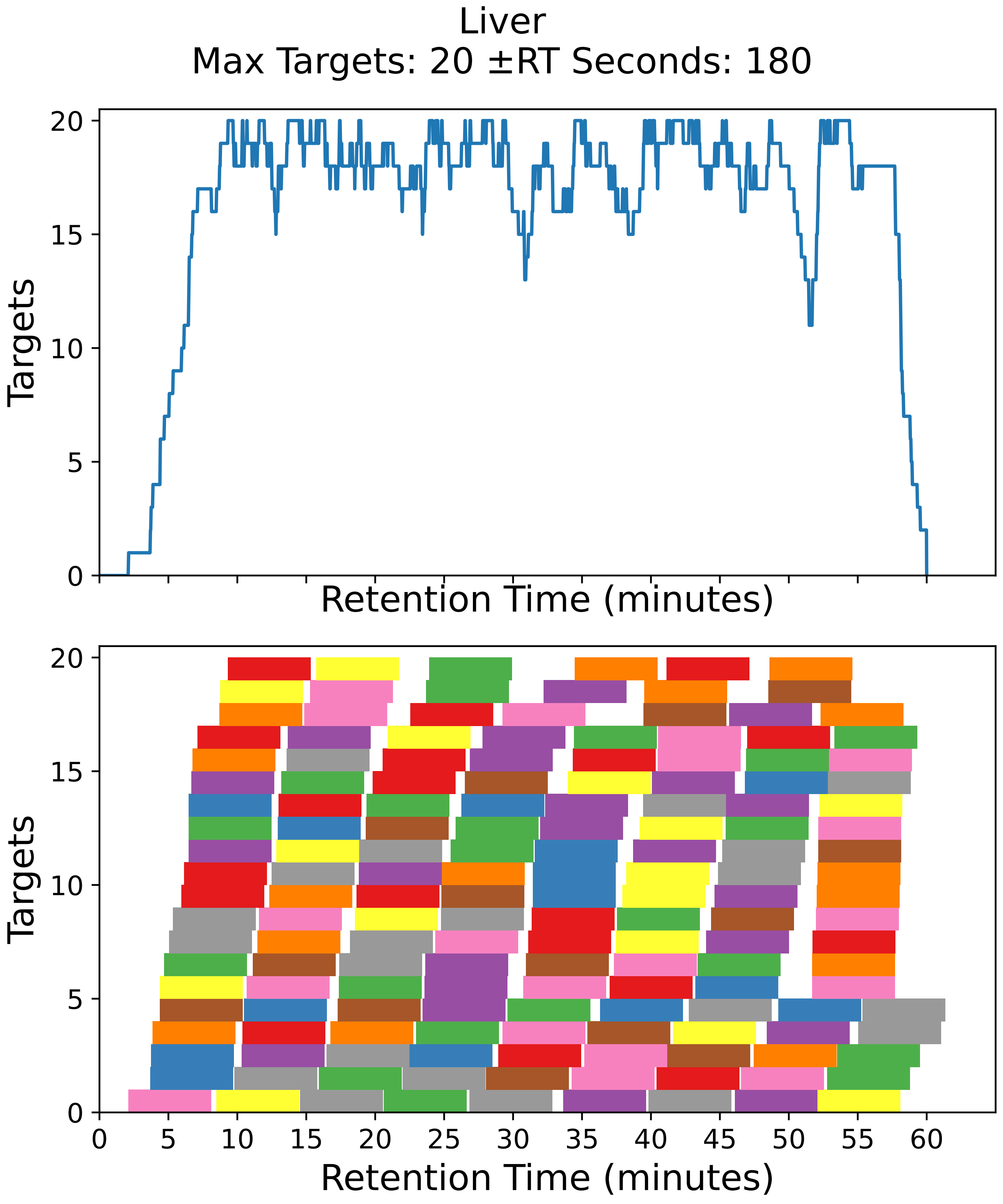

Supplementary Figure 34: Top: utilization diagram for the best fit method with instrument targeting the liver filtered subset with a maximum of 20 targets per cycle with retention times ± 180 seconds from the peptide peak apex. Bottom: individual targeting windows for the 160 peptides, monitoring 80 proteins.

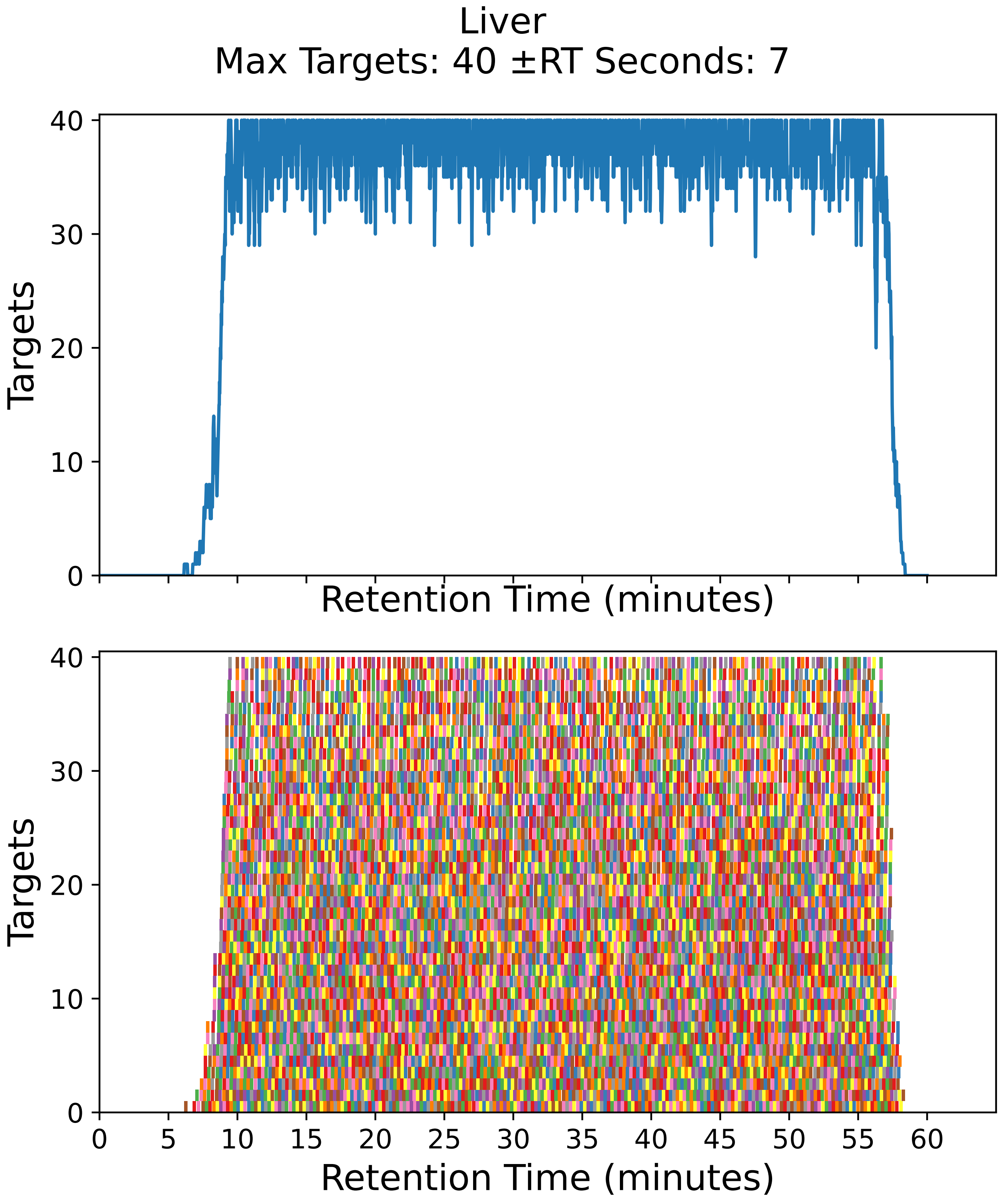

Supplementary Figure 35: Top: utilization diagram for the best fit method with instrument targeting the liver filtered subset with a maximum of 40 targets per cycle with retention times ± 7 seconds from the peptide peak apex. Bottom: individual targeting windows for the 7290 peptides, monitoring 3645 proteins.

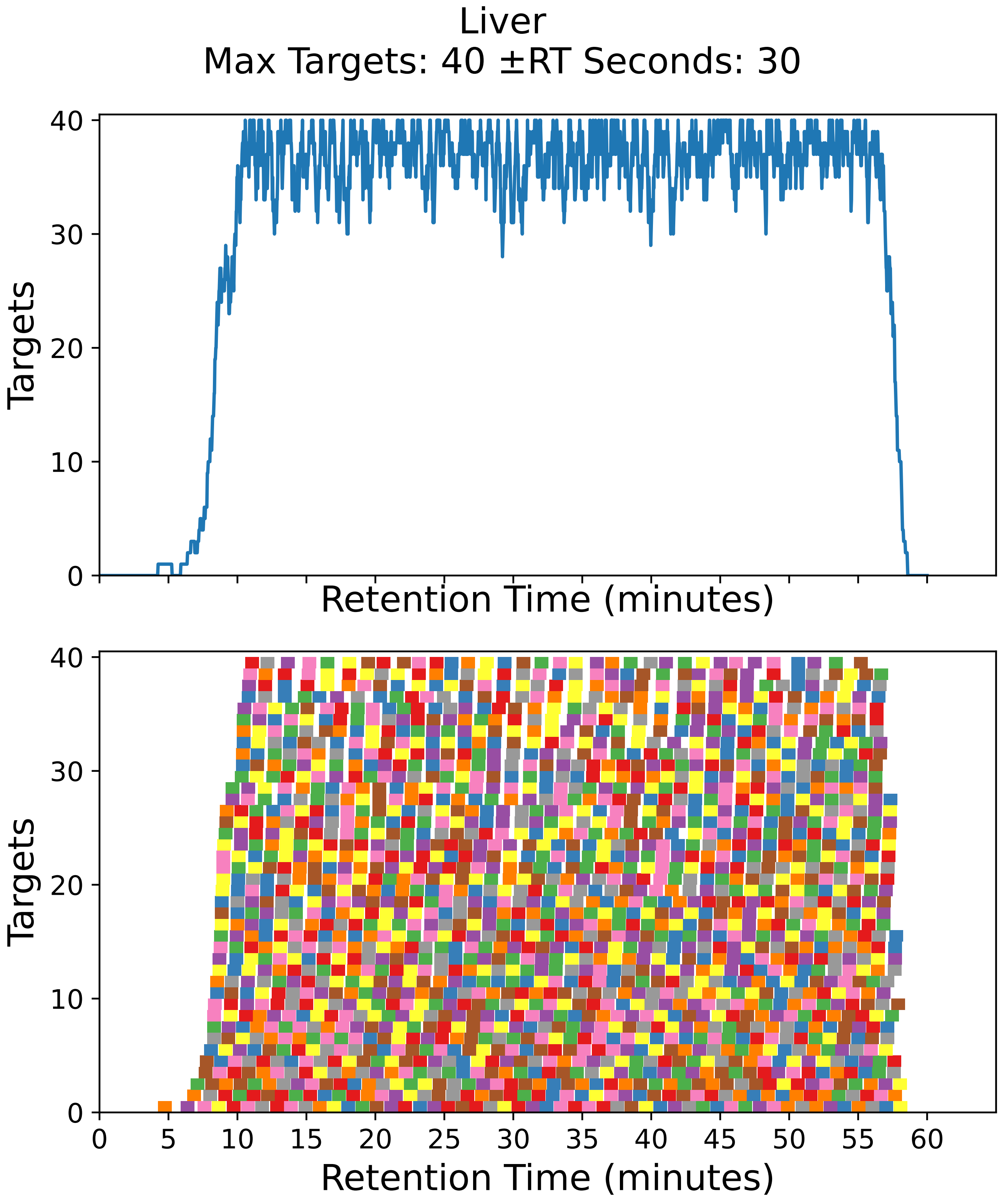

Supplementary Figure 36: Top: utilization diagram for the best fit method with instrument targeting the liver filtered subset with a maximum of 40 targets per cycle with retention times ± 30 seconds from the peptide peak apex. Bottom: individual targeting windows for the 1778 peptides, monitoring 889 proteins.

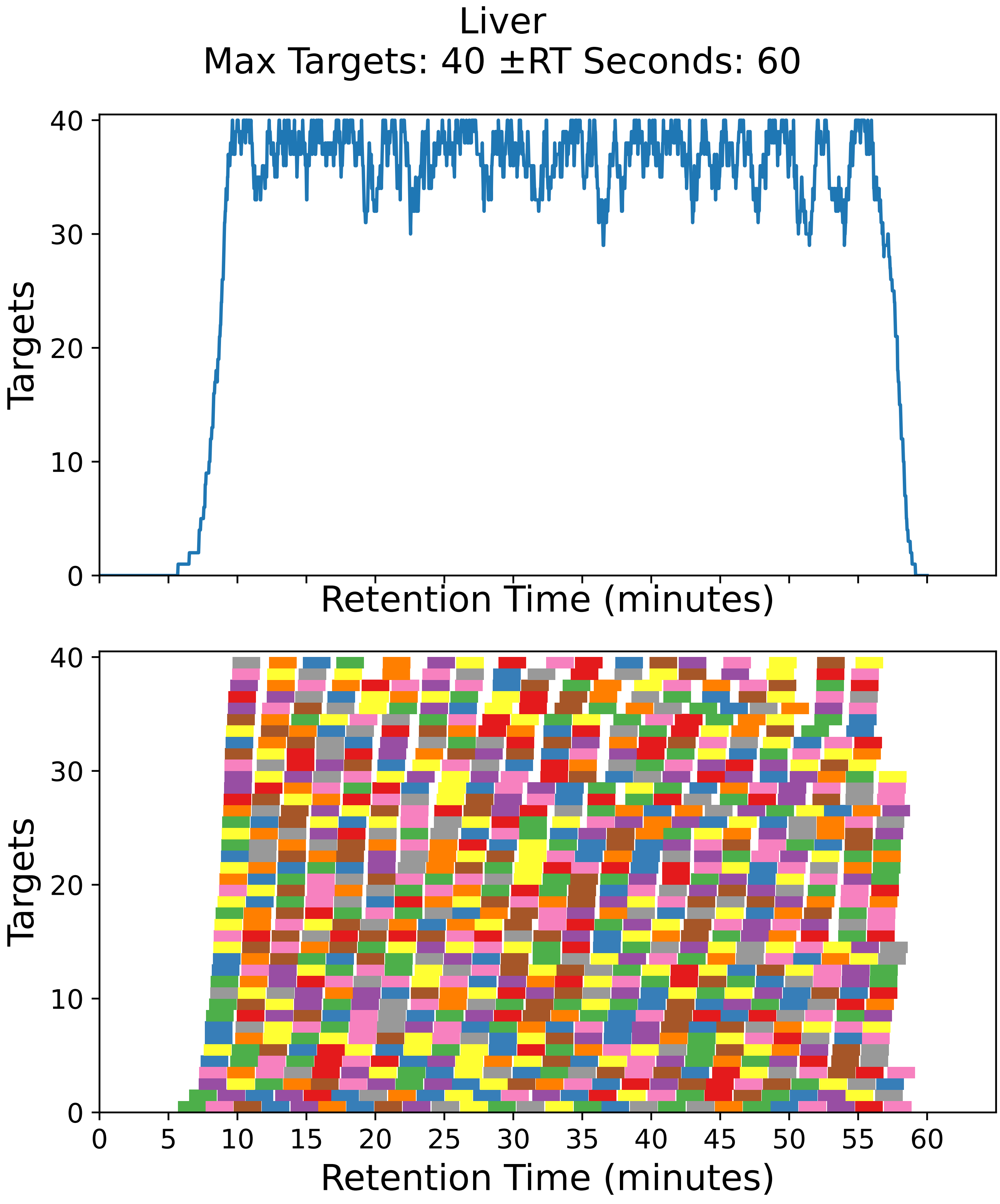

Supplementary Figure 37: Top: utilization diagram for the best fit method with instrument targeting the liver filtered subset with a maximum of 40 targets per cycle with retention times ± 60 seconds from the peptide peak apex. Bottom: individual targeting windows for the 902 peptides, monitoring 451 proteins.

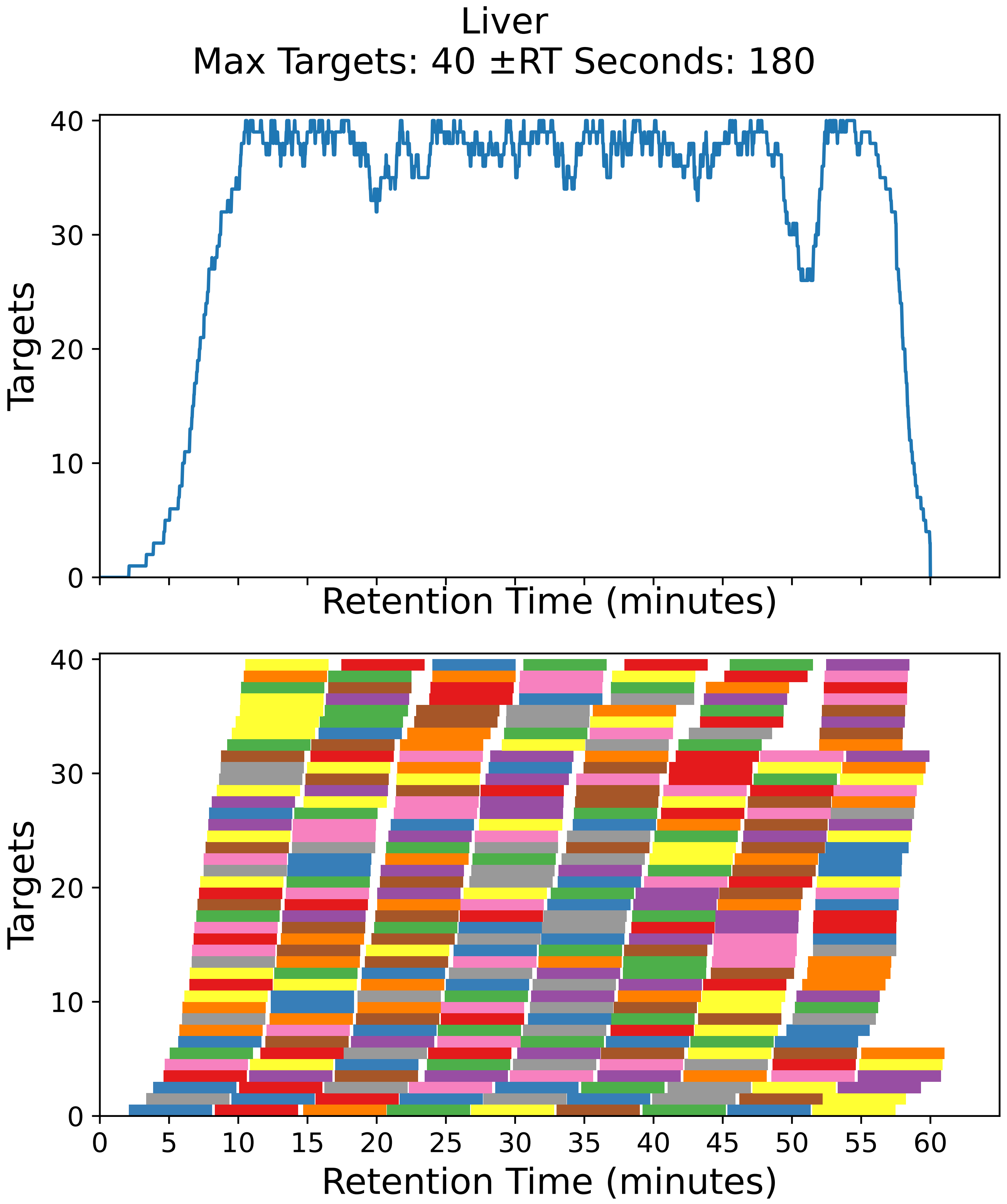

Supplementary Figure 38: Top: utilization diagram for the best fit method with instrument targeting the liver filtered subset with a maximum of 40 targets per cycle with retention times ± 180 seconds from the peptide peak apex. Bottom: individual targeting windows for the 318 peptides, monitoring 159 proteins.

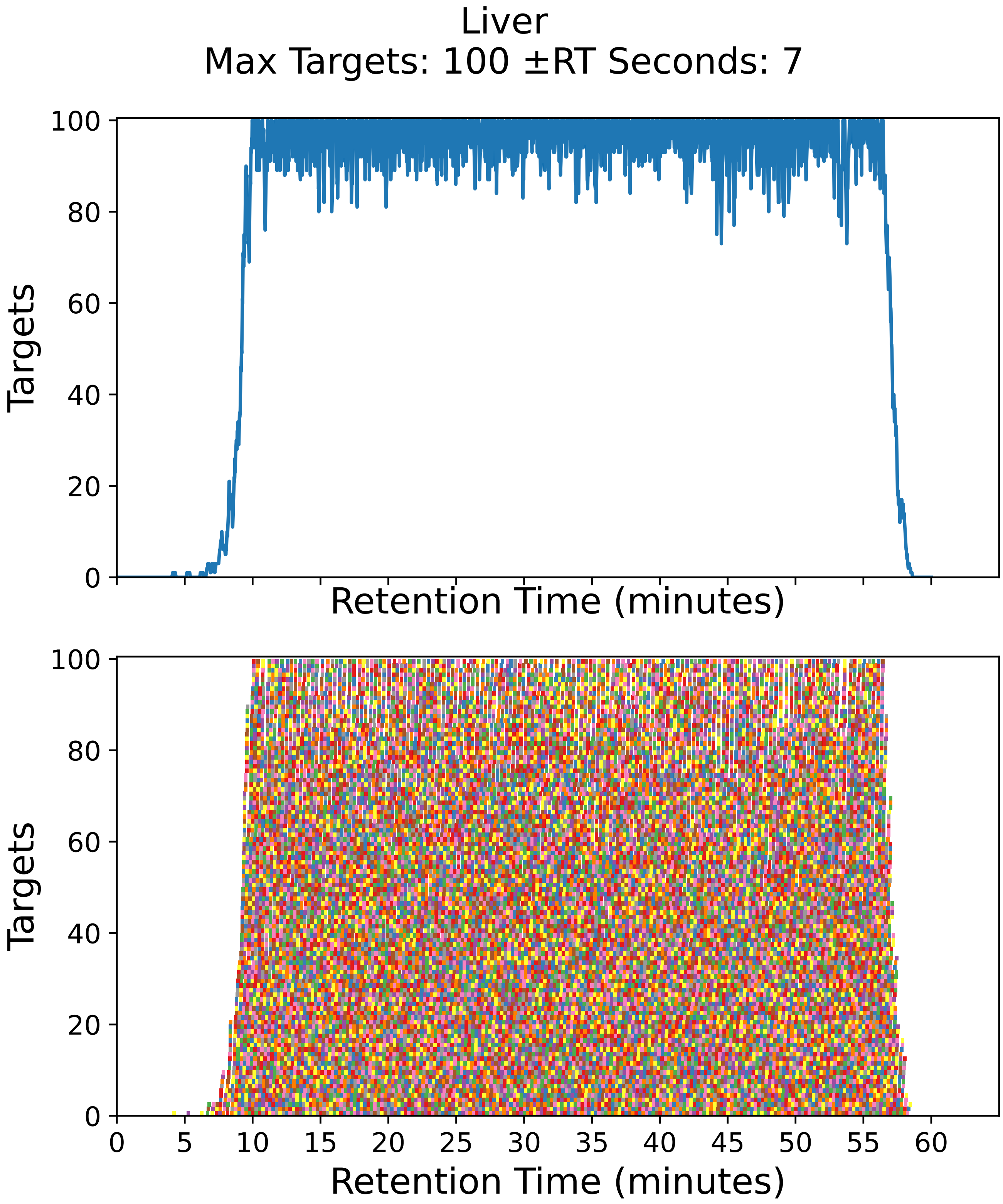

Supplementary Figure 39: Top: utilization diagram for the best fit method with instrument targeting the liver filtered subset with a maximum of 100 targets per cycle with retention times ± 7 seconds from the peptide peak apex. Bottom: individual targeting windows for the 18362 peptides, monitoring 9181 proteins.

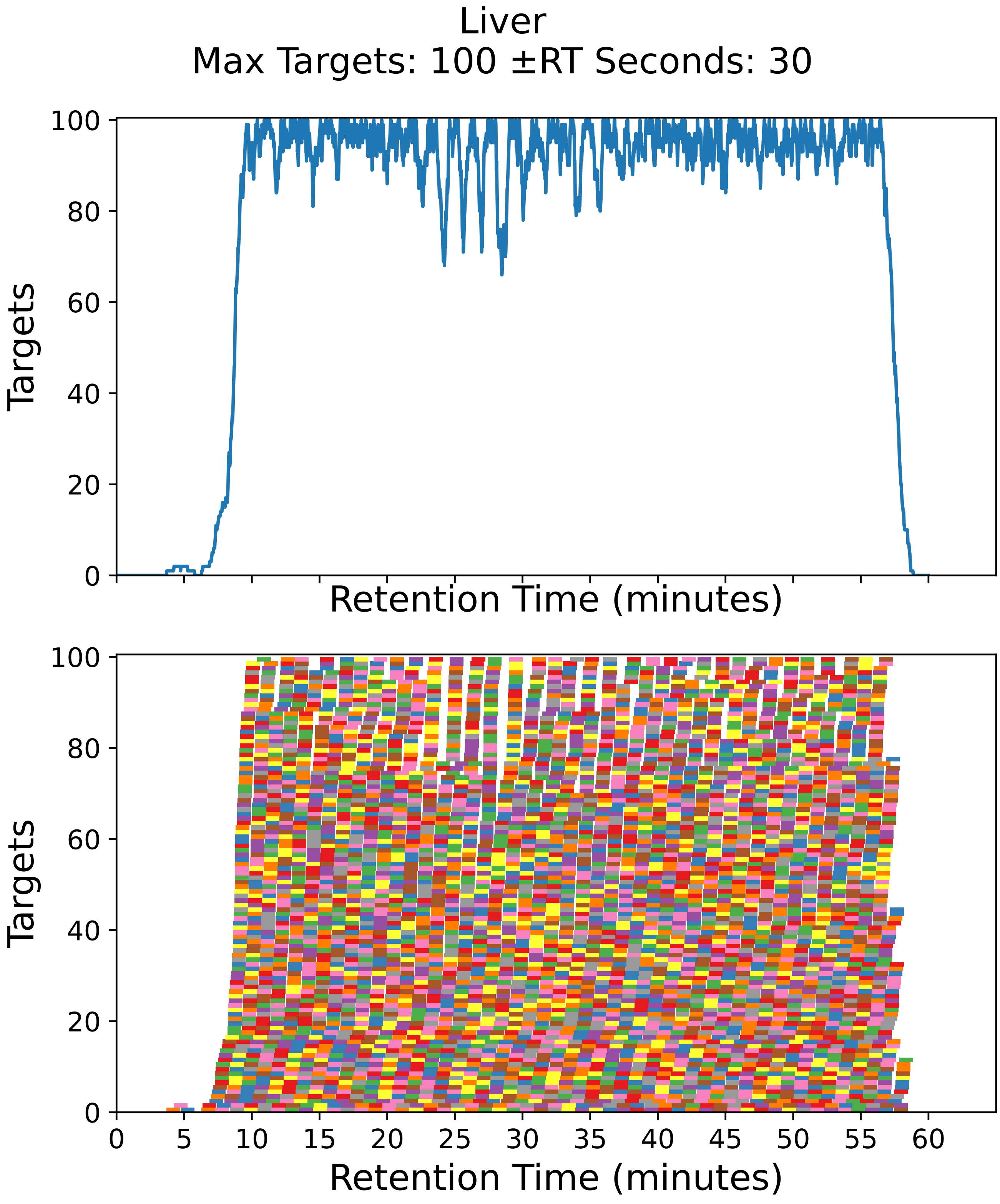

Supplementary Figure 40: Top: utilization diagram for the best fit method with instrument targeting the liver filtered subset with a maximum of 100 targets per cycle with retention times ± 30 seconds from the peptide peak apex. Bottom: individual targeting windows for the 4538 peptides, monitoring 2269 proteins.

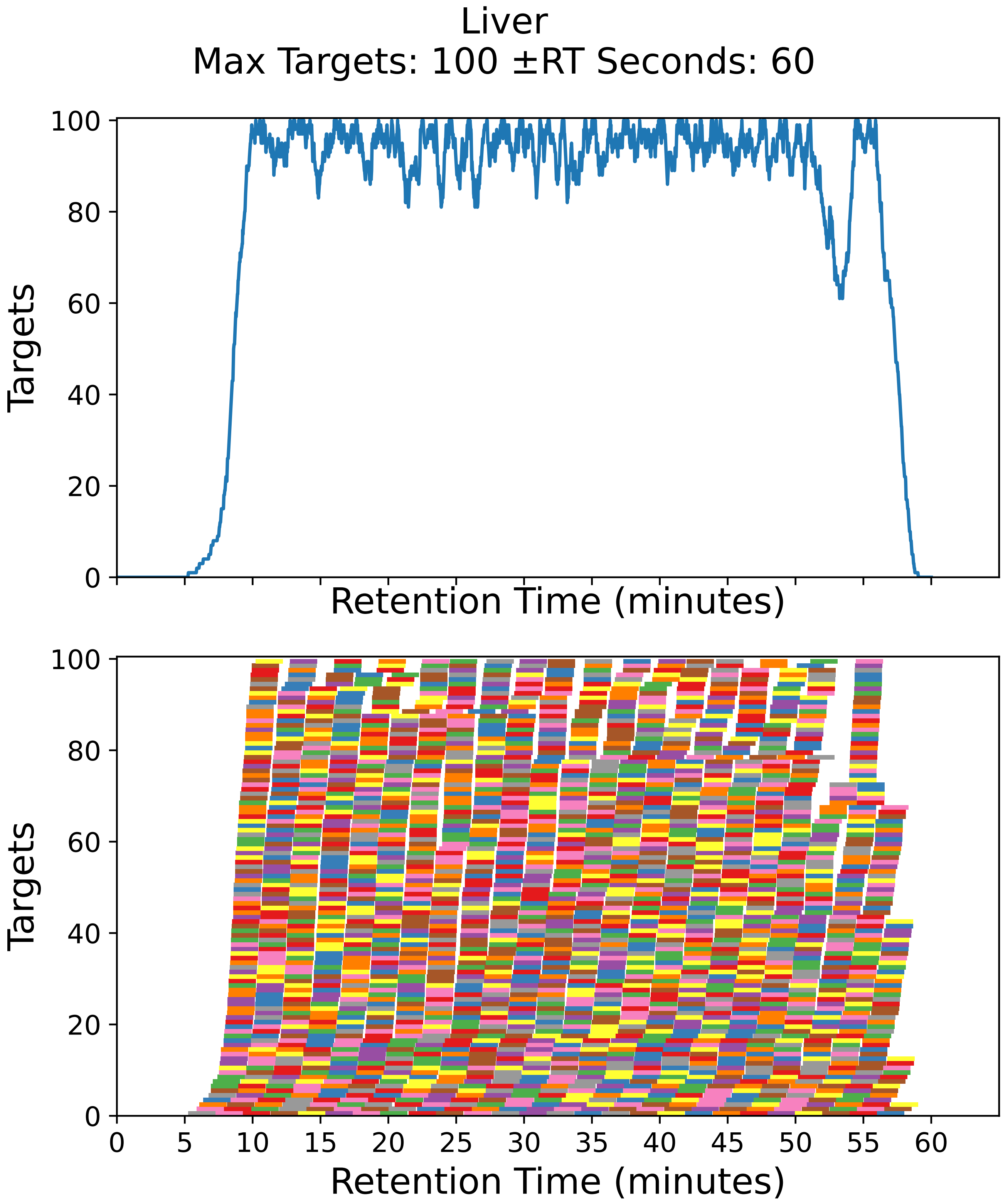

Supplementary Figure 41: Top: utilization diagram for the best fit method with instrument targeting the liver filtered subset with a maximum of 100 targets per cycle with retention times ± 60 seconds from the peptide peak apex. Bottom: individual targeting windows for the 2262 peptides, monitoring 1131 proteins.

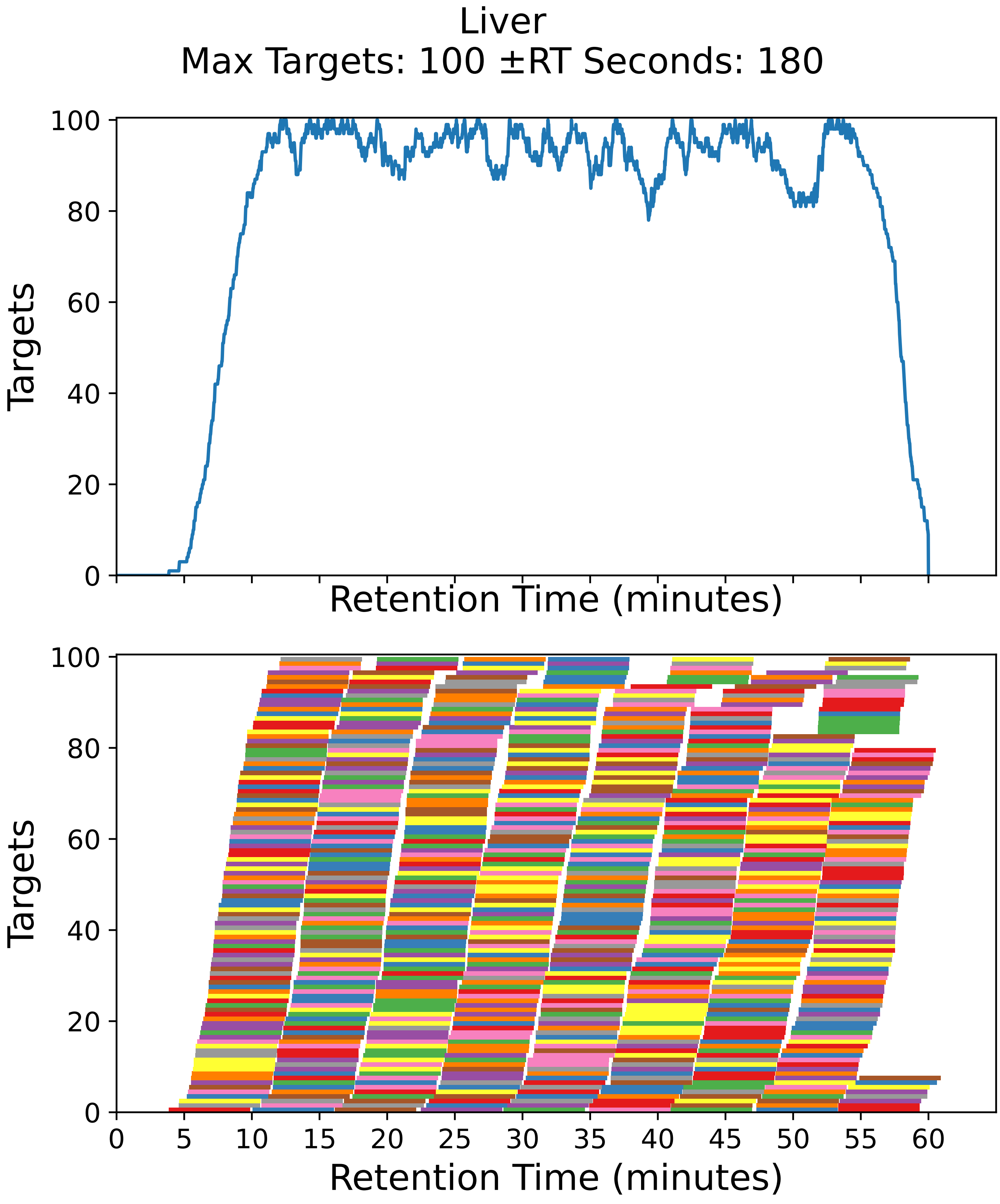

Supplementary Figure 42: Top: utilization diagram for the best fit method with instrument targeting the liver filtered subset with a maximum of 100 targets per cycle with retention times ± 180 seconds from the peptide peak apex. Bottom: individual targeting windows for the 784 peptides, monitoring 392 proteins.

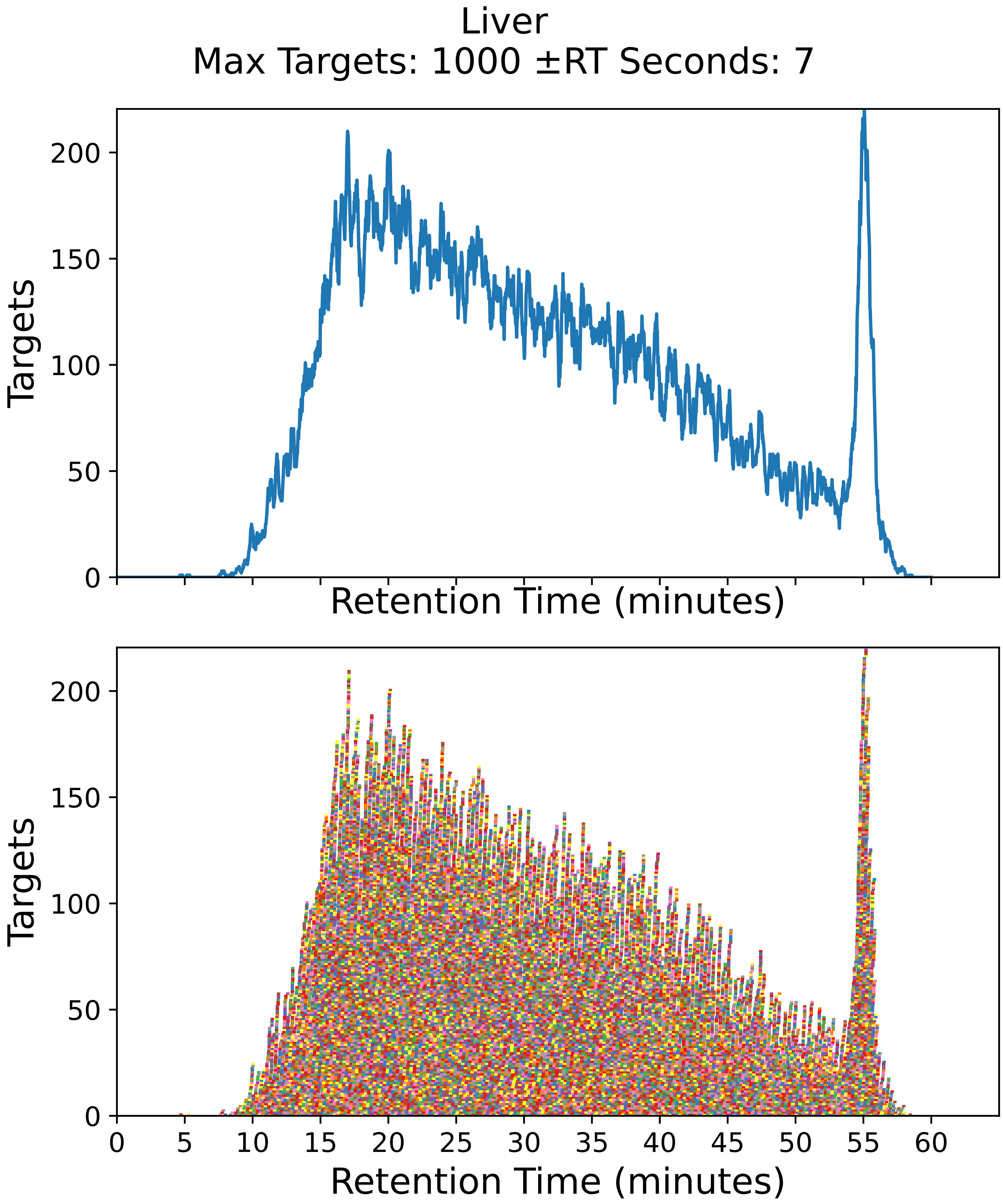

Supplementary Figure 43: Top: utilization diagram for the best fit method with instrument targeting the liver filtered subset with a maximum of 1000 targets per cycle with retention times ± 7 seconds from the peptide peak apex. Bottom: individual targeting windows for the 19462 peptides, monitoring 9731 proteins.

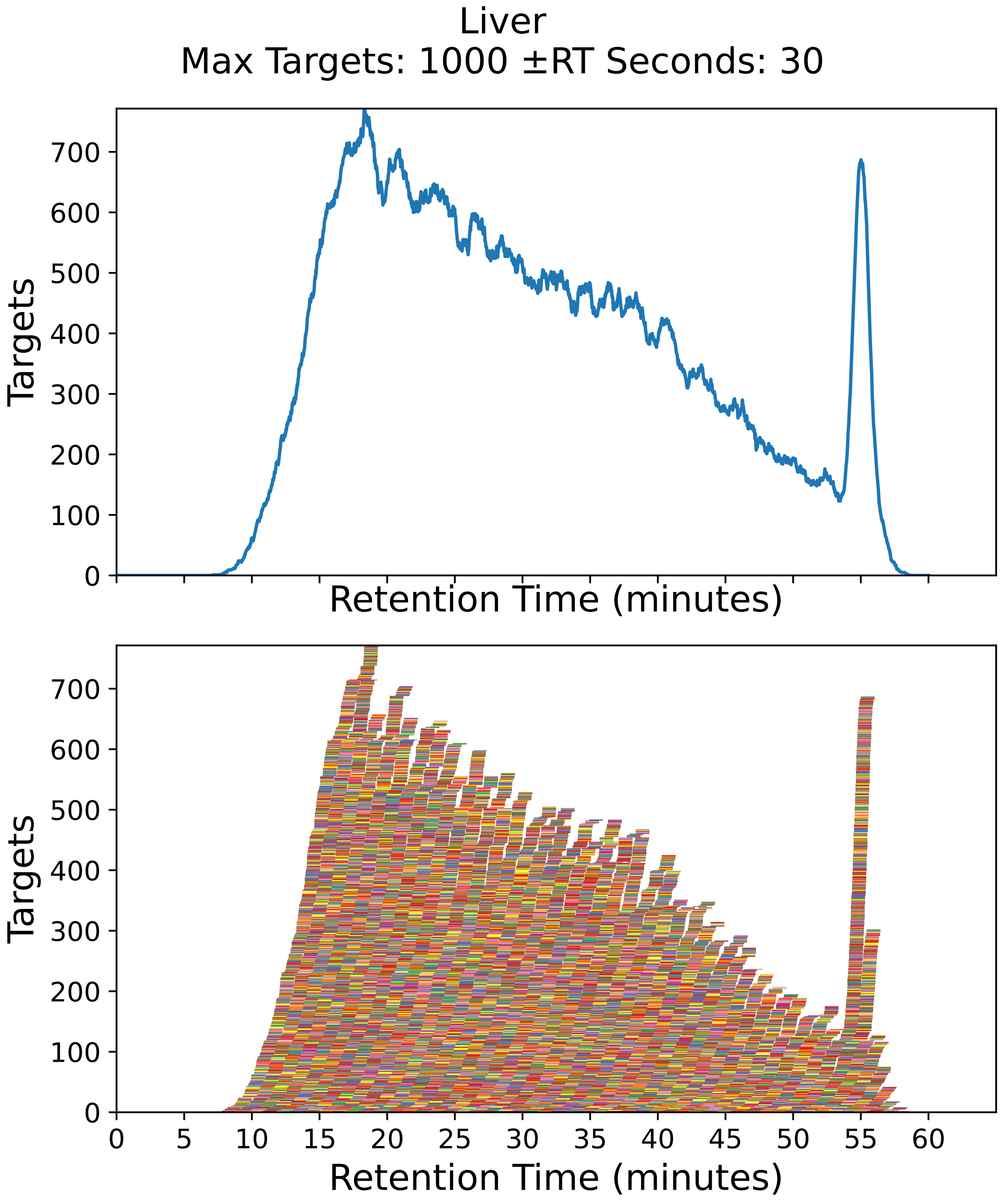

Supplementary Figure 44: Top: utilization diagram for the best fit method with instrument targeting the liver filtered subset with a maximum of 1000 targets per cycle with retention times ± 30 seconds from the peptide peak apex. Bottom: individual targeting windows for the 19462 peptides, monitoring 9731 proteins.

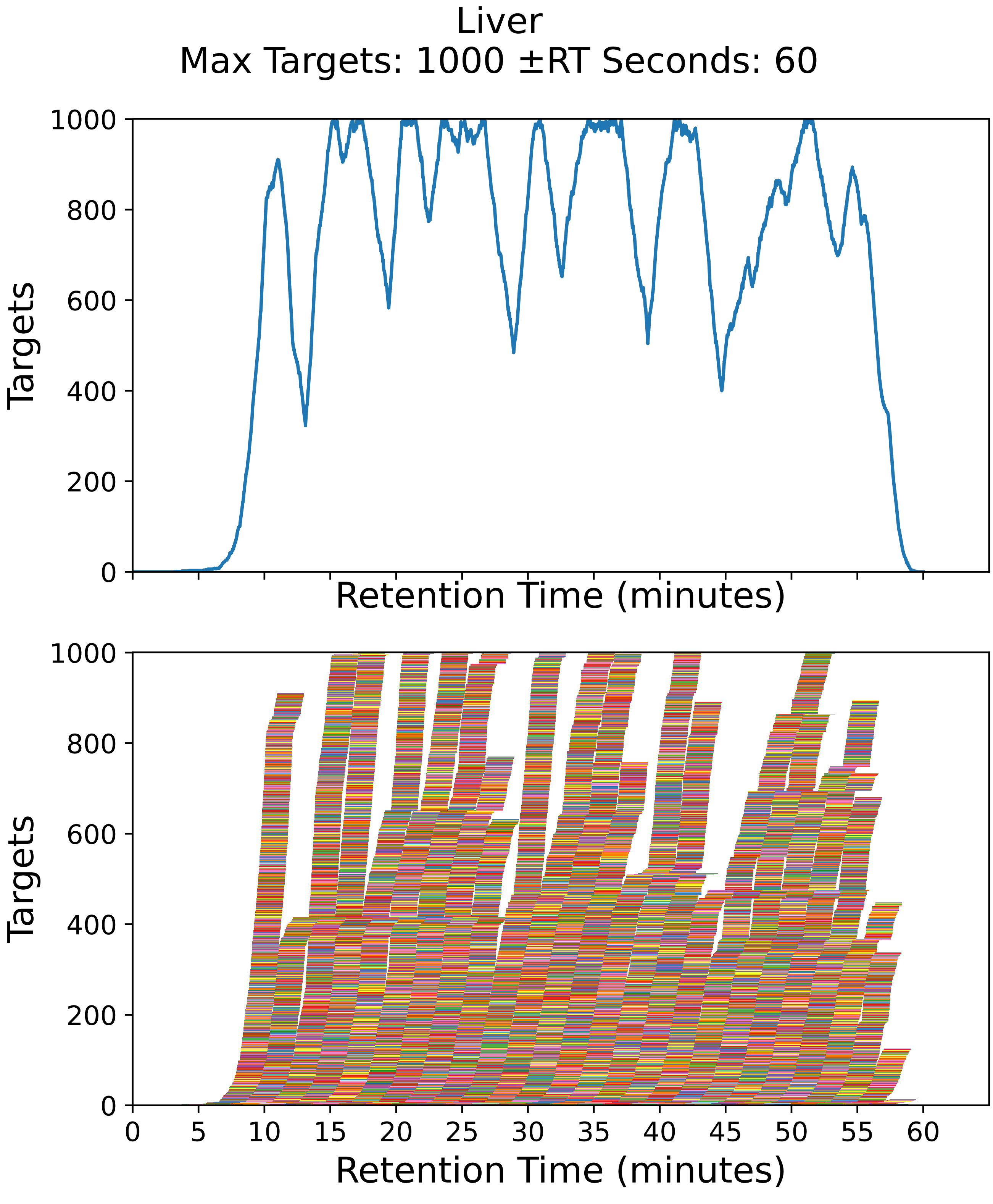

Supplementary Figure 45: Top: utilization diagram for the best fit method with instrument targeting the liver filtered subset with a maximum of 1000 targets per cycle with retention times ± 60 seconds from the peptide peak apex. Bottom: individual targeting windows for the 19462 peptides, monitoring 9731 proteins.

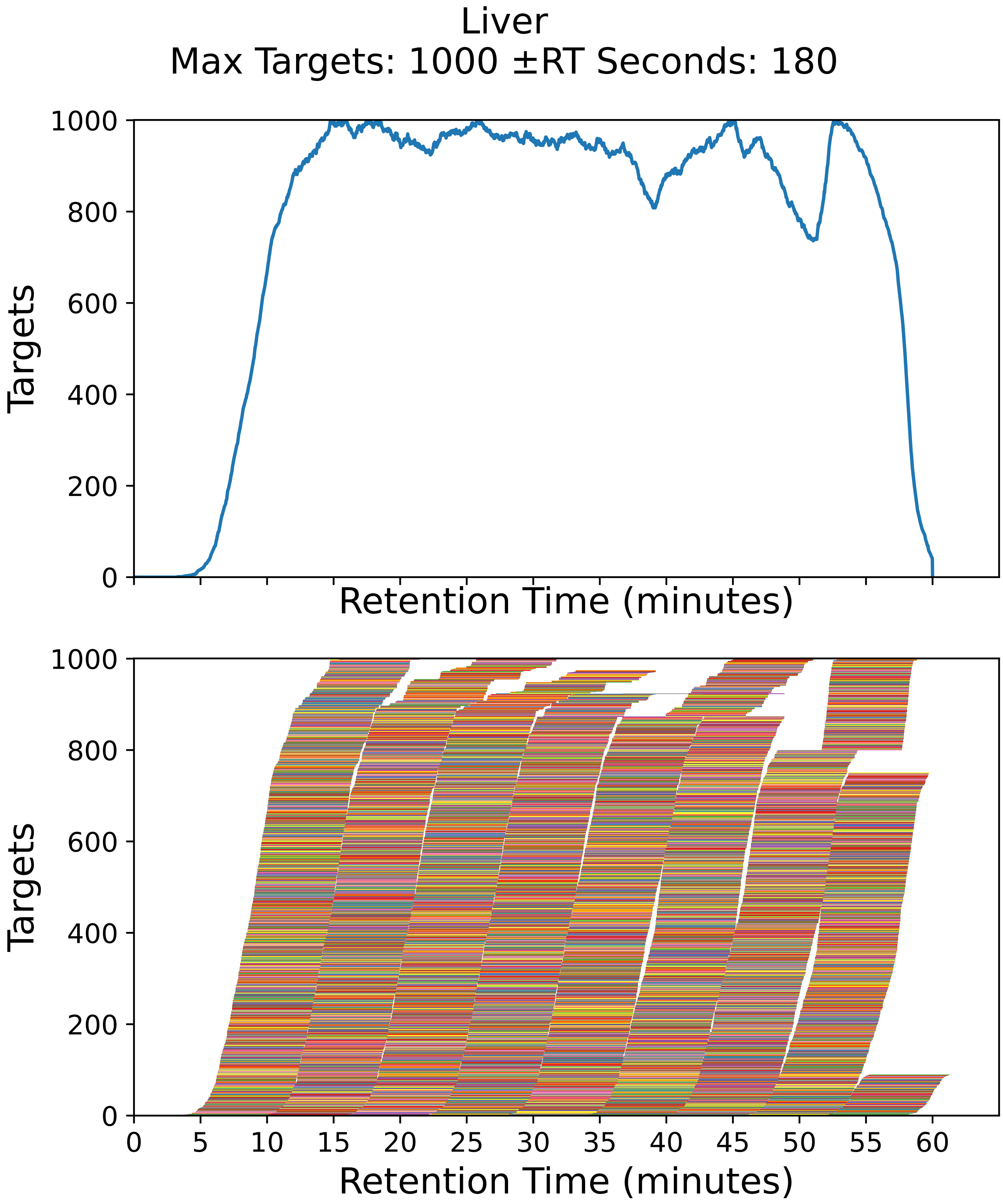

Supplementary Figure 46: Top: utilization diagram for the best fit method with instrument targeting the liver filtered subset with a maximum of 1000 targets per cycle with retention times ± 180 seconds from the peptide peak apex. Bottom: individual targeting windows for the 7612 peptides, monitoring 3806 proteins.

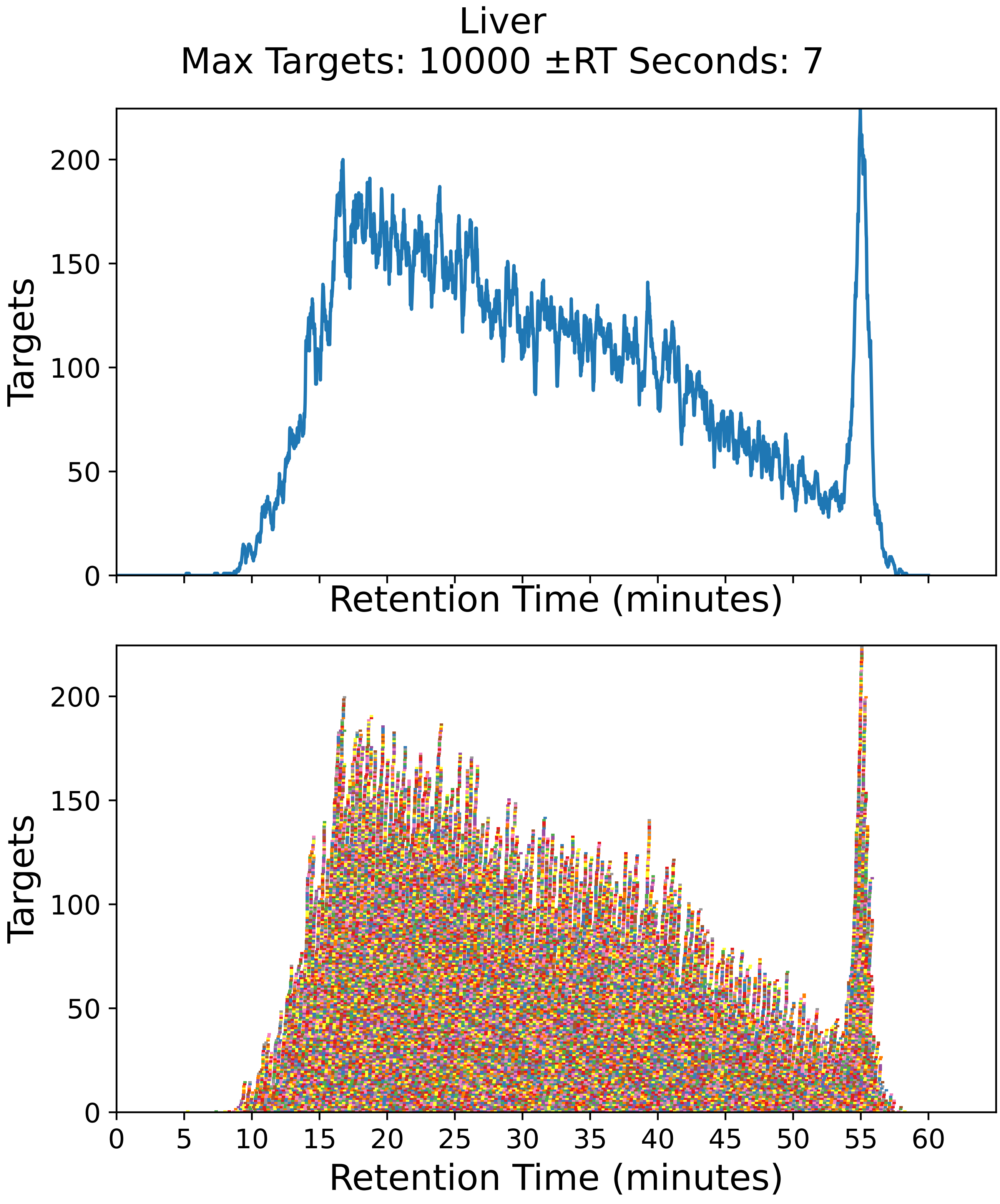

Supplementary Figure 47: Top: utilization diagram for the best fit method with instrument targeting the liver filtered subset with a maximum of 10000 targets per cycle with retention times ± 7 seconds from the peptide peak apex. Bottom: individual targeting windows for the 19462 peptides, monitoring 9731 proteins.

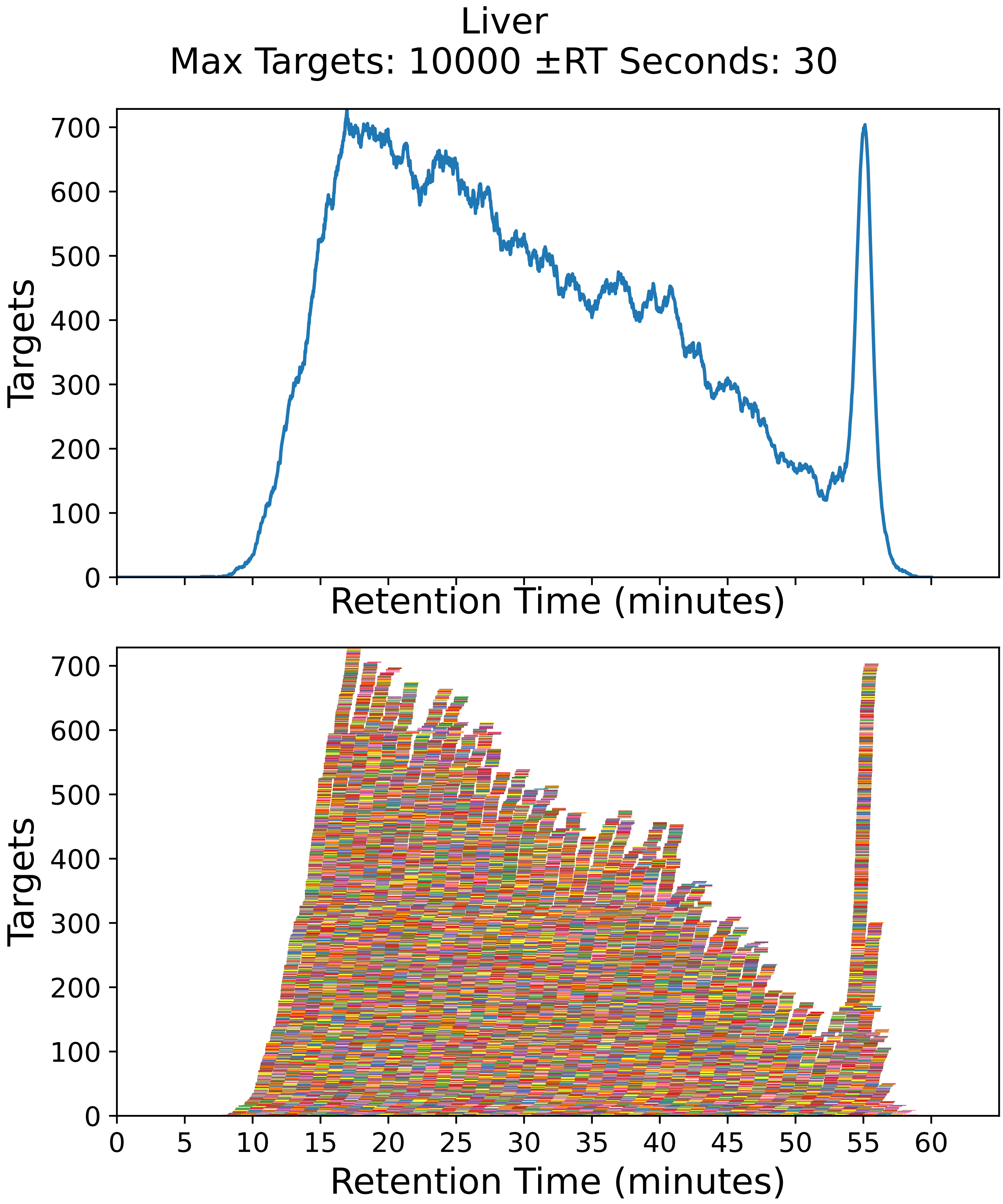

Supplementary Figure 48: Top: utilization diagram for the best fit method with instrument targeting the liver filtered subset with a maximum of 10000 targets per cycle with retention times ± 30 seconds from the peptide peak apex. Bottom: individual targeting windows for the 19462 peptides, monitoring 9731 proteins.

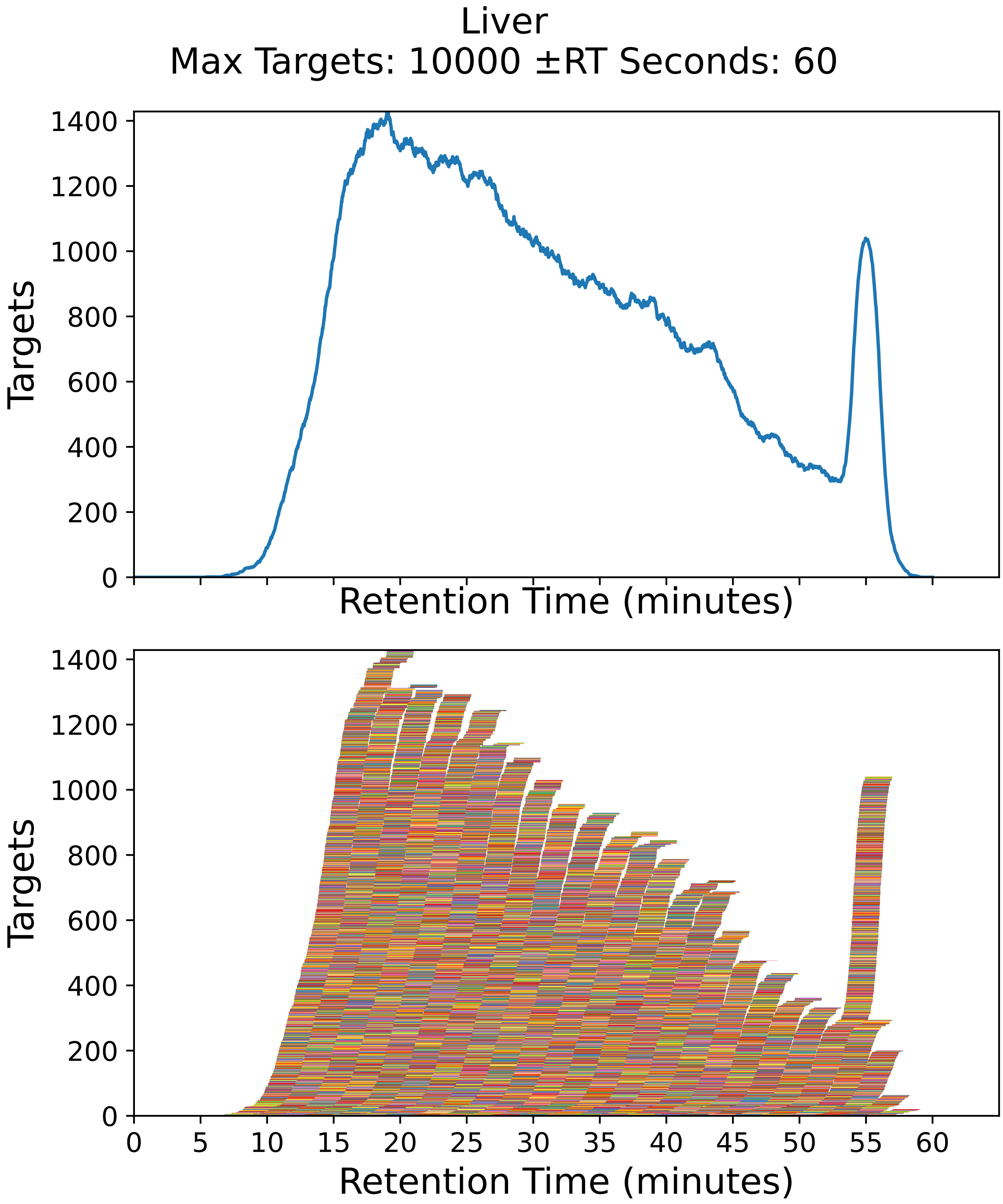

Supplementary Figure 49: Top: utilization diagram for the best fit method with instrument targeting the liver filtered subset with a maximum of 10000 targets per cycle with retention times ± 60 seconds from the peptide peak apex. Bottom: individual targeting windows for the 19462 peptides, monitoring 9731 proteins.

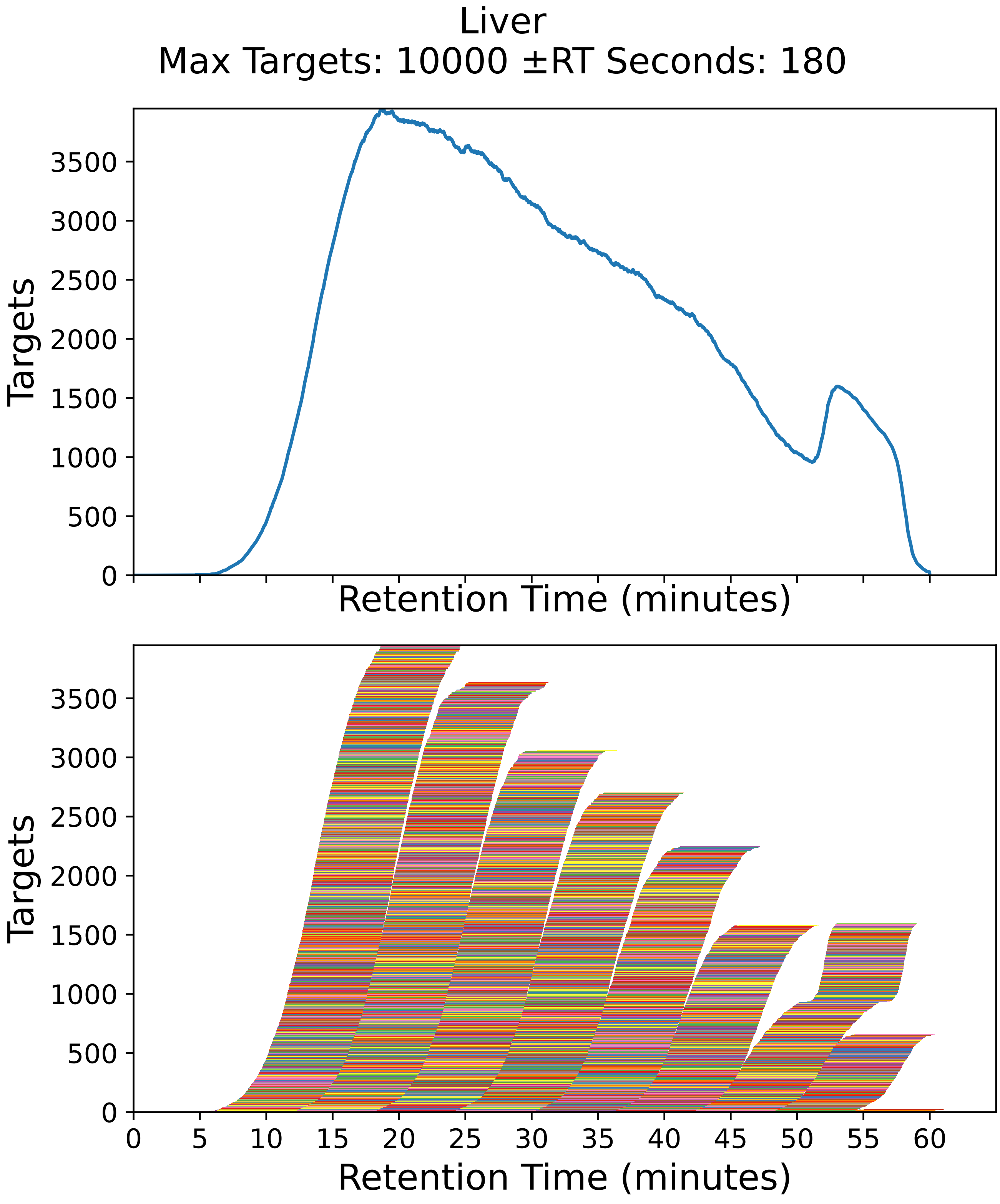

Supplementary Figure 50: Top: utilization diagram for the best fit method with instrument targeting the liver filtered subset with a maximum of 10000 targets per cycle with retention times ± 180 seconds from the peptide peak apex. Bottom: individual targeting windows for the 19462 peptides, monitoring 9731 proteins.

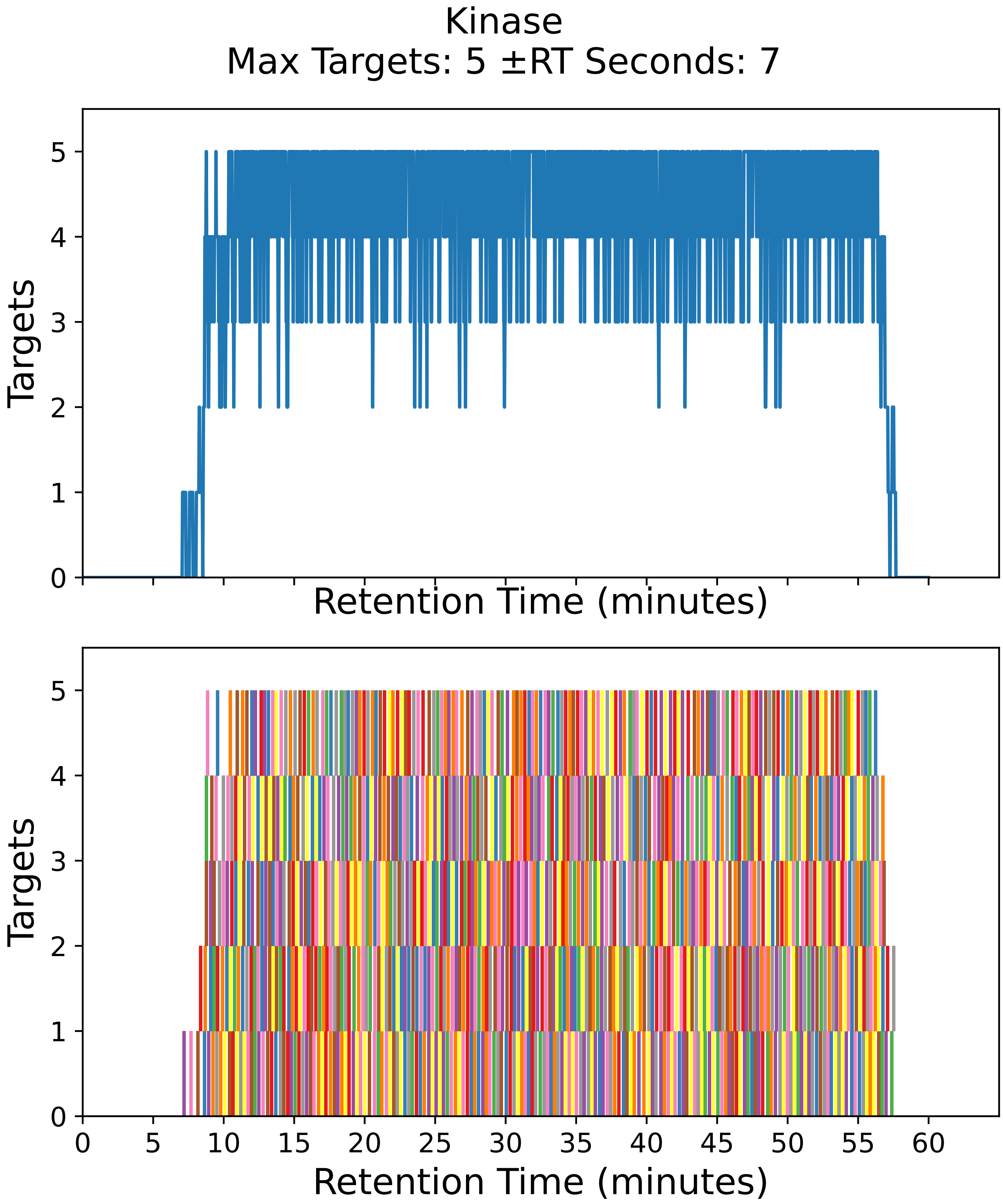

Supplementary Figure 51: Top: utilization diagram for the best fit method with instrument targeting the kinase filtered subset with a maximum of 5 targets per cycle with retention times ± 7 seconds from the peptide peak apex. Bottom: individual targeting windows for the 842 peptides, monitoring 421 proteins.

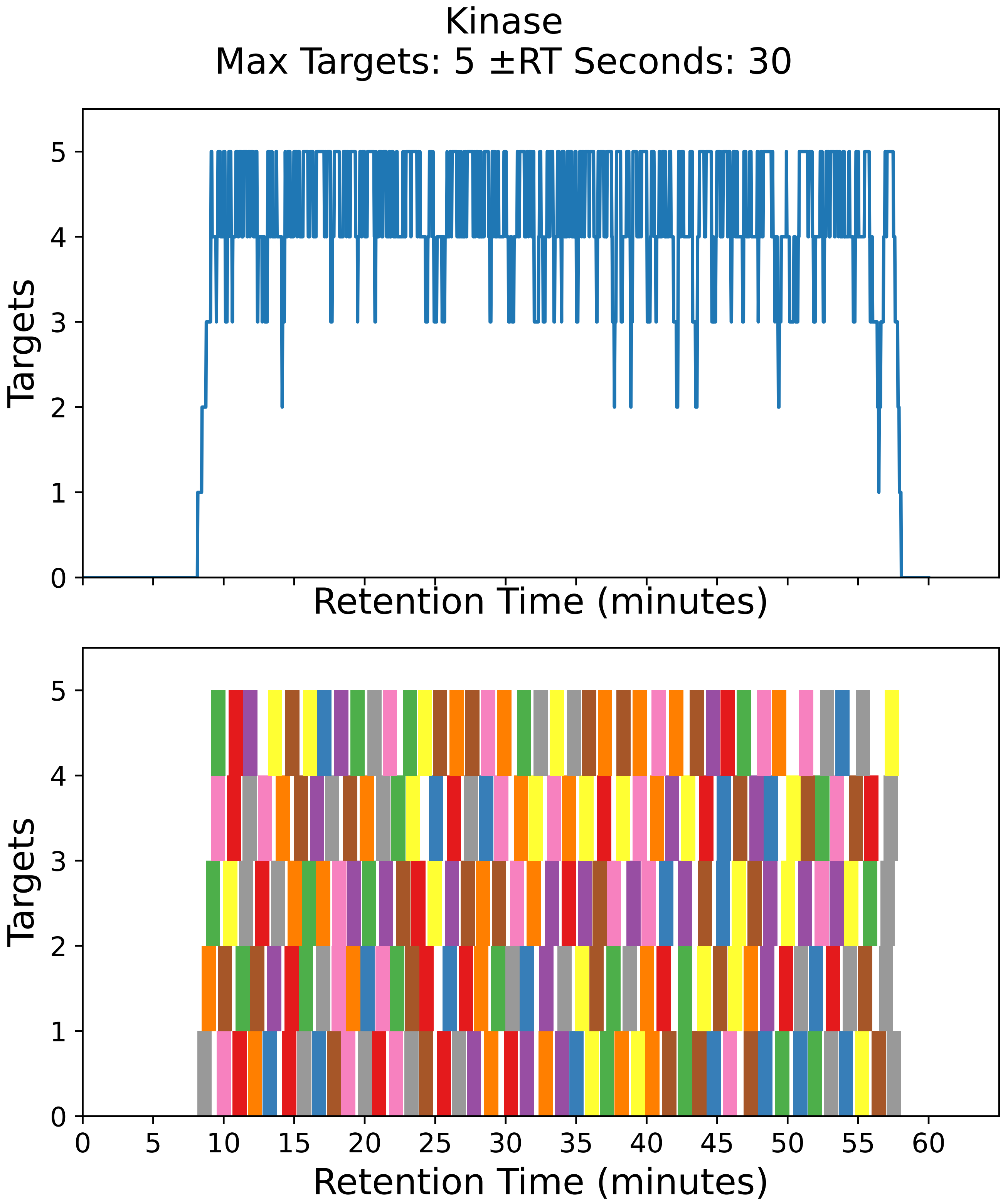

Supplementary Figure 52: Top: utilization diagram for the best fit method with instrument targeting the kinase filtered subset with a maximum of 5 targets per cycle with retention times ± 30 seconds from the peptide peak apex. Bottom: individual targeting windows for the 208 peptides, monitoring 104 proteins.

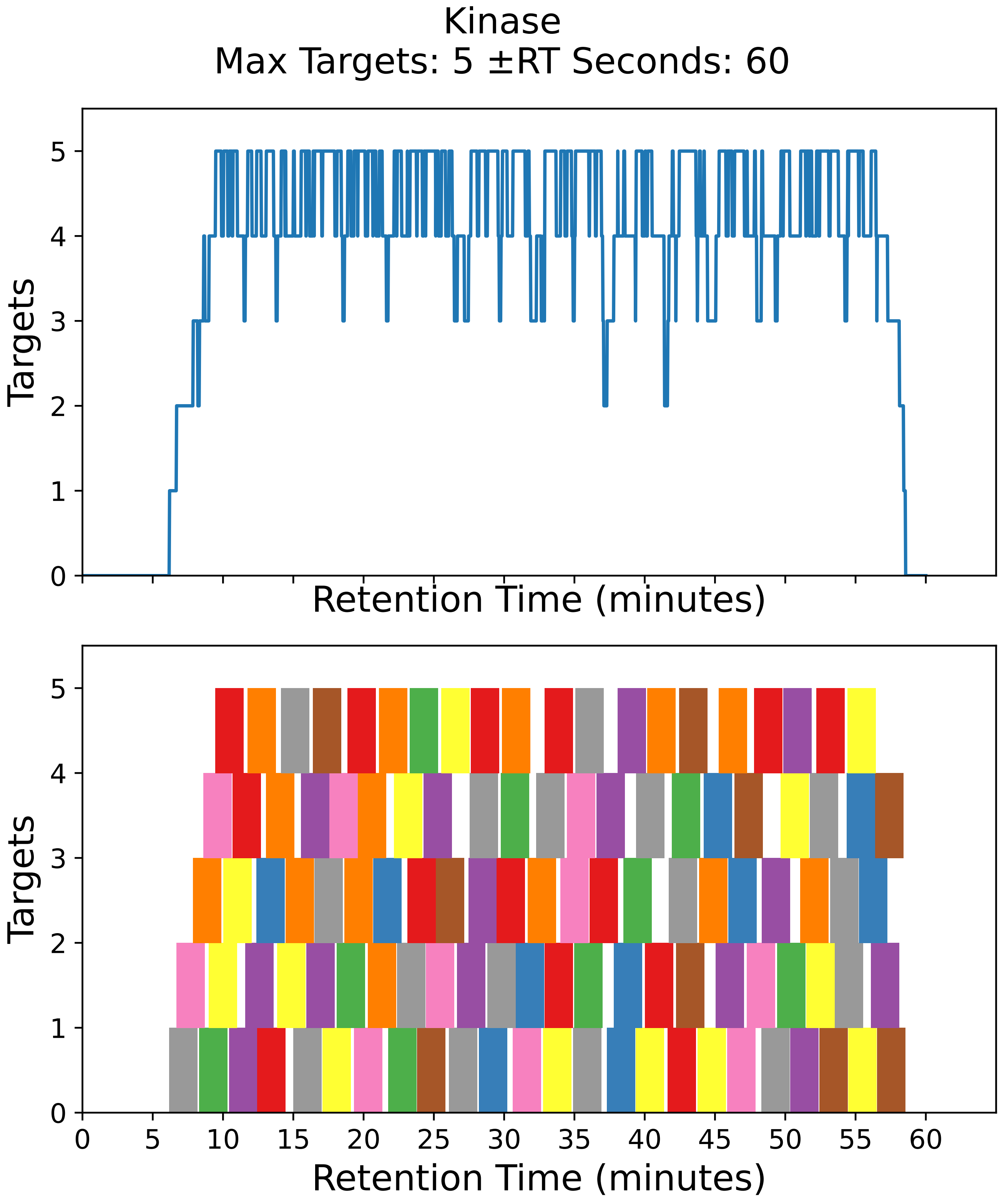

Supplementary Figure 53: Top: utilization diagram for the best fit method with instrument targeting the kinase filtered subset with a maximum of 5 targets per cycle with retention times ± 60 seconds from the peptide peak apex. Bottom: individual targeting windows for the 110 peptides, monitoring 55 proteins.

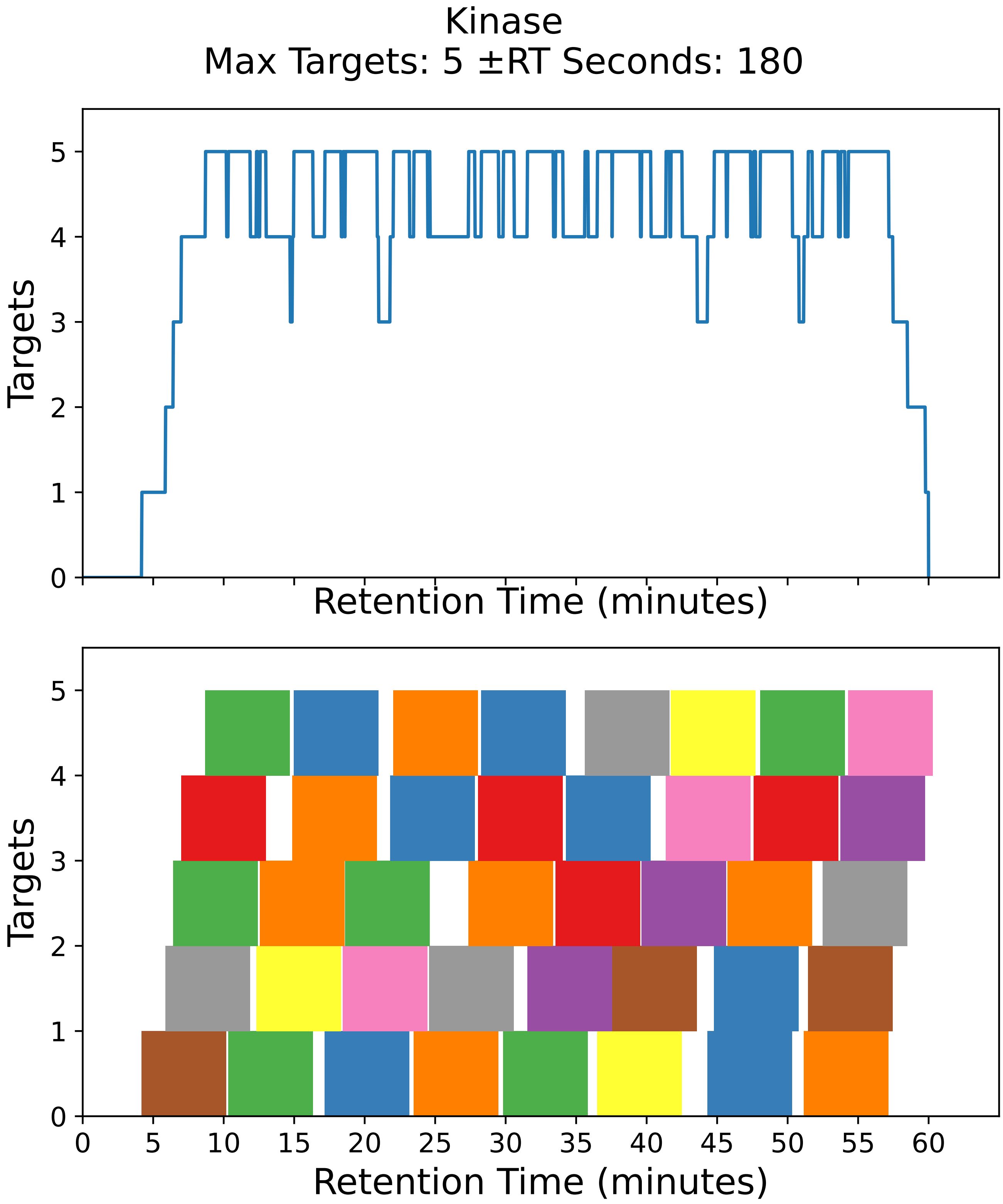

Supplementary Figure 54: Top: utilization diagram for the best fit method with instrument targeting the kinase filtered subset with a maximum of 5 targets per cycle with retention times ± 180 seconds from the peptide peak apex. Bottom: individual targeting windows for the 40 peptides, monitoring 20 proteins.

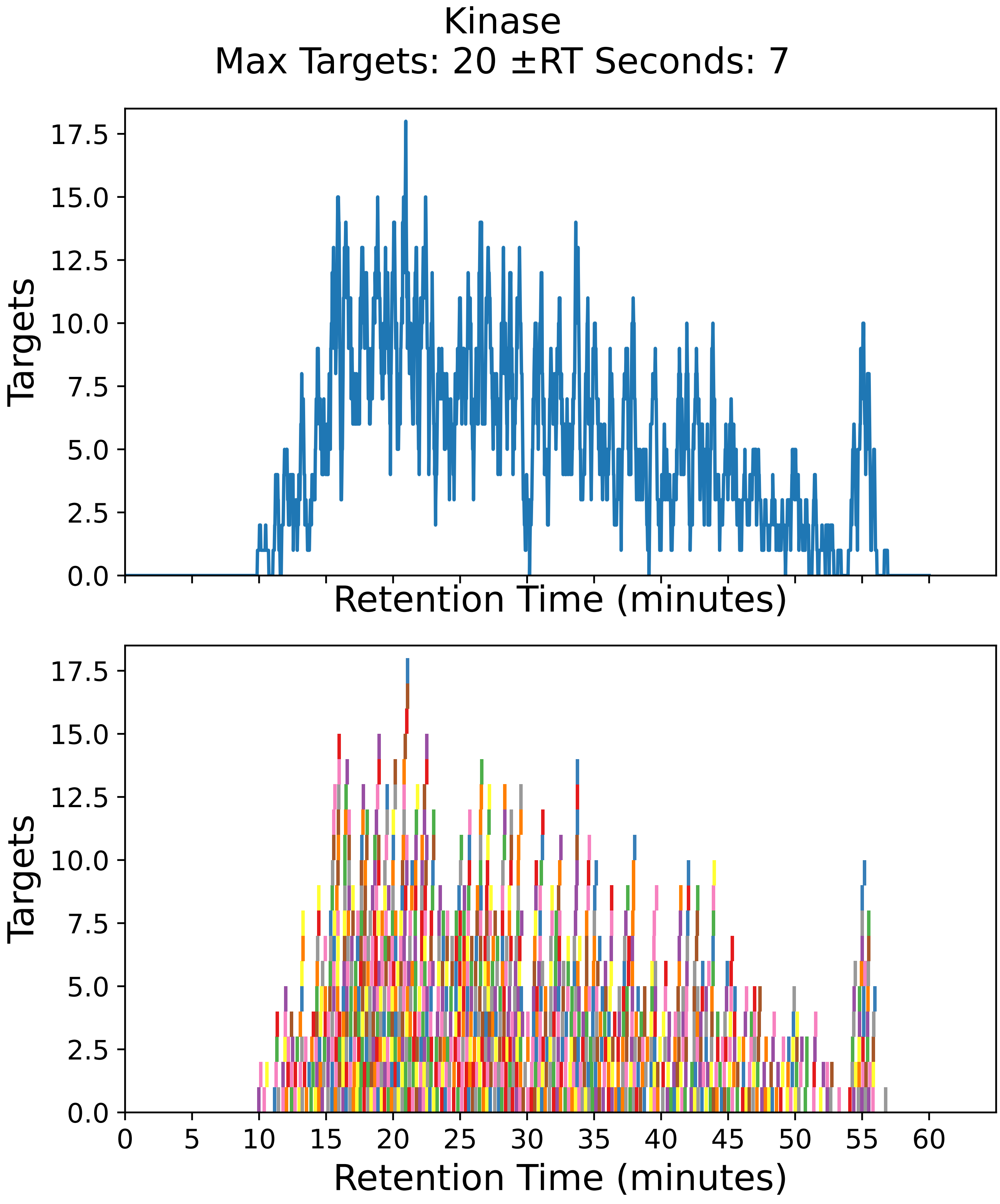

Supplementary Figure 55: Top: utilization diagram for the best fit method with instrument targeting the kinase filtered subset with a maximum of 20 targets per cycle with retention times ± 7 seconds from the peptide peak apex. Bottom: individual targeting windows for the 1022 peptides, monitoring 511 proteins.

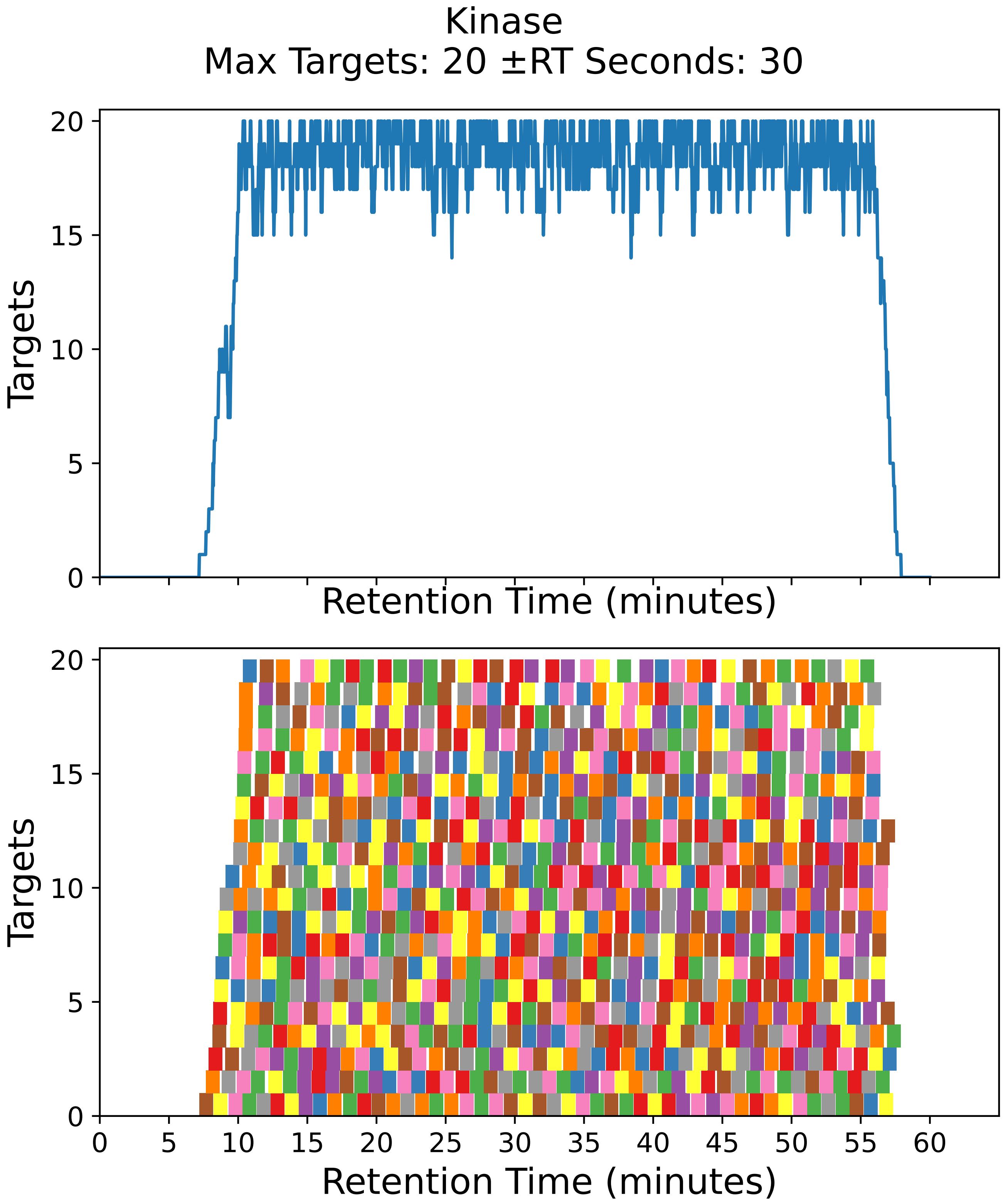

Supplementary Figure 56: Top: utilization diagram for the best fit method with instrument targeting the kinase filtered subset with a maximum of 20 targets per cycle with retention times ± 30 seconds from the peptide peak apex. Bottom: individual targeting windows for the 870 peptides, monitoring 435 proteins.

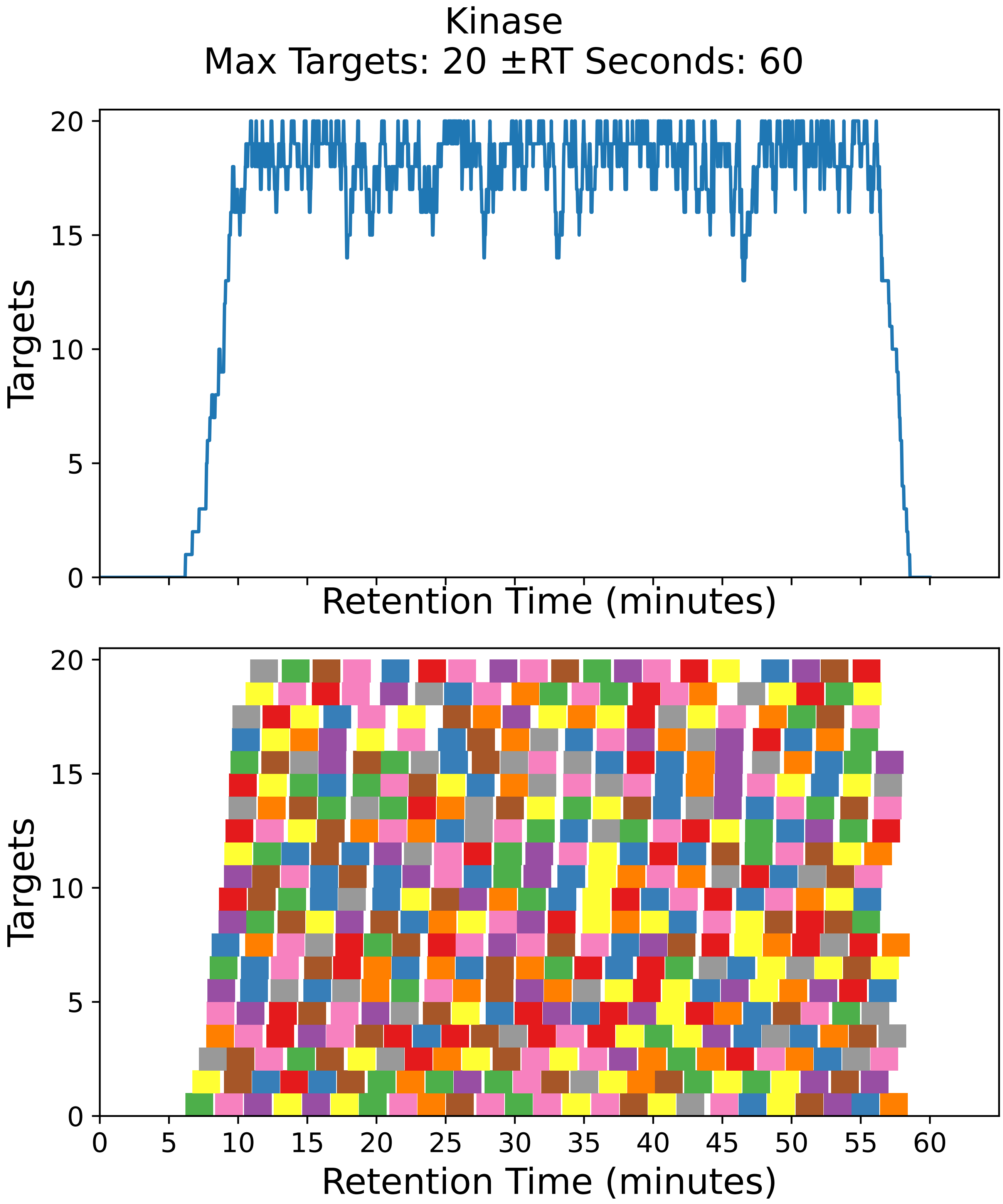

Supplementary Figure 57: Top: utilization diagram for the best fit method with instrument targeting the kinase filtered subset with a maximum of 20 targets per cycle with retention times ± 60 seconds from the peptide peak apex. Bottom: individual targeting windows for the 444 peptides, monitoring 222 proteins.

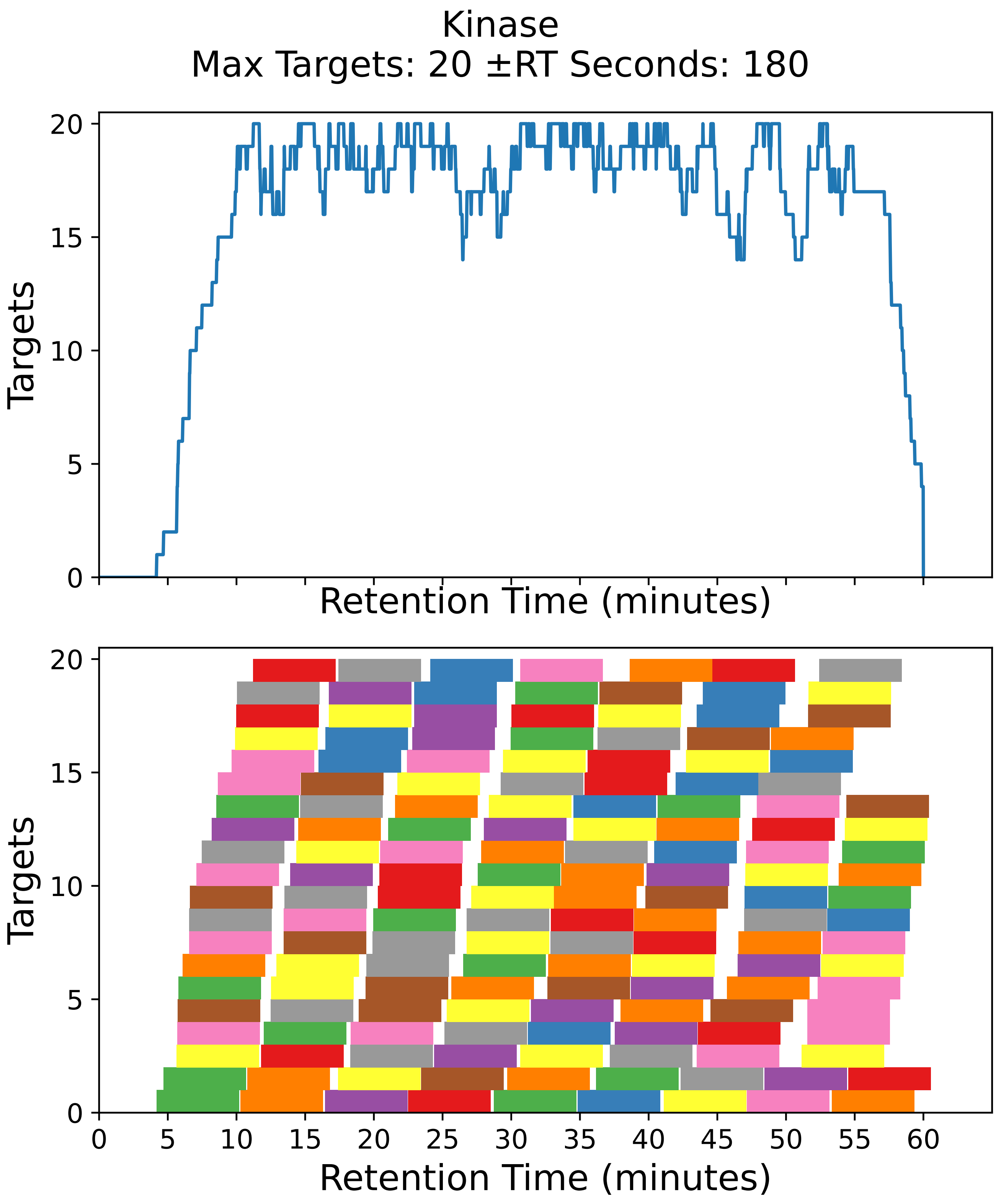

Supplementary Figure 58: Top: utilization diagram for the best fit method with instrument targeting the kinase filtered subset with a maximum of 20 targets per cycle with retention times ± 180 seconds from the peptide peak apex. Bottom: individual targeting windows for the 156 peptides, monitoring 78 proteins.

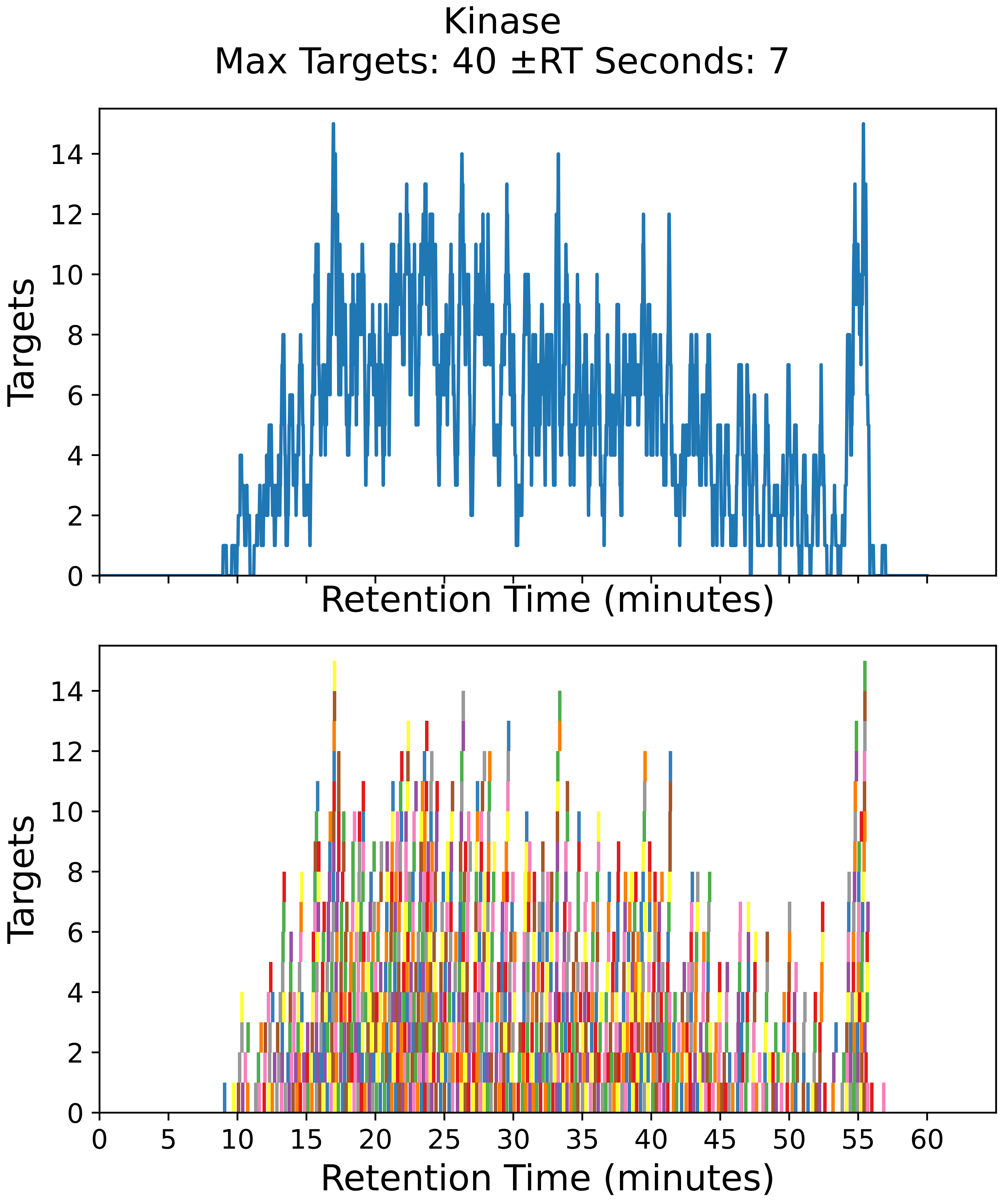

Supplementary Figure 59: Top: utilization diagram for the best fit method with instrument targeting the kinase filtered subset with a maximum of 40 targets per cycle with retention times ± 7 seconds from the peptide peak apex. Bottom: individual targeting windows for the 1022 peptides, monitoring 511 proteins.

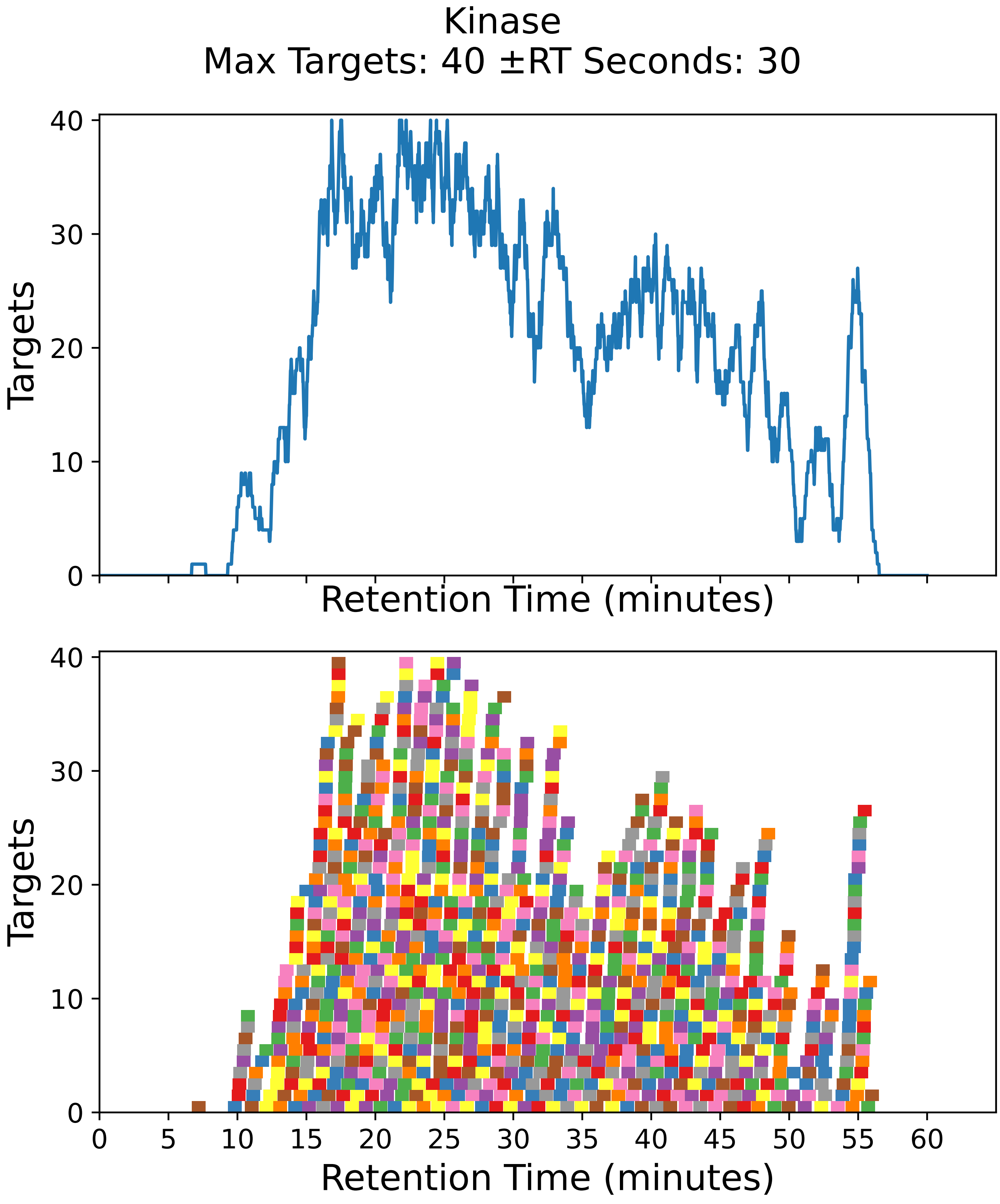

Supplementary Figure 60: Top: utilization diagram for the best fit method with instrument targeting the kinase filtered subset with a maximum of 40 targets per cycle with retention times ± 30 seconds from the peptide peak apex. Bottom: individual targeting windows for the 1022 peptides, monitoring 511 proteins.

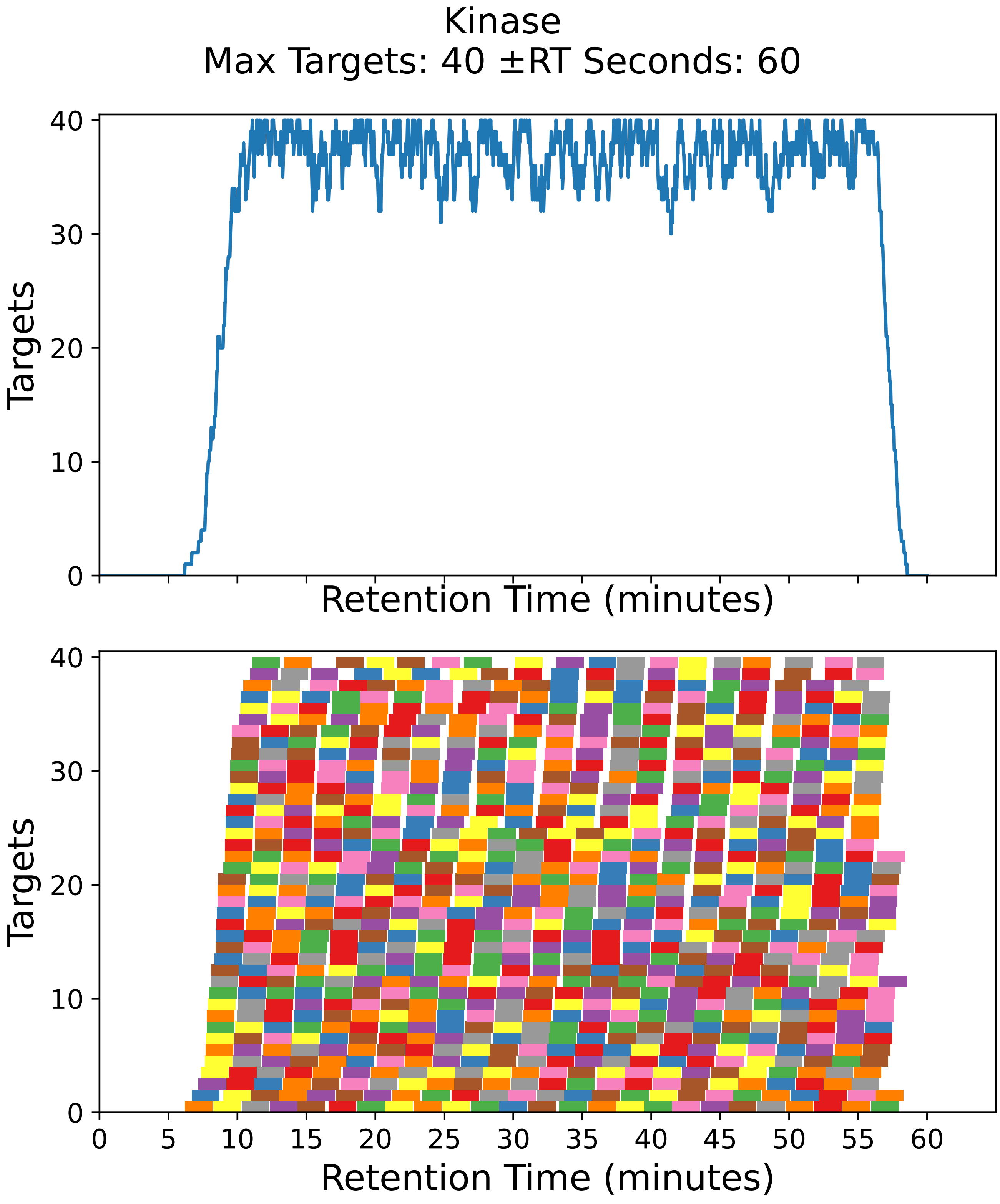

Supplementary Figure 61: Top: utilization diagram for the best fit method with instrument targeting the kinase filtered subset with a maximum of 40 targets per cycle with retention times ± 60 seconds from the peptide peak apex. Bottom: individual targeting windows for the 896 peptides, monitoring 448 proteins.

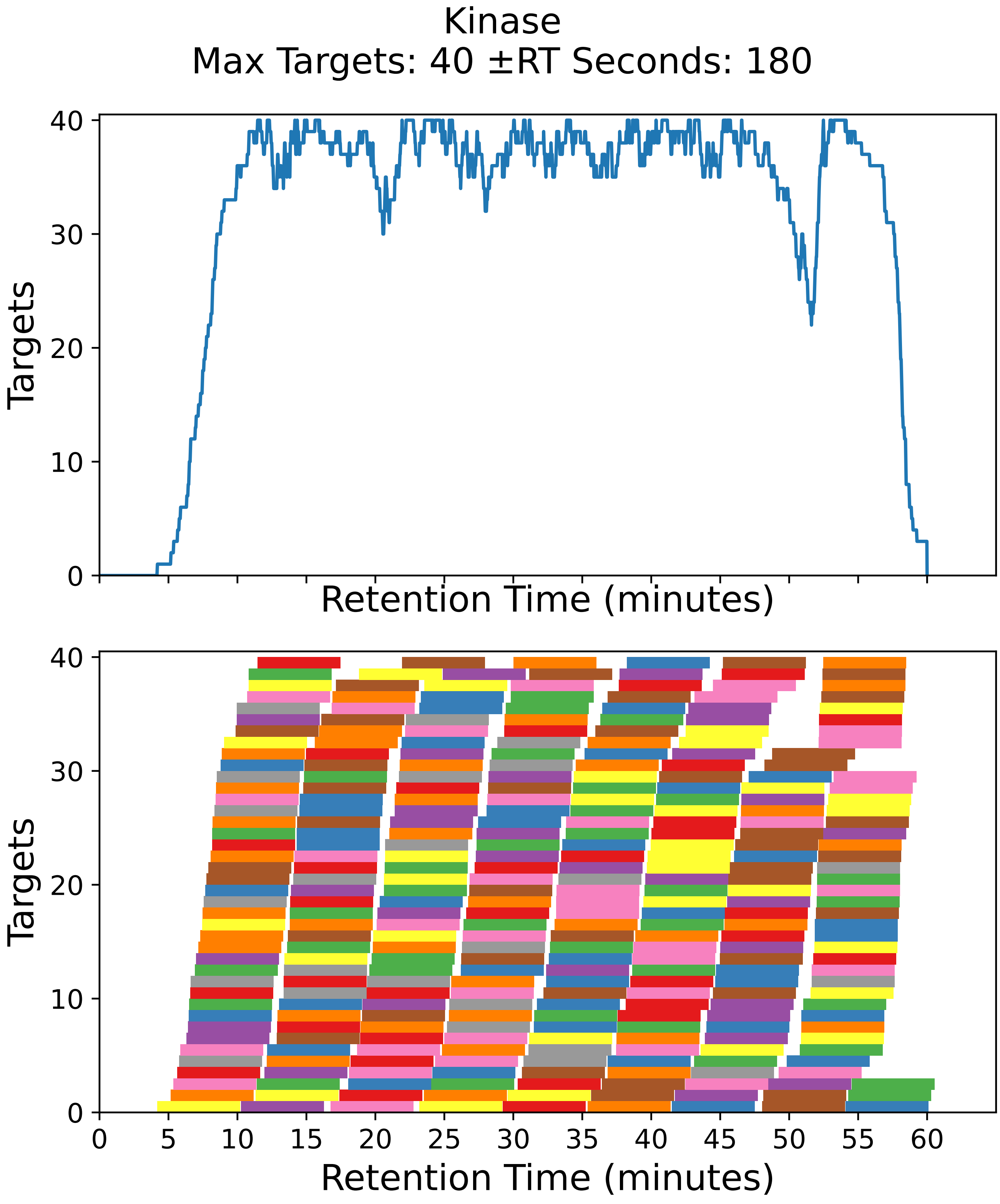

Supplementary Figure 62: Top: utilization diagram for the best fit method with instrument targeting the kinase filtered subset with a maximum of 40 targets per cycle with retention times ± 180 seconds from the peptide peak apex. Bottom: individual targeting windows for the 312 peptides, monitoring 156 proteins.

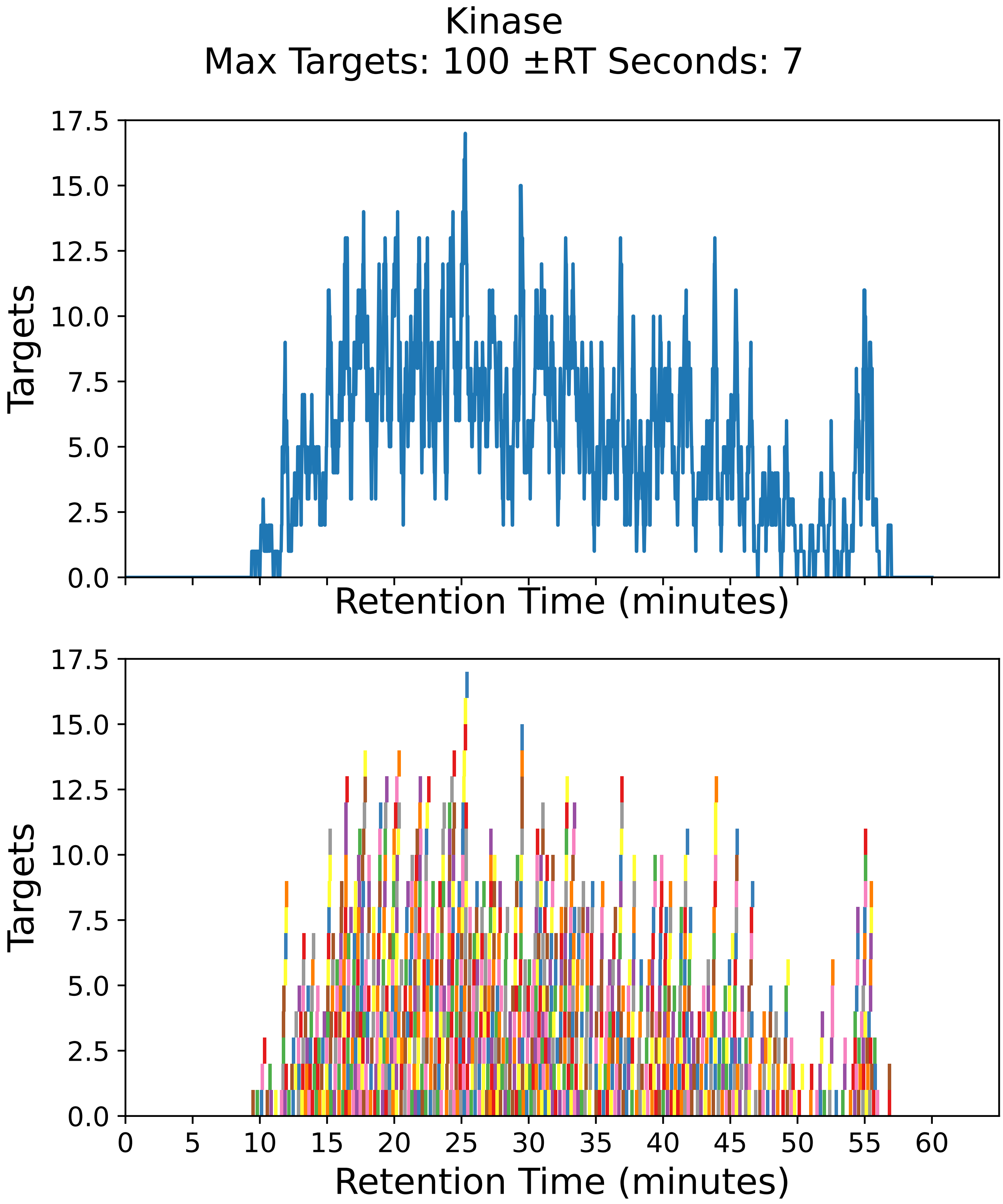

Supplementary Figure 63: Top: utilization diagram for the best fit method with instrument targeting the kinase filtered subset with a maximum of 100 targets per cycle with retention times ± 7 seconds from the peptide peak apex. Bottom: individual targeting windows for the 1022 peptides, monitoring 511 proteins.

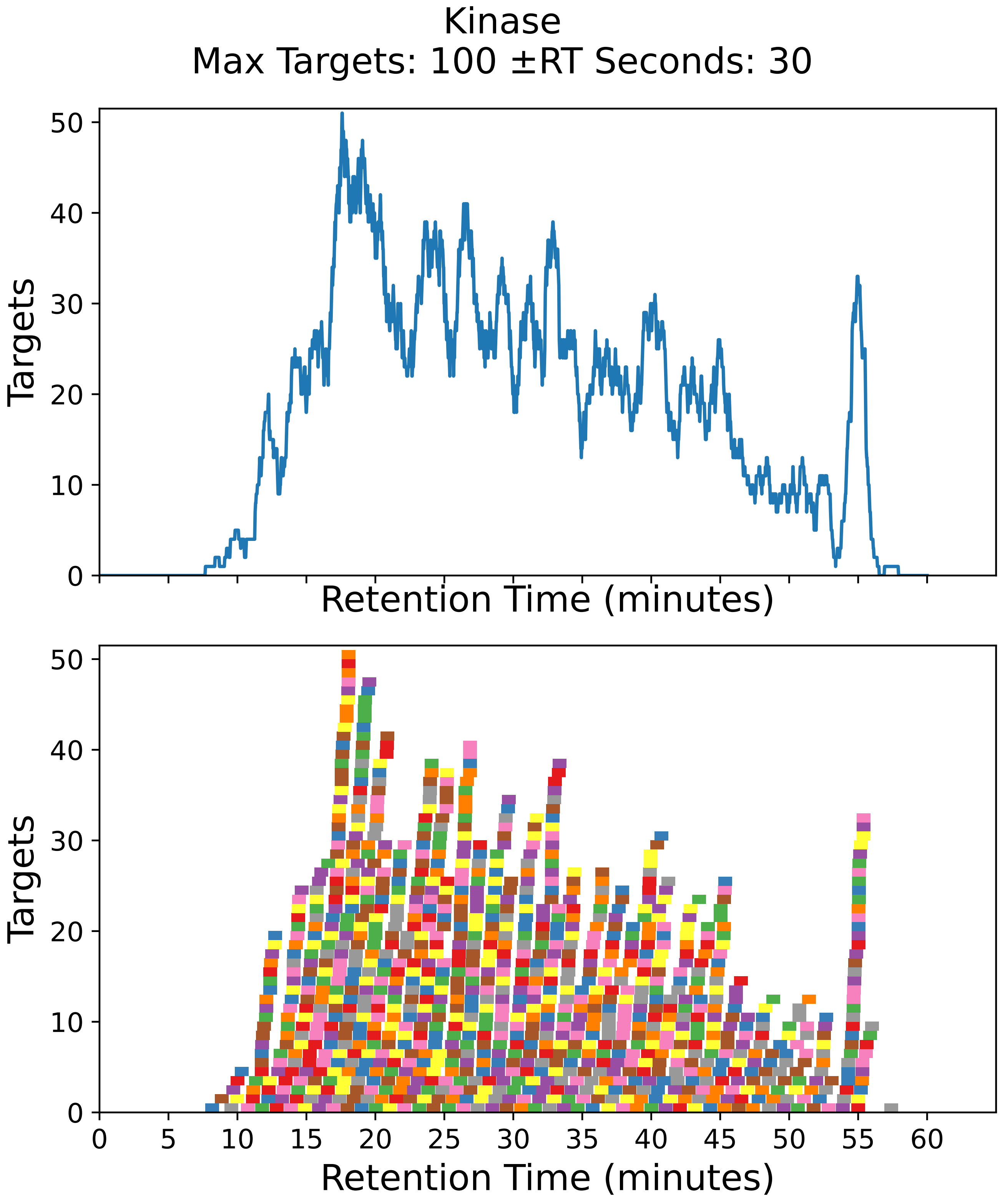

Supplementary Figure 64: Top: utilization diagram for the best fit method with instrument targeting the kinase filtered subset with a maximum of 100 targets per cycle with retention times ± 30 seconds from the peptide peak apex. Bottom: individual targeting windows for the 1022 peptides, monitoring 511 proteins.

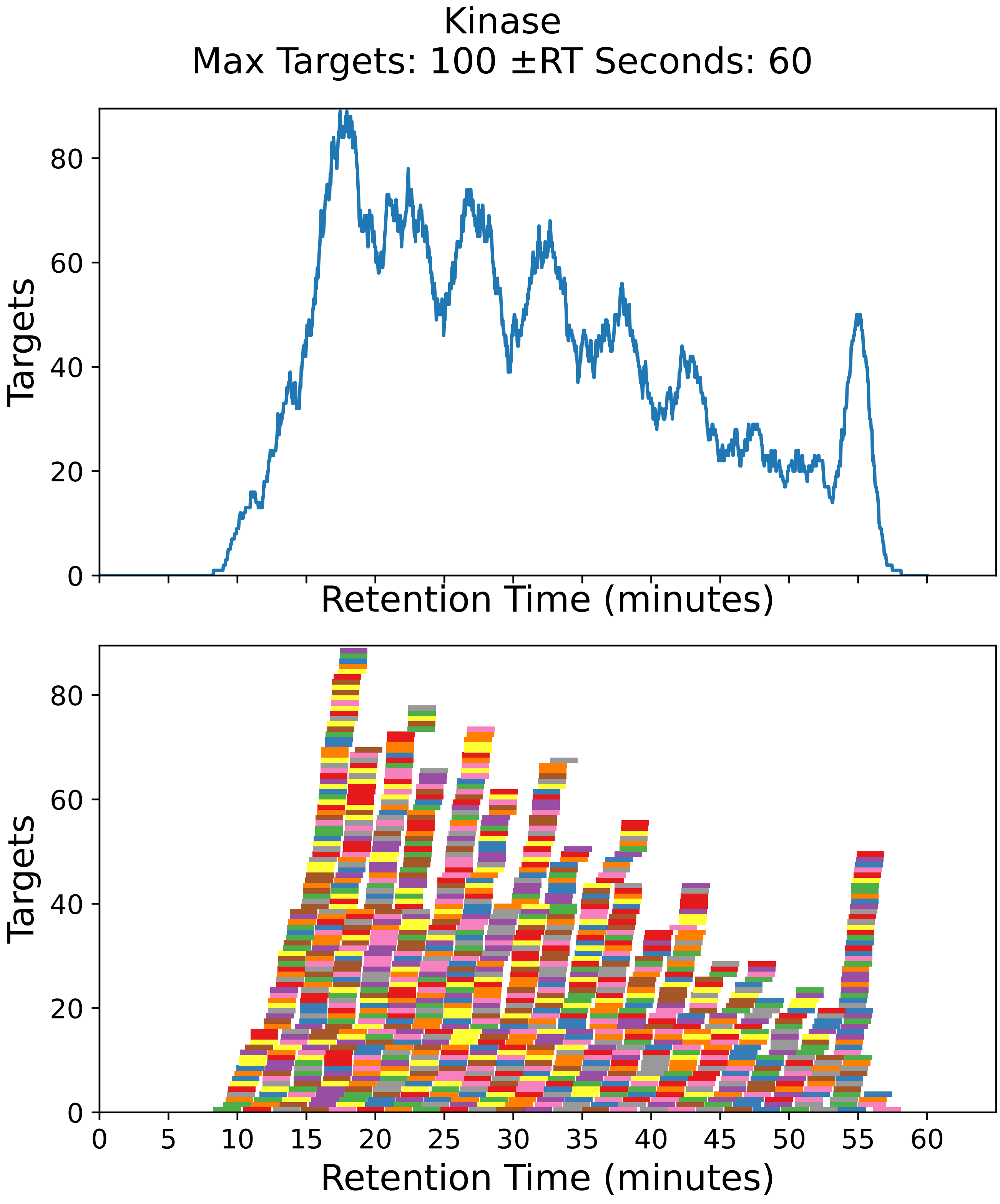

Supplementary Figure 65: Top: utilization diagram for the best fit method with instrument targeting the kinase filtered subset with a maximum of 100 targets per cycle with retention times ± 60 seconds from the peptide peak apex. Bottom: individual targeting windows for the 1022 peptides, monitoring 511 proteins.

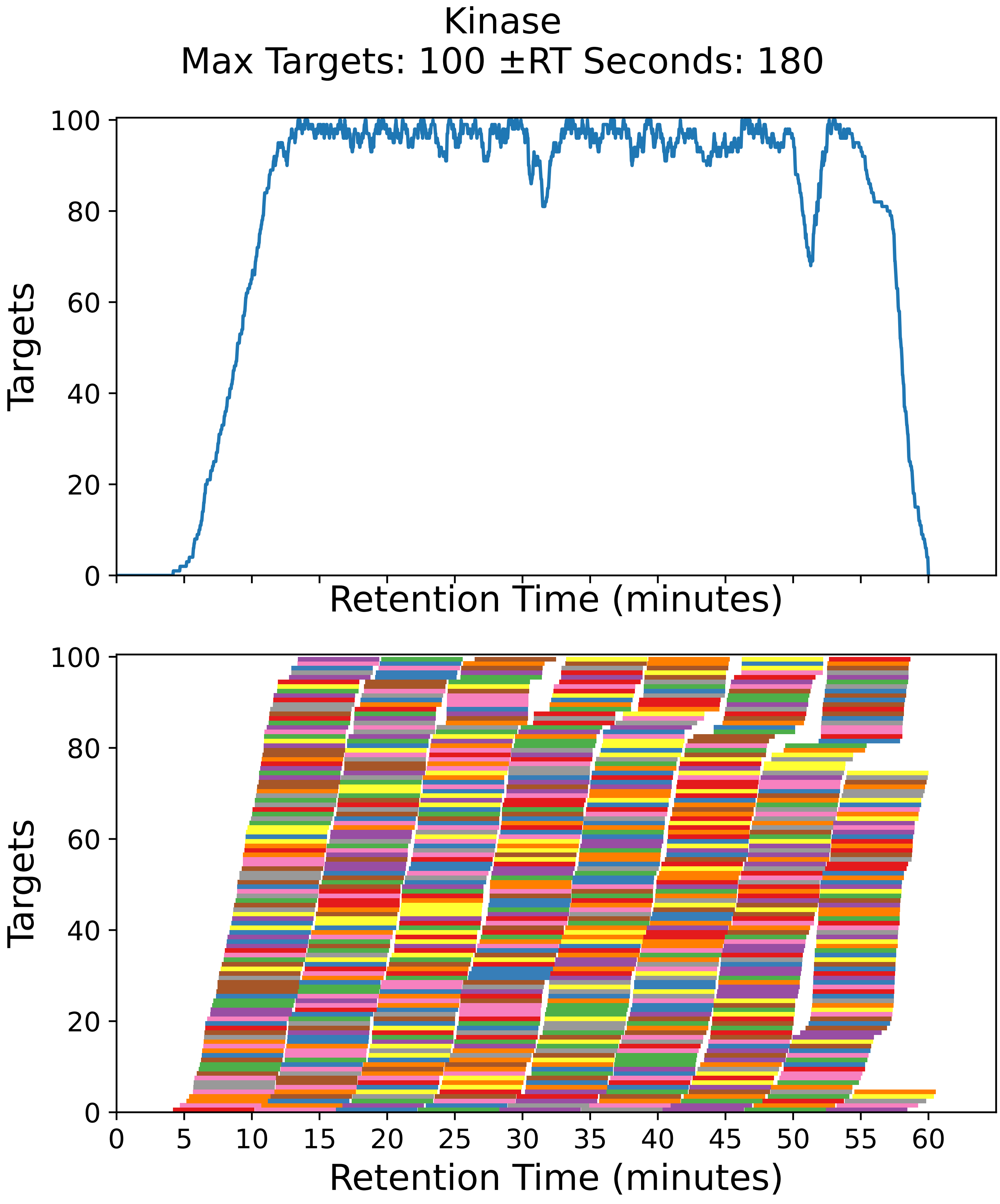

Supplementary Figure 66: Top: utilization diagram for the best fit method with instrument targeting the kinase filtered subset with a maximum of 100 targets per cycle with retention times ± 180 seconds from the peptide peak apex. Bottom: individual targeting windows for the 780 peptides, monitoring 390 proteins.

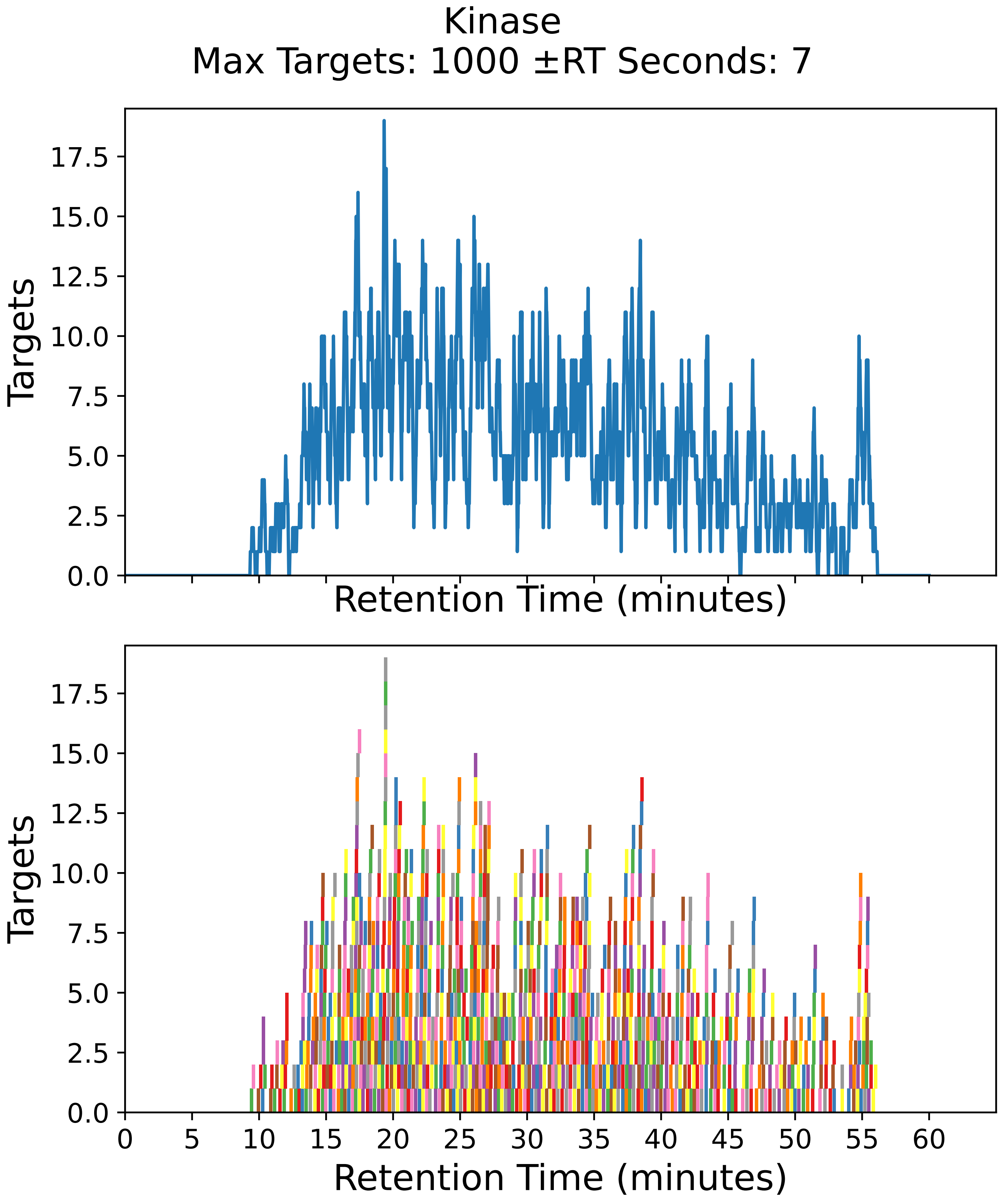

Supplementary Figure 67: Top: utilization diagram for the best fit method with instrument targeting the kinase filtered subset with a maximum of 1000 targets per cycle with retention times ± 7 seconds from the peptide peak apex. Bottom: individual targeting windows for the 1022 peptides, monitoring 511 proteins.

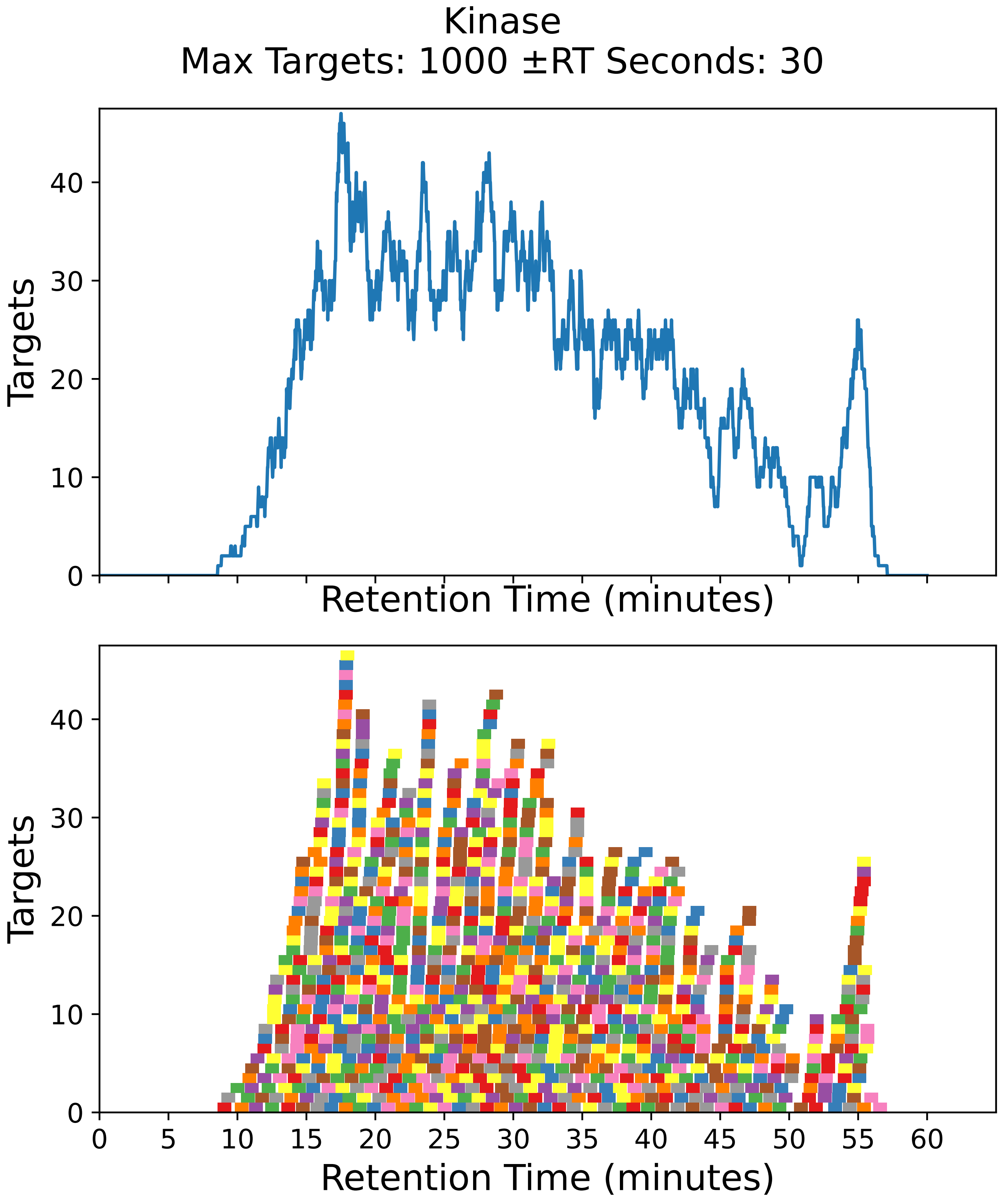

Supplementary Figure 68: Top: utilization diagram for the best fit method with instrument targeting the kinase filtered subset with a maximum of 1000 targets per cycle with retention times ± 30 seconds from the peptide peak apex. Bottom: individual targeting windows for the 1022 peptides, monitoring 511 proteins.

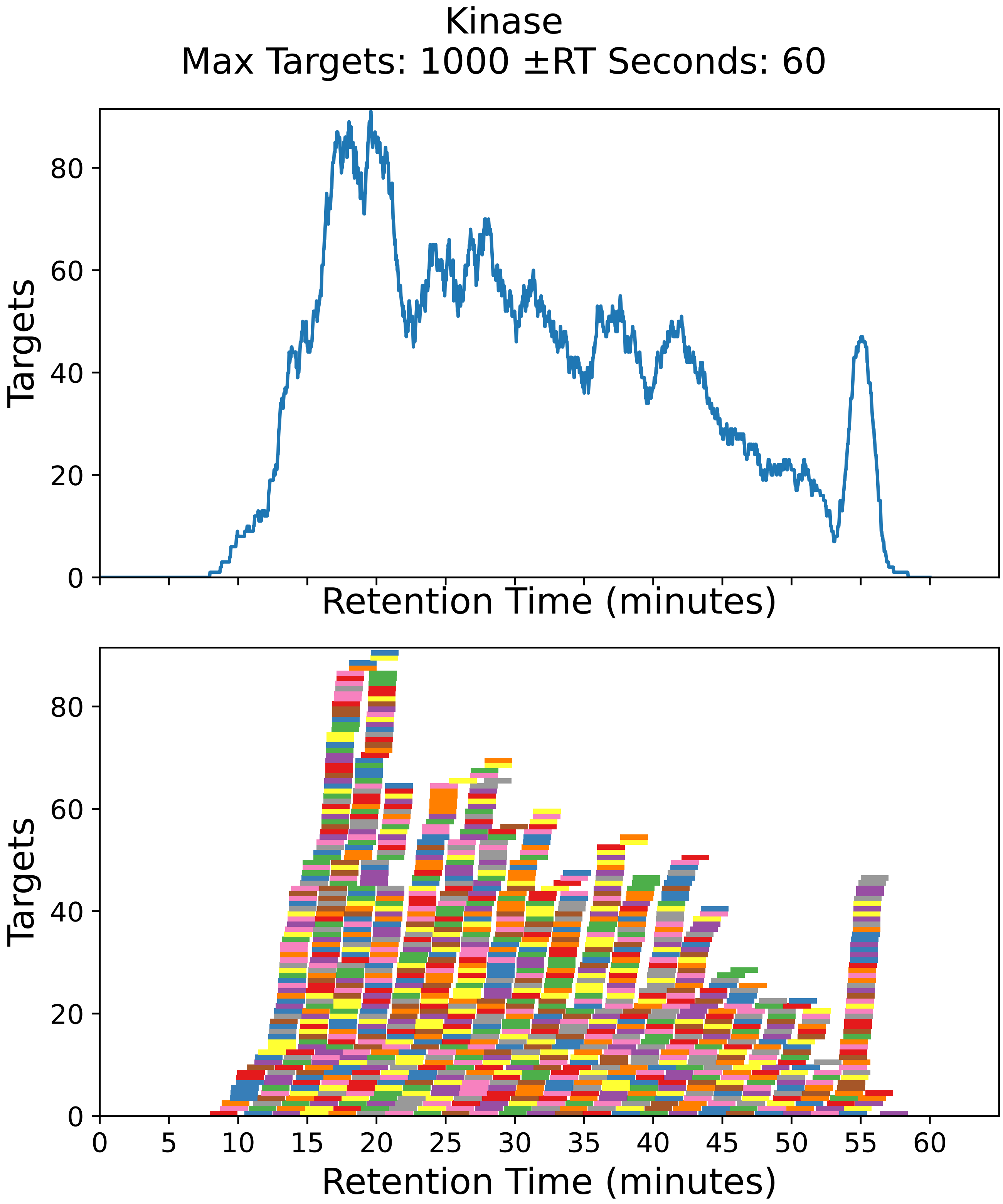

Supplementary Figure 69: Top: utilization diagram for the best fit method with instrument targeting the kinase filtered subset with a maximum of 1000 targets per cycle with retention times ± 60 seconds from the peptide peak apex. Bottom: individual targeting windows for the 1022 peptides, monitoring 511 proteins.

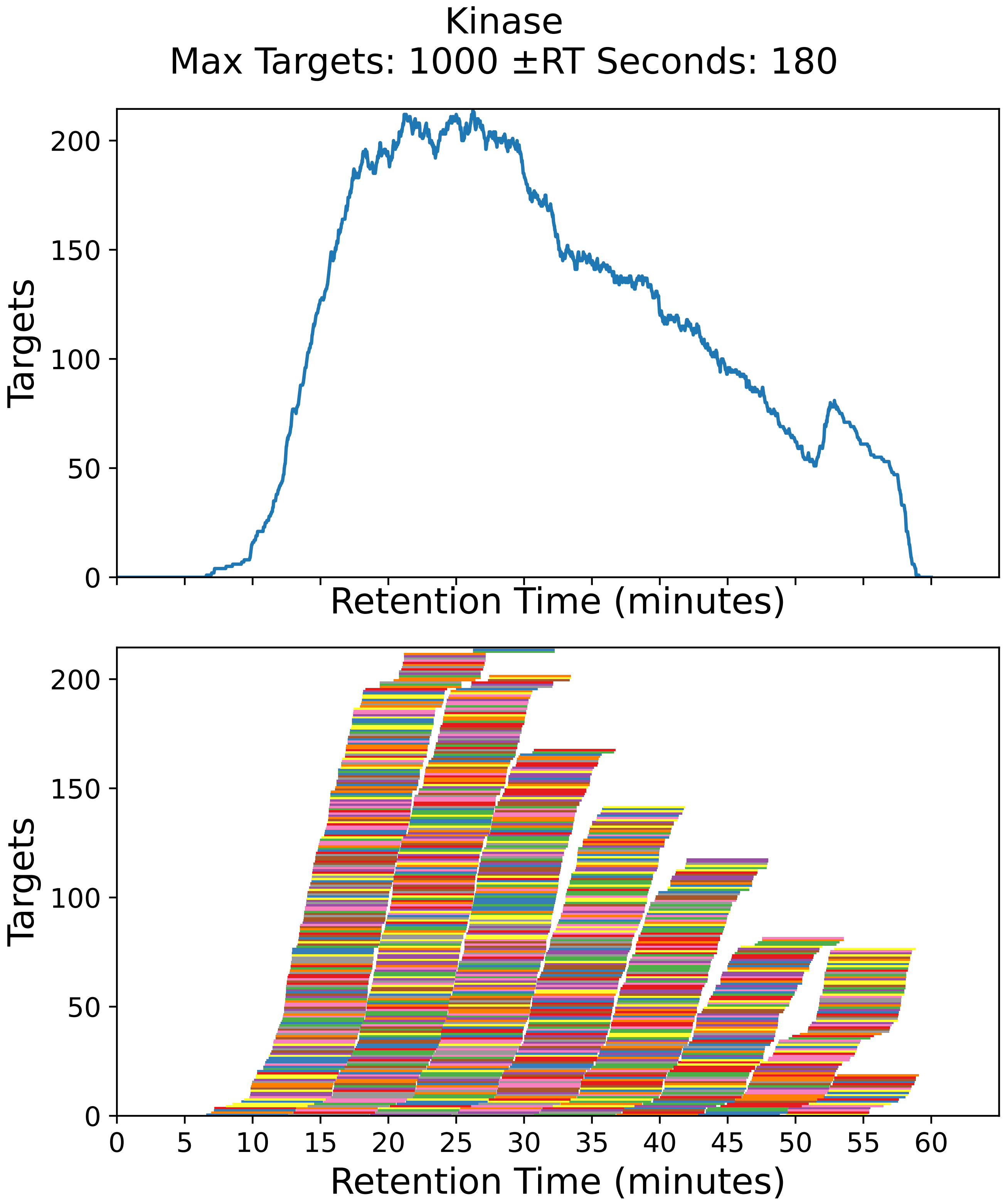

Supplementary Figure 70: Top: utilization diagram for the best fit method with instrument targeting the kinase filtered subset with a maximum of 1000 targets per cycle with retention times ± 180 seconds from the peptide peak apex. Bottom: individual targeting windows for the 1022 peptides, monitoring 511 proteins.

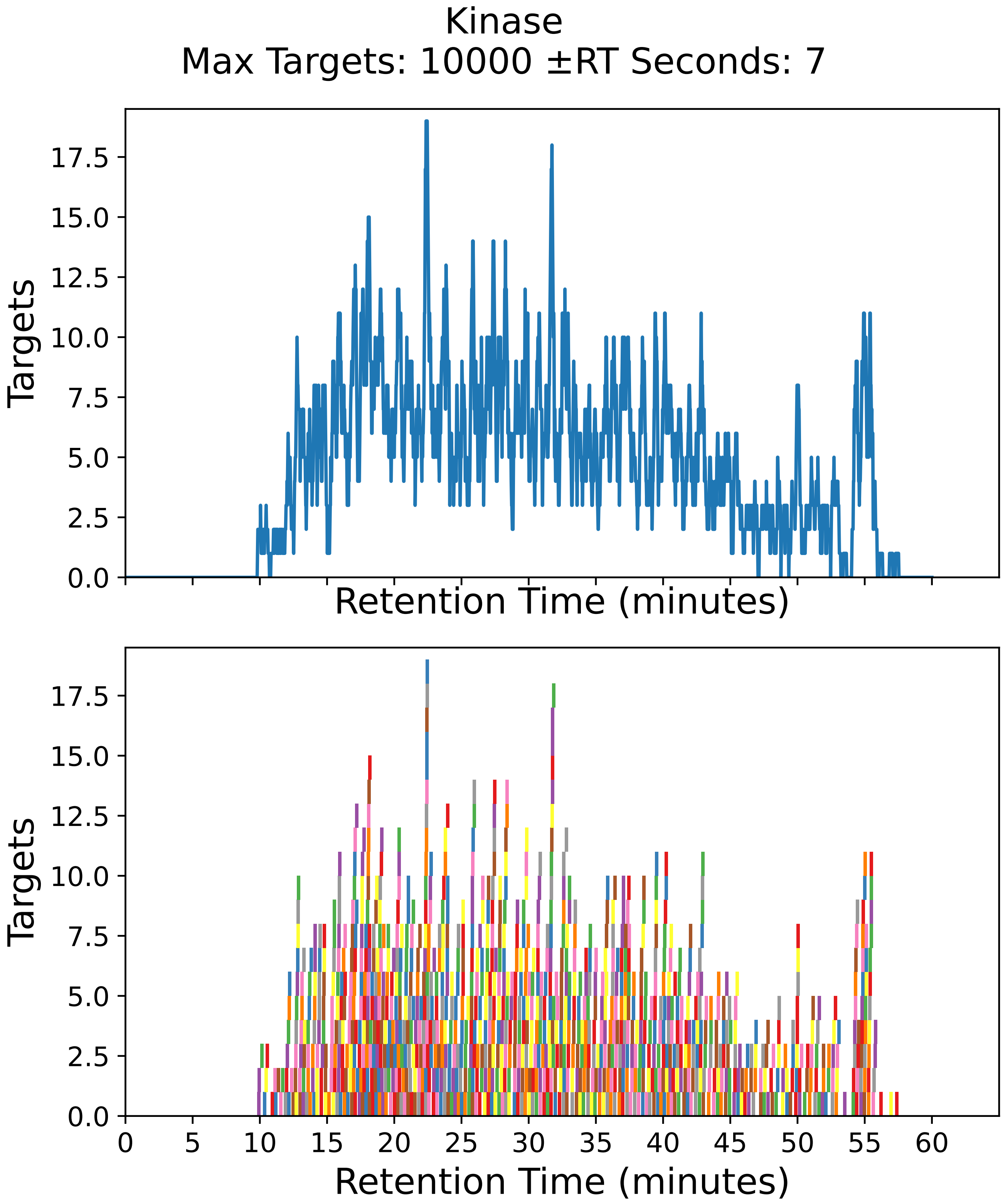

Supplementary Figure 71: Top: utilization diagram for the best fit method with instrument targeting the kinase filtered subset with a maximum of 10000 targets per cycle with retention times ± 7 seconds from the peptide peak apex. Bottom: individual targeting windows for the 1022 peptides, monitoring 511 proteins.

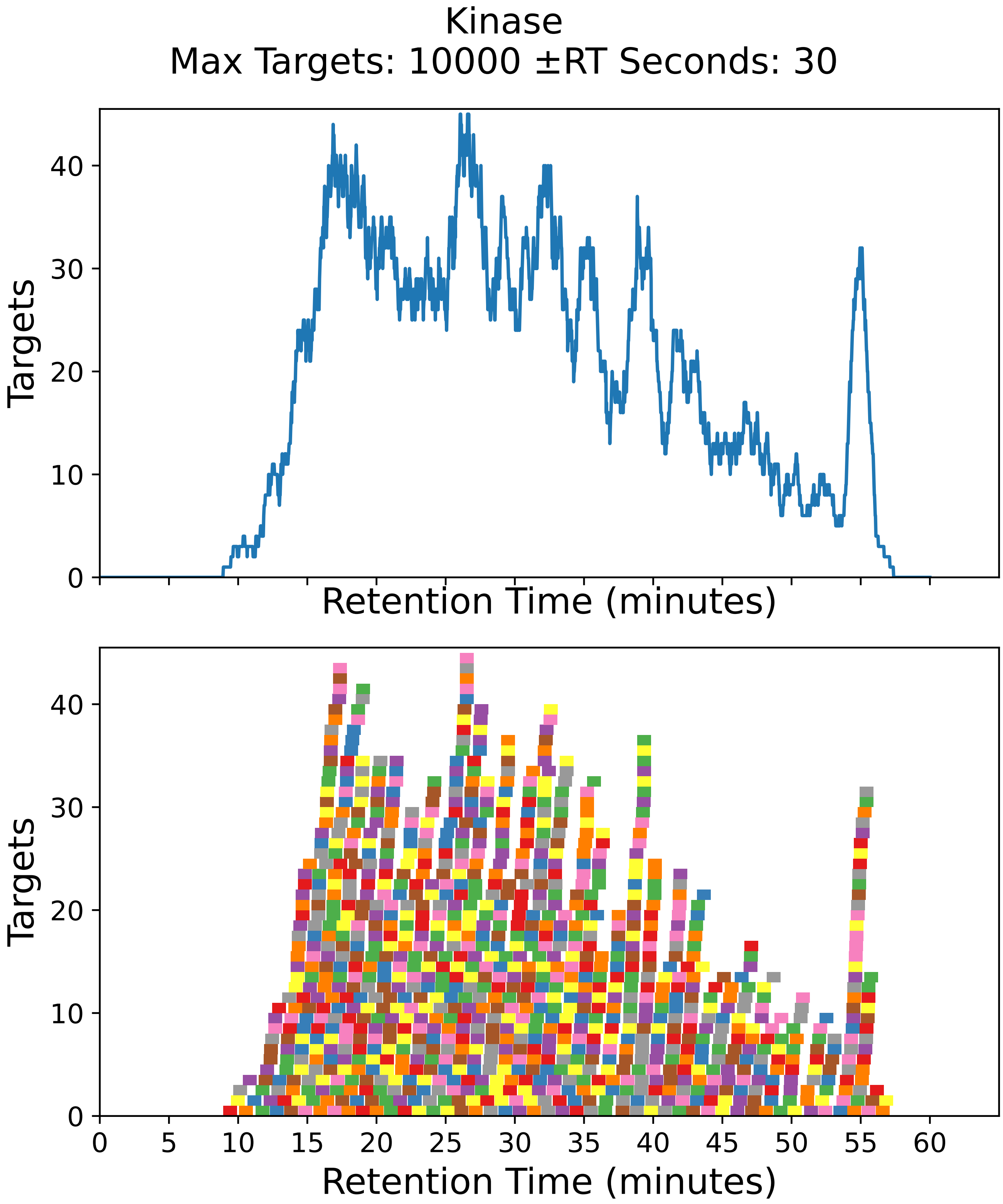

Supplementary Figure 72: Top: utilization diagram for the best fit method with instrument targeting the kinase filtered subset with a maximum of 10000 targets per cycle with retention times ± 30 seconds from the peptide peak apex. Bottom: individual targeting windows for the 1022 peptides, monitoring 511 proteins.

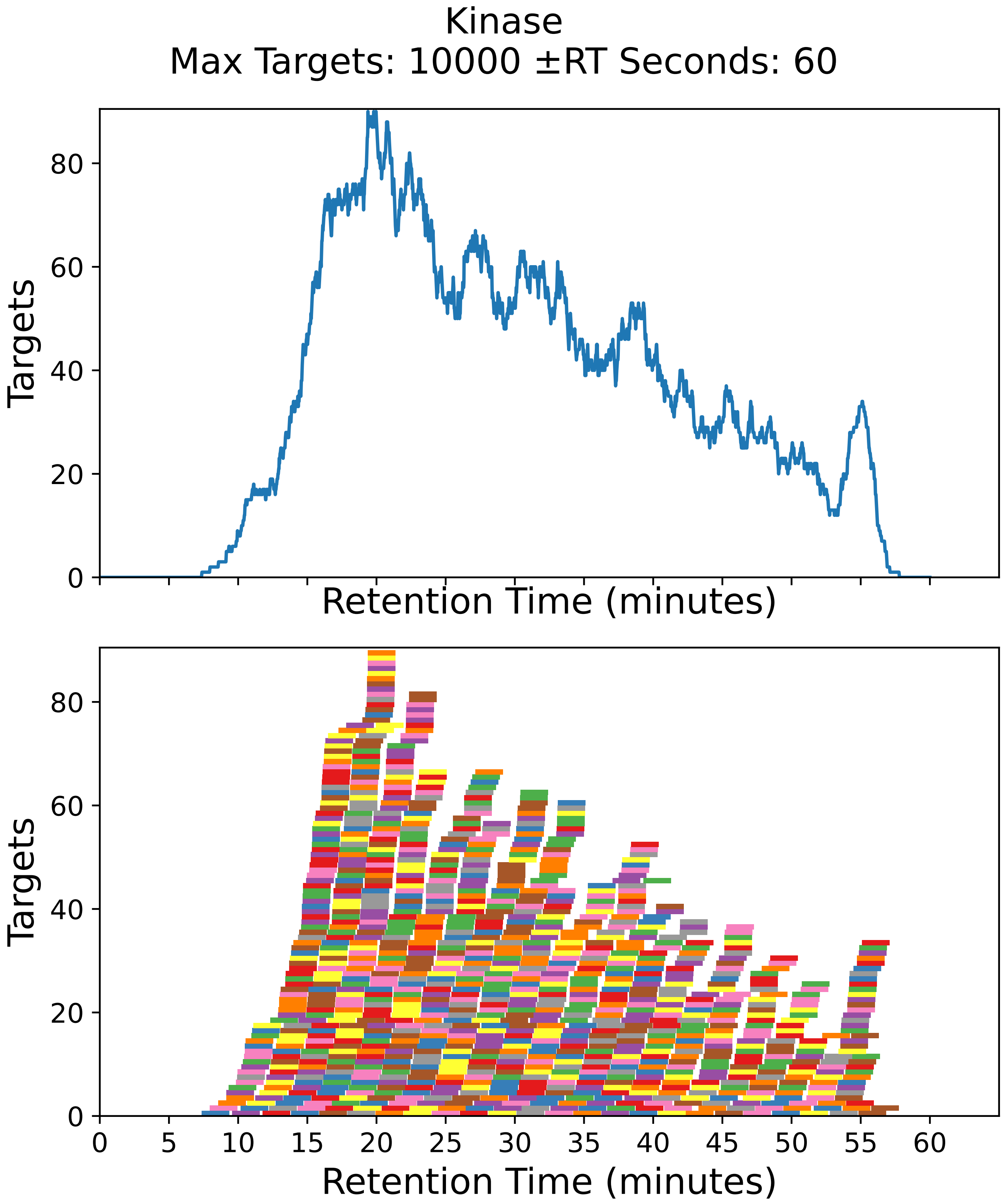

Supplementary Figure 73: Top: utilization diagram for the best fit method with instrument targeting the kinase filtered subset with a maximum of 10000 targets per cycle with retention times ± 60 seconds from the peptide peak apex. Bottom: individual targeting windows for the 1022 peptides, monitoring 511 proteins.

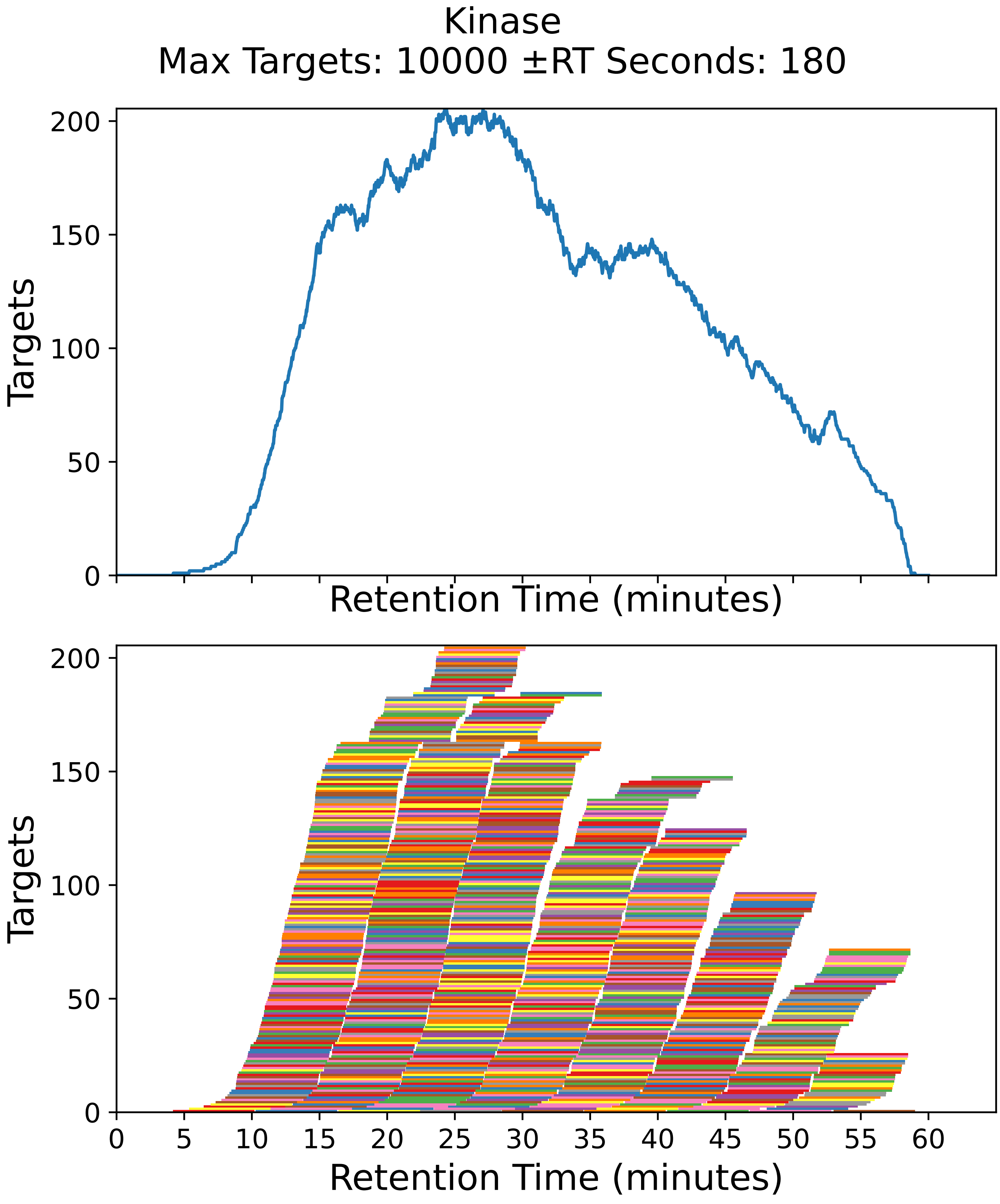

Supplementary Figure 74: Top: utilization diagram for the best fit method with instrument targeting the kinase filtered subset with a maximum of 10000 targets per cycle with retention times ± 180 seconds from the peptide peak apex. Bottom: individual targeting windows for the 1022 peptides, monitoring 511 proteins.
